# Fast pairwise coalescence enables gene-resolution scans for selection in diverse human populations

**DOI:** 10.64898/2026.05.21.726777

**Authors:** Kevin Korfmann, Sara Mathieson

**Affiliations:** Department of Biology, University of Pennsylvania, Philadelphia, PA, USA

**Keywords:** selective sweep, purifying selection, coalescence time, TMRCA, GPU, population genetics, gene-resolution coalescence gene list, pairwise coalescence, aDNA comparison

## Abstract

Pairwise coalescence times provide a way to compare genealogical variation across genomes and populations. We introduce `gamma_smc_cu`, a GPU implementation of gamma_smc with accuracy comparable to the reference method in simulations. It processes 63,190 haplotype pairs in 15.91 seconds, 67.7-fold faster than the CPU reference. Applied to the 1000 Genomes Project, we use `gamma_smc_cu` to produce gene lists conspicuous for their low coalescence time across 26 populations and a catalog of 143 genomic regions of interest. We quantified genomic covariates, such as recombination rates and the relation with haplotype statistics to evaluate the regions. We further benchmark extensively using independent simulations to compare with summary statistics such as iHS and XP-EHH. Case studies at *GRK2* and *TREML1* /*TREM2* illustrate these coalescence patterns and other regional evidence such as potential background selection or ancient genomic data derived tracks. We emphasise that positive and purifying selection can generate similar inferred profiles and provide a parameter search to show how both processes can lead to the same local signal. The software implemented here targets fast analysis capability of genealogical variation alongside a prioritization approach for regions of evolutionary significance.

## Introduction

Pairwise times to the most recent common ancestor (TMRCA) describe how genealogical branch depth varies along genome sequences. Approaches modeling the hidden correlation structure of sequences, such as `PSMC` and `MSMC`, infer coalescent histories from sequence variation [1–3]. `ASMC` enabled efficient inference of pairwise coalescence times along the genome, with applications to recent selection and deeper coalescent variation [4]. `gamma_smc` further accelerated this inference through a continuous approximation to the posterior distribution [5]. Making these estimates available across populations provides a common basis for comparing evolutionary patterns and relating them to genomic annotations.

Full pairwise inference remains computationally demanding, mainly due to its quadratic nature in the number of evaluable haplotype pairs for even modest sample sizes. The 3,202 individuals in the high-coverage 1000 Genomes cohort [6] come out to be 829,638 within-population haplotype pairs across 26 populations. Even an illustrative 1 kb grid across 3 Gb would contain approximately 2.5 *×* 10^12^ pair-by-position values. Each 1 kb interval can contain multiple segregating sites, which the inference evaluates individually. `gamma_smc` parameterizes the posterior TMRCA distribution as a Gamma distribution, avoiding time discretization [5]. One approach to make this fast can be identified as using parallel computation across pairs, which offers a route to population-scale inference. Another path would be through dating of ancestral recombination graphs, providing another route to these genealogical summaries [7–10], while methods such as cxt use local site-frequency information to infer TMRCA distributions [11, 12]. These approaches require ARG reconstruction or, for cxt, simulation-based training.

One use for pairwise TMRCA inference is the examination of genes where haplotype statistics are not available. For example, the integrated haplotype score (iHS) requires a central polymorphism with minor allele frequency (MAF) of at least 5%, and there could be a situation where the gene does not have any sites meeting this threshold [13]. Cross-population extended haplotype homozygosity (XP-EHH) provides a complementary comparison between populations, including near-fixed sweeps [14]. Our TMRCA summary includes rare variants and is computed for gene–population combinations with at least two segregating sites in the gene body. Here, we decide to focus on the gene level as a resolution target through the aggregation of per-site inferences to provide an annotation unit, which can then be compared with population or functional literature evidence. The detection power and localisation of a selected variant are separate properties, which we evaluate explicitly in simulations.

Also, of increasing importance are ancient-DNA scans, which provide further possibilities for comparison with traditional methods through analysing allele-frequency change through time. Akbari *et al.* [15] analysed a West-Eurasian dataset covering the last 18,000 years and identified 479 loci with evidence of selection. Time-series evidence from ancient DNA and modern-genome genealogical summaries differ in sampling, geographic coverage and the histories to which they are sensitive. However, comparing signals from both datasets can be useful, in particular when signals are shared throughout time-scales.

Here, we introduce `gamma_smc_cu`, a CUDA implementation of the `gamma_smc` forward–backward algorithm, and evaluate its accuracy and population-scale runtime against the reference implementation. The implementation functionally provides the same output as `gamma_smc`, which is pairwise inference of coalescence times per site and we use it to provide TMRCA values of genes across populations. Therefore, we apply it to 19,119 autosomal protein-coding genes across 26 populations, focussing in particular on regions with low coalescence times. These genome-wide analyses quantify dependence on gene length, site count, recombination and background selection (BGS), and compare gene-body and flanking iHS/XP-EHH values. We use simulations to benchmark the detection and localisation capability of our approach across four sweep scenarios, and we choose two main examples to show the scope of the resource. First, we focus on *GRK2*, a low-rank gene within a region with prior selection evidence, and *TREML1* /*TREM2*, a region with another interesting coalescence pattern. Finally, we use simulations to examine whether positive and purifying selection can generate similar inferred profiles. And generally, the software can also analyse other species with phased haplotype data.

## Results

### GPU-accelerated gamma_smc_cu

The GPU acceleration of `gamma_smc_cu` is achieved by parallelizing the computation of the `gamma_smc` forward–backward algorithm by running calculations on several haplotype pairs simultaneously via thread blocks (Methods). Across 14 demographic configurations from eight species in stdpopsim (*Homo sapiens*, *Pan troglodytes*, *Pongo pygmaeus*, *Canis familiaris*, *Bos taurus*, *Arabidopsis thaliana*, *Drosophila melanogaster* and *Anopheles gambiae*), `gamma_smc_cu` and the reference `gamma_smc` yield high correlations between inferred and true log-TMRCA from msprime [16] (Fig. 1b). In this benchmark, the median of the simulation-specific median Pearson correlations is 0.833 for `gamma_smc_cu` and 0.832 for the reference, with some ranging differences from *−*0.002 to +0.031. Median natural-log TMRCA RMSE is 0.657 and 0.655, respectively; and simulation-specific medians range from RMSE roughly 0.5 to 1(Fig. 1c). Direct correlations between the two methods have a median of 0.994 across all simulation parameters and are thus essentially interchangeable.

For YRI chromosome 22, the 356 haplotypes give 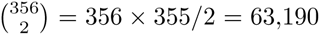 distinct pairs, counting each pair once. We processed all pairs across 313,277 sites in three timed repetitions after warm-up. We measured a total elapsed time of 15.91 s (range 15.19–18.21 s) for `gamma_smc_cu` on one NVIDIA V100S. CPU/ASMC native total times on the same machine and panel were 1,077 s (17.9 min) for the CPU reference, 7,103 s (118.4 min) for single-worker ASMC and 996 s (16.6 min) for eight-worker ASMC [4] on an Intel Xeon Gold 6148 (Fig. 1d,e). The CUDA workflow is thus about 67.7-fold faster than the CPU reference and 62.6-fold faster than ASMC with eight-workers.

**Figure 1:**
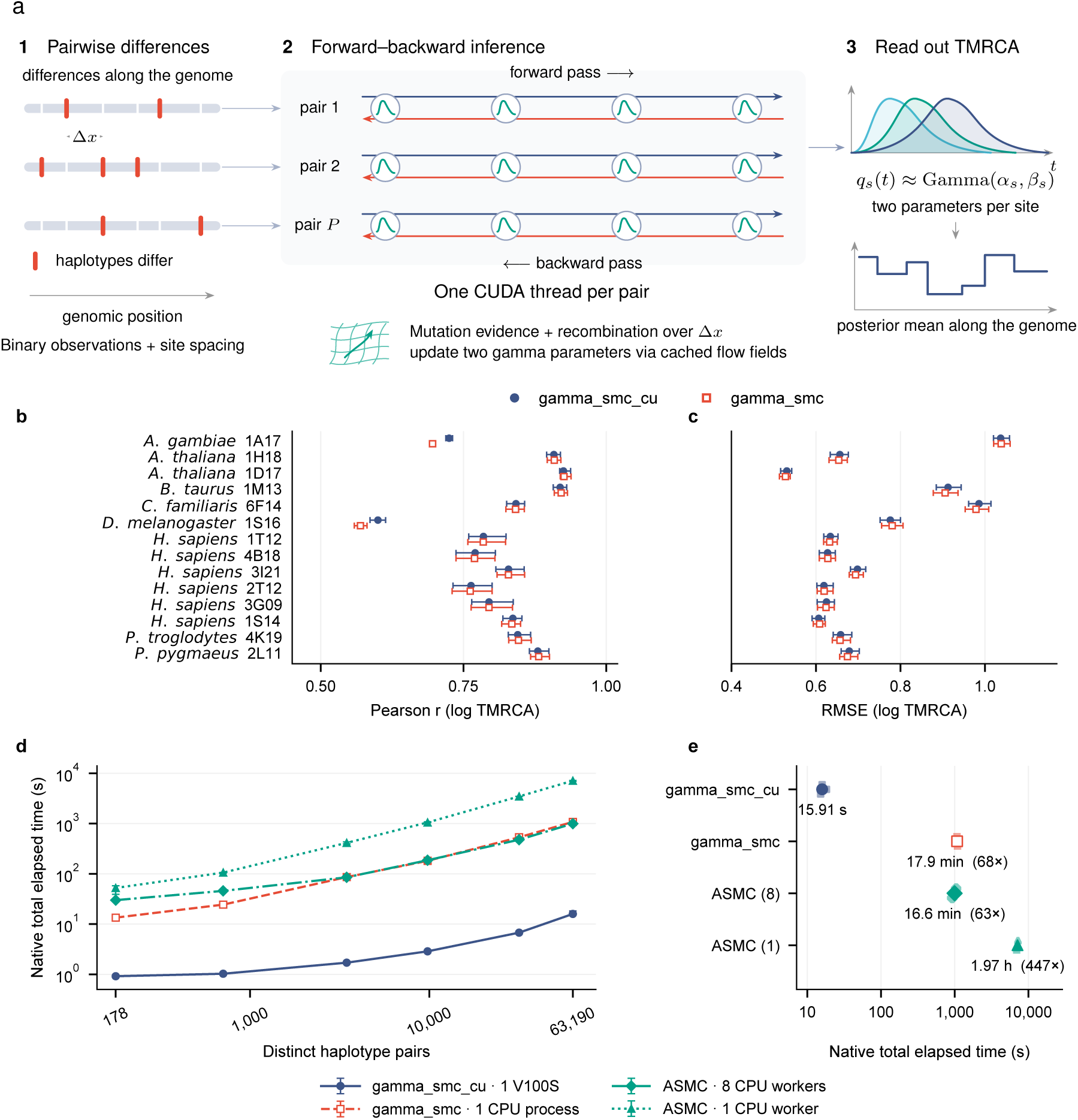
gamma_smc_cu: algorithm, accuracy and runtime. (a) Each CUDA thread processes one haplotype pair in both directions, updating a gamma distribution using pairwise differences and distances between consecutive sites (Δ*x*, in base pairs). (b,c) We compare inferred and true log-TMRCA using correlation and RMSE across 14 stdpopsim configurations from eight species. We simulated one 5 Mb sequence with msprime and analysed 190 haplotype pairs per configuration. Points and whiskers show medians and interquartile ranges across pairs. (d,e) We measured runtimes for YRI chromosome 22 of 356 haplotypes with a total of 313,277 sites on an NVIDIA V100S GPU and Intel Xeon Gold 6148 CPU. Large points show medians and small points show individual runs.

### Genome-wide application to the 1000 Genomes Project

Our analyses were conducted with `gamma_smc_cu` on 3,202 individuals (6,404 haplotypes) from the 1000 Genomes project [6]. In this analysis we considered all 22 autosomes and computed TMRCAs for all 829,638 within-population pairs of haplotypes across 26 different populations. These populations belong to five continental groups: AFR, EUR, EAS, SAS, AMR, with sample sizes ranging from 74 to 179 diploid individuals (10,878 to 63,903 haplotype pairs per population). Our inference pipeline operates pair-wise on the GPU, decoding the sites of each chromosome as one inference block, in chunks of 512 pairs. This preserves full-chromosome forward–backward context. GPU memory scales with chromosome length and pair-chunk size. We averaged the per-site (SNP) posterior TMRCAs within each of the 19,119 autosomal GENCODE v46 protein-coding genes in our pipeline [17]. This was accomplished by taking the geometric mean across segregating sites within each gene for each pair of haplotypes, then the geometric mean was taken again across all within-population haplotype-pair estimates (see Methods). For each population independently, genes were ranked by TMRCA relative to the population’s genome-wide distribution. Hence, a lower TMRCA percentile identifies a gene with more recent inferred coalescence relative to the population’s genome-wide distribution. Importantly, this aggregation does not automatically lead to the detection of positive selection, since alternative explanations such as purifying selection can also lead to a reduction in diversity and, by extension, a reduction in coalescence times, which we investigate in the later parts of the study.

The Manhattan plot in Fig. 2a displays continuous gene ranks. Genes with a rank in the lowest-1% define an initial screen of the strongest signals, however genes outside this cut-off may very-well still be of interest. We also apply further cutoffs of 0.5%, 2% and 5% uniformly to the entire gene set while holding other filters fixed (see Supplementary Table S13).

### Masking of segmental duplications

Among 19,119 genes, we selected 1,123 (5.9%) that overlap by at least 50% with the genomic super duplications from the genomicSuperDups UCSC table [18], because short read mapping in duplicated regions biases the heterozygosity calculation and consequently coalescence time estimates [19, 20]. Table S5 shows the comparison between masked and unmasked top 50 results.

### Previously identified selected loci

We examined within-population TMRCA ranks of a limited set of 23 reference loci previously investigated in selection scans (e.g., [13, 14, 21, 22]) or functional and trait-association studies (locus-specific references in Table 1). Across the 26 populations, 19 loci had a minimum within-population rank below 10%, 16 below 5%, and eight below 1%. Some examples include *SLC24A5* (0.04% in TSI) [23], *LCT* (0.29% in CEU) [24], and *EDAR* (0.46% in CHB) [25]. These results show the utility of within-population TMRCA ranks for highlighting previously studied loci.

Specifically, four of the 23 reference loci had minimum within-population TMRCA ranks above 10% in our analysis, showing that previously implicated loci need not show low TMRCA ranks in the sampled populations. Further, *HBB* represents an example of balancing selection, and *MC1R* is reported to be under population-specific constraint. We also note that *OCA2* does not fall within the lowest 10% of TMRCA ranks in any sampled population, while the Tibetan population in which *EPAS1* is associated with high-altitude adaptation is not represented in the 1000 Genomes dataset analysed here. The classical HLA genes instead have known deep coalescence times, ranking above the 99th percentile across all 26 populations. Notably, in the European *ALDH2* /*SH2B3* region, linkage can produce low gene-body TMRCA across neighbouring genes without resolving which gene is the target of selection. We provide more detailed locus interpretations and supporting citations in the Supplementary part 3.7.

We also compared modern-population coalescent depths at 474 variants derived from the summary statistics of Akbari *et al.* [15]. These variants are separated by at least 340 kb within chromosomes and have reported posterior probabilities of selection of at least 0.988, with all but five meeting the 0.99 threshold.

The pairwise TMRCA of each Akbari variant was summarised over a *±*25 kb window using full-chromosome inference across all 26 1000 Genomes populations, yielding a window-level coalescent-depth summary at each aDNA location (Fig. 2b,c). Interestingly, a large set of variants identified in the ancient data still have striking population specific patterns, as primarily shown by the left panel of the heatmap, for example the variant near *SLC24A5* has shallower coalescence in European than in African or East Asian panels, whereas other aDNA peaks have much deeper coalescence.

**Figure 2:**
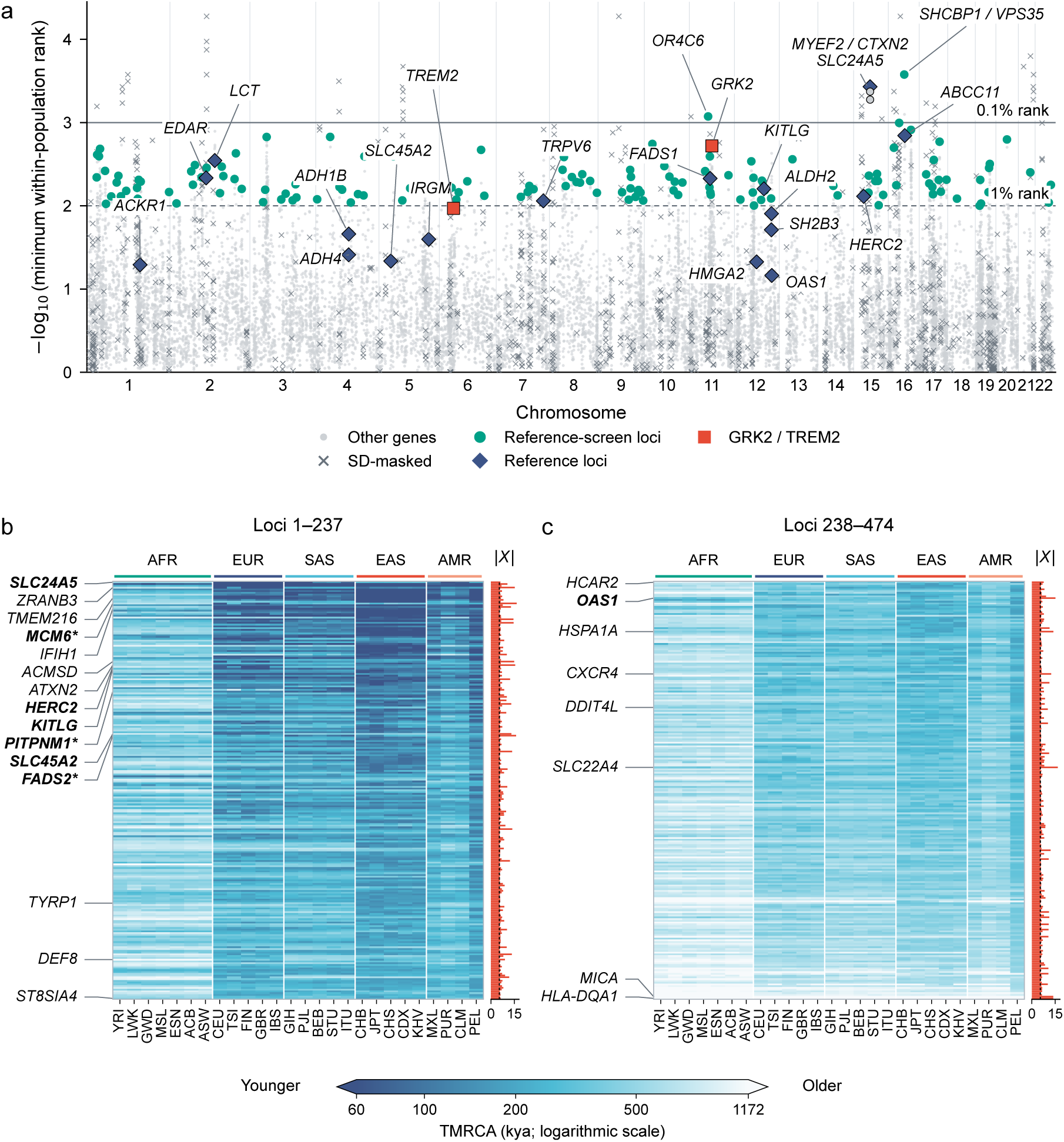
Genome-wide coalescent ranks and highlighted ancient-DNA regions. (a) Per gene *−* log_10_ minimum within-population rank across 26 populations (18,982 genes). Symbols identify other genes, segmental-duplication exclusions, 143 representative genes from this study, reference loci and the two main case studies. Lines mark 1% and 0.1% ranks. (b– c) TMRCA in *±*25 kb GRCh38 windows at 474 variants from Akbari *et al.* [15]. Rows continue between panels in order of minimum TMRCA across populations. Darker colours indicate younger coalescence on a logarithmic scale, which were clipped at the 2nd and 98th percentiles (60–1172 kya; 28 years per generation). Side bars show *|X|* and dashed lines mark 5.45. Bold heatmap labels identify the same genes as in panel a or nearby genes marked with an asterisk. The starred labels *PITPNM1* *, *MCM6* * and *FADS2* * refer to genes near *GRK2*, *LCT* and *FADS1*, respectively.

### Candidate identification

We decided on using five step filtering approach to identify and group candidate genes (Methods). First, we excluded genes overlapping segmental duplications to reduce mapping errors, leaving 17,996 genes. Second, we selected 620 genes with TMRCA ranks below 1% in at least one population to focus on unusually recent coalescence. Third, we required ranks below 5% in every population of at least one continental group to focus on shared signals, leaving 579 genes. Fourth, we excluded genes within 500 kb of the 23 reference loci to focus on regions beyond those already considered. The remaining 540 genes are referred to as stage-4 genes. Finally, we grouped nearby candidates using a 1 Mb clustering threshold, giving 143 loci: 82 multi-gene clusters and 61 clusters containing a single candidate gene (Supplementary Tables S2 and S3).

**Table 1:** TMRCA ranks at 23 reference loci. We ranked genes within each population using geometric-mean TMRCA from `gamma_smc_cu`. Lower percentiles indicate more recent coalescence. The rank range is the difference between the highest and lowest available percentiles across 26 populations. We list genes by their lowest percentile.

| Gene | Chr | Min rank | Pop | Range | Biology | Ref. |
|---|---|---|---|---|---|---|
| *Minimum rank* < 1% |  |  |  |  |  |  |
| *SLC24A5* | 15 | 0.04% | TSI | 22.60% | Skin pigmentation | [23] |
| *ABCC11* | 16 | 0.14% | CHB | 29.10% | Earwax | [26] |
| *LCT* | 2 | 0.29% | CEU | 80.22% | Lactase persistence | [24] |
| *EDAR* | 2 | 0.46% | CHB | 67.07% | Hair thickness | [25] |
| *FADS1* | 11 | 0.47% | ESN | 32.56% | Long-chain PUFA | [27] |
| *KITLG* | 12 | 0.63% | MXL | 19.02% | Pigmentation | [28] |
| *HERC2*<sup>\*</sup> | 15 | 0.77% | FIN | 74.69% | Eye color | [29] |
| *TRPV6* | 7 | 0.87% | JPT | 84.80% | Calcium absorption | [30] |
| *Minimum rank* 1–5% |  |  |  |  |  |  |
| *ALDH2* | 12 | 1.24% | TSI | 29.24% | Alcohol metabolism | [31] |
| *CASP12* | 11 | 1.63% | TSI | 52.78% | Sepsis resistance | [32] |
| *SH2B3* | 12 | 1.95% | TSI | 30.07% | Immune/celiac | [33] |
| *ADH1B* | 4 | 2.18% | CHS | 64.83% | Alcohol metabolism | [34] |
| *IRGM* | 5 | 2.52% | FIN | 78.45% | Crohn’s/autophagy | [35] |
| *ADH4*<sup>†</sup> | 4 | 3.88% | CHS | 79.20% | Alcohol metabolism | [34] |
| *SLC45A2* | 5 | 4.61% | TSI | 83.81% | Skin pigmentation | [36] |
| *HMGA2* | 12 | 4.72% | PEL | 26.39% | Height/growth | [37] |
| *Minimum rank* 5–10% |  |  |  |  |  |  |
| *ACKR1* | 1 | 5.11% | ESN | 86.97% | Malaria (Duffy) | [38] |
| *OAS1* | 12 | 6.87% | GWD | 73.16% | Antiviral immunity | [39] |
| *TYRP1* | 9 | 8.49% | KHV | 44.91% | Pigmentation | [13] |
| *Minimum rank* ≥ 10% |  |  |  |  |  |  |
| *OCA2* | 15 | 24.80% | JPT | 49.48% | Eye color | [40] |
| *MC1R* | 16 | 25.69% | JPT | 64.31% | Pigmentation | [41] |
| *EPAS1* | 2 | 50.94% | PEL | 28.38% | Tibetan altitude | [42] |
| *HBB* | 11 | 82.69% | PEL | 16.27% | Sickle / balancing | [43] |

<sup>\*</sup> Gene flagged by the unique-base UCSC genomicSuperDups overlap mask (Methods).
<sup>†</sup> Linked ADH-cluster comparison; Li *et al.* [34] studied *ADH1B*, not an independent *ADH4* sweep.

We present the ranks and 1% threshold along with a sensitivity analysis in the SI. Also, cutoffs of 0.5%, 1%, 2% and 5%, with all other filters fixed, keeps 58, 143, 233 and 277 loci, respectively. For each focal gene we provide rank counts per population after applying the rest of the filters (Supplementary Table S14).

We present seven case studies showing different coalescence patterns and how these relate to haplotype evidence and previous studies [15, 44, 45]. *GRK2*, *IFIH1* and *SLC6A15* are representatives in the 143-locus catalogue. The remaining examples fall outside the lowest-1% cutoff: *TREM2* /*TREML1*, *CCDC92* and *BPIFA2* have ranks below 2% in at least one population, whereas *CLEC6A* reaches only 3.30% and does not pass the continental replication filter even when the rank cutoff is relaxed to 5%. All seven case studies were selected post hoc (Methods).

The population patterns vary among these loci, for example *GRK2* falls within the lowest 1% in nine of the ten South Asian and European populations, with the remaining population just outside this cutoff. *IFIH1* falls below 1% in all ten, as does *SLC6A15* across all five East Asian populations. The lowest ranks for *CCDC92*, *CLEC6A* and *BPIFA2* occur in CDX, KHV and GIH, respectively. We use Galwey’s effective population counts to account for this dependence in a descriptive score summarising consistency across populations [46] (Methods).

**Table 2:** Seven case studies selected for detailed follow-up. We selected these cases after examining the results. The rank gives the TMRCA percentile in the focal population, and its continental group defines which populations we use as a comparison. Only *GRK2*, *IFIH1* and *SLC6A15* belong to the 1% catalogue. Depl:Enr divides the number of sites where the ALT allele is less common in the focal population by the number where it is more common, using the pooled populations outside its continental group for comparison. We calculated this ratio and maximum *F*_ST_ from genotypes within *±*500 kb of each gene midpoint.

| Gene | Chr | Rank | Pop | Continental Group | Depl:Enr | Max *F*<sub>ST</sub> | Biology |
|---|---|---|---|---|---|---|---|
| *GRK2 and TREM2* |  |  |  |  |  |  |  |
| ***GRK2*** | 11 | 0.19% | GIH | SAS+EUR | 7.1:1 | 0.43 | *β*-adrenergic kinase |
| ***TREM2*** | 6 | 1.26% | IBS | all non-AFR | 5.3:1 | 0.29 | Microglial innate immunity |
| *Additional case studies* |  |  |  |  |  |  |  |
| *IFIH1* | 2 | 0.44% | IBS | EUR | 8.5:1 | 0.51 | MDA5 dsRNA sensor (prior pos. ctrl) |
| *CCDC92* | 12 | 1.85% | CDX | EAS | 6.5:1 | 0.53 | WHR/metabolic GWAS locus |
| *CLEC6A* | 12 | 3.32% | CDX | EAS | 5.9:1 | 0.46 | Dectin-2, fungal immunity |
| *SLC6A15* | 12 | 0.54% | CHS | EAS | 5.2:1 | 0.60 | Brain amino-acid transporter |
| *BPIFA2* | 20 | 1.28% | GIH | SAS | 6.6:1 | 0.29 | Salivary antimicrobial protein |

### Gene length, number of polymorphic sites, recombination rate and BGS

We studied correlations of TMRCA ranks with gene length, the number of polymorphic sites, recombination rate and background selection (BGS). Each population provided around 14,200 genes with complete information, after excluding genes that overlapped with segmental duplications. We used *B^′^*, the estimated fraction of neutral genetic diversity remaining under BGS, to measure BGS strength. Lower values indicate stronger BGS. TMRCA ranks were correlated most strongly with *B^′^* and recombination rate across the 26 populations, with a smaller correlation with site number and a very weak correlation with gene length (Supplementary Fig. S6a,b and Supplementary Table S8). After adjustment for the other genomic parameters, the correlations with recombination rate and *B^′^* stayed positive, whereas the correlation with gene length became negative. These results show a relationship between genomic context and TMRCA ranks, including the association between the number of polymorphic sites and coalescence times. Next, we matched genes in the bottom 1% of TMRCA ranks with genes on other chromosomes that had similar values for all four genomic parameters. We also matched whether iHS could be calculated within the gene (Supplementary Fig. S6c,d). Among 2,008 matched gene–population cases with available iHS scores, low-TMRCA genes more often fell into the top 5% of haplotype scores than their controls. The difference was 10.9 percentage points for iHS (95% chromosome-jackknife interval 5.6–16.2) and 22.9 points for XP-EHH (12.9–32.9), both measured within gene bodies. Among 277 matched cases without iHS scores, the XP-EHH difference was 8.5 percentage points, with an interval spanning zero (*−*4.0 to 20.9).

For the focal genes, *GRK2* in GIH remained among the lowest 1% of TMRCA values relative to matched genes and had a high gene-body XP-EHH score. In contrast, the gene-body XP-EHH score for *TREM2* in IBS was near the middle of the distribution. Overall, low-TMRCA genes with iHS scores showed stronger support from haplotype data than their controls. For genes without iHS scores, the XP-EHH comparison was less conclusive.

### Prior literature and case studies

We compared our case-study loci with PopHumanScan [44] and classical selection scans [13, 14, 21, 22, 47], together with the more recent scans of Johnson & Voight [45] and Akbari *et al.* [15].

Our case studies combine TMRCA ranks with allele-frequency, iHS, nSL and *H*_12_ summaries (Tables 2 and 3). CLUES2 analyses use posterior marginal trees at focal variants to examine allele-frequency histories under the specified demographic and selection model (Methods). The five additional cases and *LCT* comparison are shown in Supplementary Fig. S8; the detailed literature comparison for *TREML1* /*TREM2* is in Supplementary Table S6. In total, we report 13 focal-variant estimates and their confidence intervals in Supplementary Table S7.

### Gene-level coalescence at *GRK2* and *TREML1* /*TREM2*

Our two main case studies are presented in detail below, which show how gene-level coalescence landscapes complement haplotype statistics alongside further allele-frequency analysis in the form of CLUES2 (Fig. 3). We present more detailed population summaries, haplotype analysis and literature summary see the Supplementary Sections 3.1 and 3.2.

**Figure 3:**
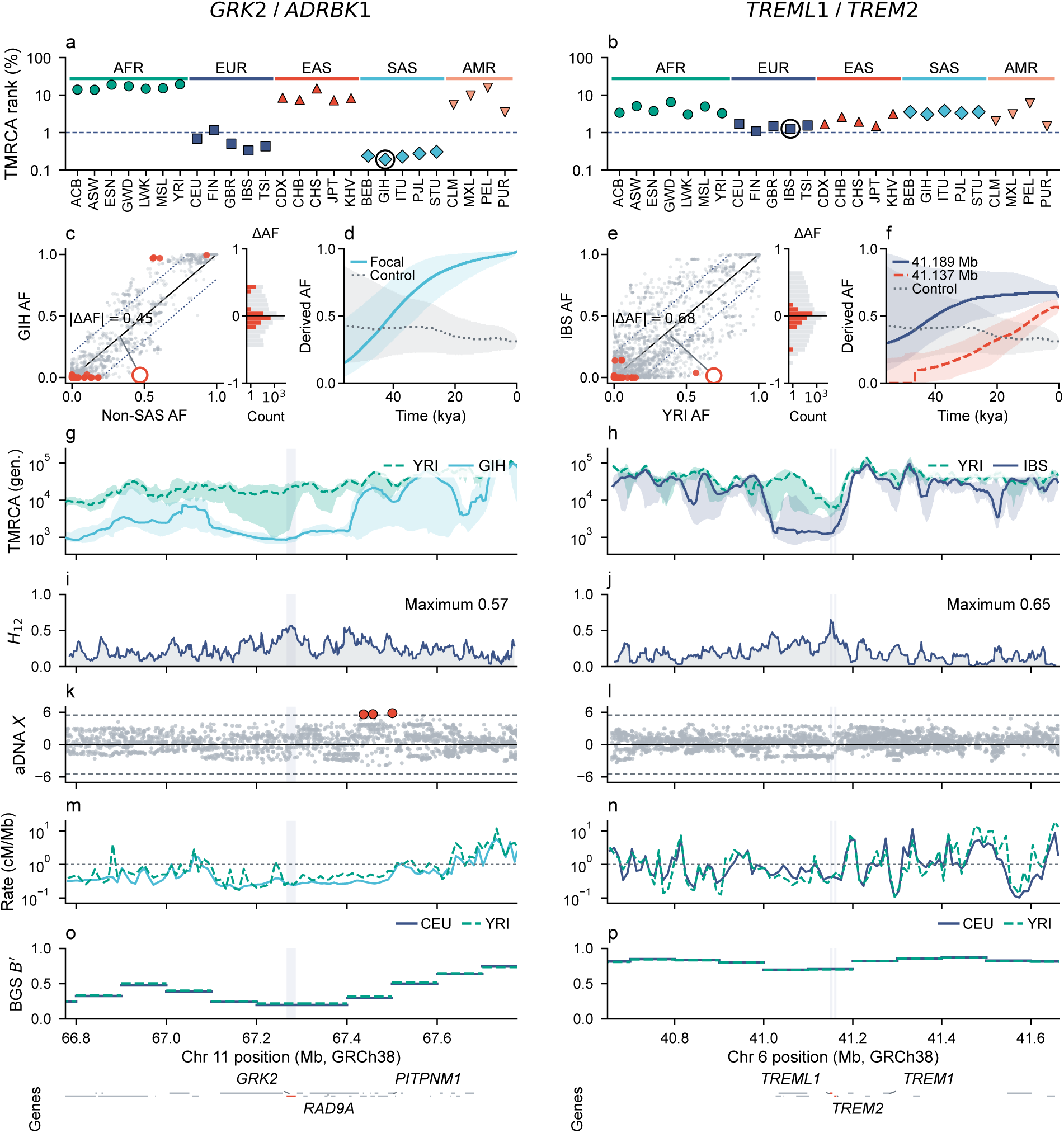
Regional evidence at *GRK2* (left) and *TREML1* /*TREM2* (right). (a,b) We rank TMRCA across populations. Circles mark GIH and IBS. Dashed lines mark the lowest 1%. (c,e) We compare ALT-allele frequencies between focal and comparison populations and plot their differences as histograms. Coral markers identify variants within the focal gene. (d,f) We infer allele-frequency trajectories with CLUES2 and show 95% credible intervals. (g,h) We smooth TMRCA over 150 sites and show medians and interquartile ranges across 20 haplotype pairs. Solid lines show GIH or IBS, and dashed lines show YRI. (i,j) We calculate *H*_12_ in 400-variant windows, shifted by 50 variants. (k,l) Ancient DNA statistic *X* [15] with thresholds selected at *±*5.45 is presented. (m,n) We infer recombination rates with fastrho, average them in 10-kb windows and scale them using *Ne* = 10,000. (o,p) We show Buffalo–Kern predictions of diversity relative to the neutral expectation (*B^′^*) for CEU and YRI in 100-kb bins. Lower values are indicative of stronger background selection. Gaps mark excluded bins.

#### *GRK2*/*ADRBK1*: a strong SAS/EUR candidate

Previous haplotype scans reported evidence of selection at chr11q13.2 [13, 44, 45, 48]. Our analysis ranks *GRK2* below 1% in nine of the ten SAS and EUR populations, with FIN just outside this cutoff (range 0.19–1.16%; Fig. 3a). In GIH, *GRK2* has the shallowest TMRCA among the 13 genes examined in the region (Supplementary Fig. S3). Only 41 of the 400 variant positions examined in its 20 kb gene body are polymorphic in GIH, and none meet the MAF*≥* 5% iHS filter. Pairwise TMRCA therefore highlights recent coalescence where no gene-body variants meet the allele-frequency threshold for iHS (Fig. 3g). BEB has eligible variants and has a gene-body iHS rank of 0.44%.

An *H*_12_ peak of about 0.569 within the *GRK2* region (Fig. 3i) and an *F*_ST_ of 0.43 in the *±*500 kb neighbourhood provide further support for the regional signal (Table 3). Across this neighbourhood, 7.1 times as many sites have a lower ALT-allele frequency in GIH than in pooled non-SAS haplotypes as have a higher frequency. SAS and EUR carriers of the regional high-frequency allele also share similar haplotypes. Within 25 kb on either side of the marker, pairs of SAS and EUR carriers have only 1.20 times as many SNP differences as pairs of SAS carriers. SAS carriers and SAS noncarriers differ 8.88 times as much as pairs of SAS carriers (bootstrap intervals in Supplementary Section 3.1).

At this marker, chr11:67,407,126, 120 kb downstream of *GRK2*, the derived G allele is nearly fixed in GIH (98.1%), compared with 53% in non-SAS haplotypes. We also use CLUES2 [49, 50] to estimate a selection coefficient of *s*^ = 0.00519 (Fig. 3d, see Supplementary Table S7). Akbari *et al.* [15] report a significant ancient-DNA signal in a *PITPNM1* intron approximately 214 kb downstream of *GRK2*, whereas the *GRK2* gene-body maximum is *|X|* = 2.57 (Fig. 3k).

#### TREML1 /TREM2 : coalescence and haplotype analyses

At chr6p21.1, *TREM2* has a TMRCA rank below five percent in eighteen of the nineteen non-African populations studied. It ranks below two percent in ten of those populations, but never below one percent (range 1.07–5.95%, see Fig. 3b). The rankings for *TREML1* were also similar in these populations (1.56–5.67%). On the other hand, the rankings for African *TREM2* were between 3.05 and 6.50%. The regional pattern in IBS includes shallow coalescence combined with a peak value of *H*_12_ at about 0.65 located inside *TREML1* (Fig. 3h,j). The maximum value of *H*_12_ obtained for each of the seven non-African populations ranged from 0.58 to 0.73. This is in contrast to the maximum value of *H*_12_ obtained for YRI, which was 0.18. However, there are differences among populations regarding the exact location where the maximum value of *H*_12_ is reached.

The iHS statistic was not computable in the gene body for *TREM2* in any EUR or EAS population. We were able to estimate gene-body iHS for *TREML1*, but this did not provide strong evidence of a selective sweep in these populations. Conversely, in BEB, ITU, and STU, we identified five *TREML1* /*TREM2* gene– population hits based on gene-level iHS statistics that were among the top 0.4%. The classical candidate lists we examined did not include either gene, but a previously reported YRI Flex-Sweep interval annotated *TREML1* and partially overlapped *TREM2* [51]. More details are described in Supplementary Table S6.

Restricting the Akbari ancient-DNA data [15] to variants labelled PASS in the published summary statistics and located within the *TREML1* –*TREM2* interval, we find no significant signal: the maximum *|X|* is 2.10 and the maximum posterior probability is 0.070 (Fig. 3l).

CLUES2 estimates derived-allele selection coefficients between *−*0.00185 and 0.00361 at seven regional markers in IBS. Only one approximate 95% interval lies entirely above zero – the other six include zero (Fig. 3f, Supplementary Table S7). The common variation examined here is distinct from the rare *TREM2* R47H Alzheimer’s disease risk allele [52, 53].

At both loci, we compare regional TMRCA estimates with ASMC [4] and cxt [11] to check the patterns across inference methods (Supplementary Fig. S2). We further included recombination estimates from fastrho and BGS tracks for CEU/YRI to give a more general genomic perspective to the observed low coalescence (Fig. 3m–p). Together, these analyses localise low coalescence within both regions and connect it to population-specific haplotype and allele-frequency patterns.

**Table 3:** TMRCA, haplotype and allele-frequency summaries at five selected loci and seven more case studies. “TMRCA rank” shows where each gene ranks within its population. “iHS/nSL frac rank” ranks genes by the fraction of sites with absolute normalised scores above 2. The bold values mark ranks below 0.05. For sites within *±*500 kb of each gene midpoint, we compare ALT-allele frequencies between the focal population and pooled populations outside its continental group. “Dep:enr ratio” divides the number of sites where the ALT allele is less common in the focal population by the number where it is more common. “Max *F*_ST_” gives the highest Hudson *F*_ST_ across these sites. “Top var AF diff” gives the largest allele-frequency difference, keeping its sign. Positive values indicate a higher frequency in the focal population and negative values a lower frequency. “Undef.” means we could not calculate a score for the gene.

| **Gene** | **Pop** | **TMRCA** | **iHS** | **nSL** | **dep:enr** | **max** | **top var** |
|---|---|---|---|---|---|---|---|
|  |  | **rank** | **frac rank** | **frac rank** | **ratio** | ***F***<sub>**ST**</sub> | **AF diff** |
| *Literature reference loci* |  |  |  |  |  |  |  |
| *LCT* | CEU | 0.0029 | **0.0015** | **0.0045** | 12.6 | 0.64 | +0.68 |
| *SLC24A5* | GBR | 0.0006 | undef. | undef. | 10.2 | 0.71 | +0.71 |
| *ABCC11* | CHB | 0.0014 | 0.726 | 0.740 | 10.4 | 0.79 | −0.80 |
| *EDAR* | CHB | 0.0046 | **0.0027** | **0.0038** | 6.4 | 0.84 | +0.86 |
| *KITLG* | MXL | 0.0063 | 0.134 | 0.161 | 9.0 | 0.30 | +0.38 |
| *GRK2 and TREM2* |  |  |  |  |  |  |  |
| ***GRK2*** | GIH | 0.0019 | undef. | undef. | 7.1 | 0.43 | −0.45 |
| *GRK2* | BEB | 0.0024 | **0.0044** | **0.0118** | – | – | – |
| ***TREM2*** | IBS | 0.0126 | undef. | undef. | 5.3 | 0.29 | −0.40 |
| *Additional examples* |  |  |  |  |  |  |  |
| *IFIH1* | IBS | 0.0044 | 0.743 | 0.061 | 8.5 | 0.51 | +0.59 |
| *CCDC92* | CDX | 0.0185 | **0.0056** | **0.0066** | 6.5 | 0.53 | +0.58 |
| *CLEC6A* | CDX | 0.0332 | **0.0275** | **0.0334** | 5.9 | 0.46 | −0.49 |
| *SLC6A15* | CHS | 0.0054 | **0.0305** | 0.732 | 5.2 | 0.60 | −0.61 |
| *BPIFA2* | GIH | 0.0128 | 0.741 | 0.748 | 6.6 | 0.29 | −0.36 |

### TMRCA similarity under positive and purifying selection

To examine whether the shallow profiles are specific to positive selection, we compared simple models fitted to the genomic TMRCA profiles at *GRK2* in GIH and *TREM2* in IBS (Fig. 4). We fitted sweeps and recurrent purifying (negative) selection separately, using the empirical observation as the target and available recombination maps. Weighted log errors for the sweeps were 0.905 and 0.626 on 24 simulation runs, compared to 1.211 and 0.899 for the closest neutral control we tested. Although we did not reproduce all of the local features, the coalescence valleys were replicated well. Search trajectories and obtained parameters are shown in Supplementary Fig. S5.

**Figure 4:**
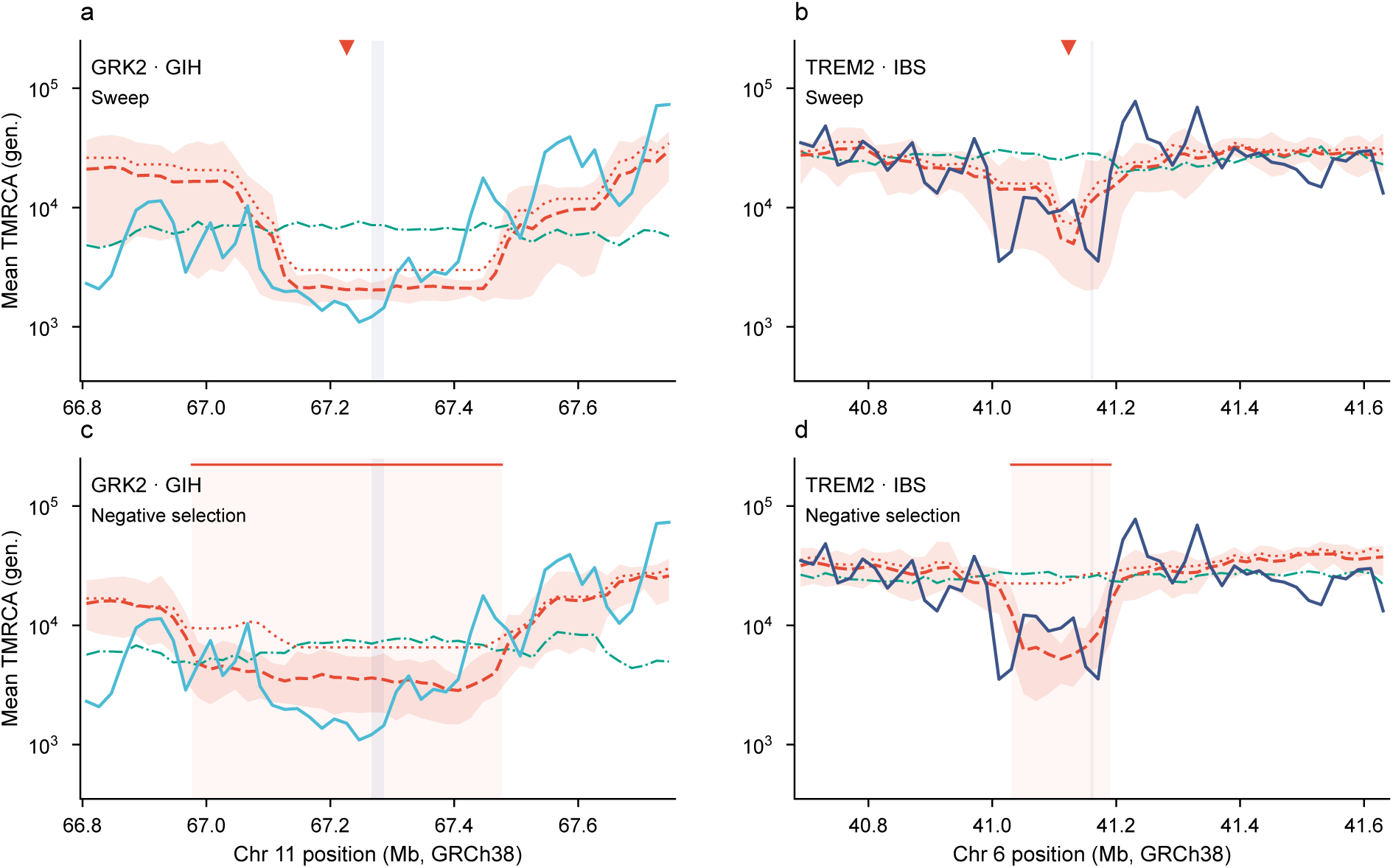
Similar inferred TMRCA profiles under positive and purifying selection. (a,b) Conditioned hard sweeps; (c,d) recurrent purifying (negative) selection. The plots on the left are for *GRK2* from GIH; while those on the right are for *TREM2* from IBS. The cyan-navy colored solid lines are empirical estimates of `gamma_smc_cu`; the coral colored dashed lines are `gamma_smc_cu` inferred model; the coral colored dotted lines are the TMRCA of the true genealogical history of the gene; and the teal dash-dotted lines are the optimal neutral control that have been tested. The coral colored regions denote 10th-90th percentiles of simulation replicates. Profiles are arithmetic means across 20 pairs in physical 20 kb bins, with simulated profiles also averaged across replicates. All panels share the 350–250,000-generation log scale of Fig. 3. The central 960 kb of each 3 Mb simulation is shown. Inference of the sweep does use neutral markers, whereas negative-selection inference includes selected and neutral sites. The parameter search trajectories are shown in Supplementary Fig. S5.

For *GRK2*, the purifying selection model exhibited an inferred-profile replicative error of 0.817, as opposed to 1.233 of its neutral fitted control. The true profile had an error of 1.108, thus demonstrating that a closer match of TMRCA need not imply an equally close match in underlying ancestry. At *TREM2*, the replicate profile error of the purifying-selection model was 0.598, versus 0.839 for its fitted neutral control model (evaluation was based on 24 replicates). This means that purifying selection by itself is capable of producing coalescence valleys similar to the ones observed empirically. These fits show that such a signature is not unique to positive selection.

### Detection and localisation depend on sweep history and score definition

We evaluated the gamma_smc TMRCA output and compared it with iHS and XP-EHH in a benchmark with 1,000 two-population simulations (Supplementary Fig. S7). Our simulations contained a known hard-sweep target, and used the gamma_smc with the methods aggregation approach (Methods). Incomplete and nearfixed histories were conditioned on specified final frequencies. We simulated both recently completed and older selective sweeps, in which the beneficial allele had reached 100% frequency a median of 688 and 2,688 generations before sampling, respectively.

We show how true and estimated TMRCA, iHS and XP-EHH vary around the known selected site (Supplementary Fig. S7a–d). We display the first replicate of each sweep history and use subsequently all replicates to estimate performance. Across the replicates, iHS near the selected site was weaker and less often computable after the beneficial allele reached fixation. For recently completed sweeps, gamma_smc estimated a median TMRCA of 1,311 generations in the target interval, compared with a true median of 1,745 generations. For older completed sweeps, the median estimate was 2,704 generations compared with a true median of 3,292 generations. The gamma_smc approach did indeed capture the decrease in the TMRCA associated with the sweep, but in the present benchmarks, it tended to underestimate TMRCA.

`gamma_smc` detected nearly all recently completed sweeps and all older completed sweeps in 5-kb target intervals, while iHS rarely detected either group (Supplementary Fig. S7). XP-EHH also detected nearly all recently completed sweeps, but detected fewer older sweeps. For incomplete sweeps, XP-EHH detected more replicates than either `gamma_smc` or iHS. Some of these differences reflected whether we could calculate each method’s detection statistic. After fixation, some target intervals contained no variants with usable iHS values. When we examined the flanking regions, it allowed us to evaluate iHS in every recently completed-sweep replicate and detect most of these sweeps. For `gamma_smc`, some narrow intervals contained fewer than the two variable sites required to summarise TMRCA. Increasing the intervals from 5 to 20,kb provided enough sites and enabled `gamma_smc` to detect every sweep approaching fixation or recently fixed. Wider intervals therefore generally improved detection in these simulations.

For the comparison of localization accuracy using windows of size 5kb, the median error for `gamma_smc` in sweeps close to fixation or already fixed was around 9 kb, whereas that for the strongest XP-EHH score in each window was around 20-30 kb. For incomplete sweeps, XP-EHH located the selected site slightly more accurately than `gamma_smc`.

We utilized neutral simulations to normalise the statistics, calibtrating detection thresholds for a 5% false-positive rate, yet still observed rates still differed between methods. `gamma_smc` and iHS stayed below this rate in the target intervals, while XP-EHH even produced no false positives in this experiment (Supplementary Tables S9–S11).

## Discussion

We present `gamma_smc_cu`, a GPU-based approach to make pairwise coalescence inference feasible at the population scale. Our efficiency benchmarking was carried out by comparing its performance on chromosome 22 where `gamma_smc_cu` runs through 63,190 pairs in 15.91 s compared to 17.9 minutes for the CPU version. We, then, continue with our software on 26 human populations from the 1000 Genomes Project and generated a list of 143 regions of interest with gene-level resolution in their coalescence time. Our contribution is therefore the combination of inference software with gene highlighting and benchmarking subsequent interpretations.

Fast TMRCA inference for each haplotype pair can enable many downstream applications, as shown for instance by the ASMC [4] method. It should be noted that `ASMC` requires a user-supplied demographic history, while `gamma_smc` uses a constant-size flow field as an assumption of the original algorithm [5] which we kept, since our aim was algorithmic parity with faster run-time. Our implementation accelerates the existing `gamma_smc` model and does not estimate a new population-size history or reconstruct a joint ancestral recombination graph. Under these assumptions genealogical variation is implicitly portrayed on a gene-level, more strictly on a per-site level, but we aggregate the model output based on described methods. In the end the user can designed the how to aggregate the inference output.

Genome-wide comparisons show that genes inside regions with higher recombination rates or higher predicted diversity under BGS (*B^′^*, indicating weaker BGS) overall have higher TMRCA ranks (Supplementary Fig. S6b). Gene length shows only a weak marginal association with TMRCA rank, but its partial correlation is negative after controlling for site count, recombination rate and *B^′^* (Supplementary Table S8). Recombination, background selection and gene architecture contribute to these ranks, so low TMRCA alone does not establish positive selection. The enrichment of strong iHS and XP-EHH signals among low-TMRCA genes also provides additional support for their prioritization as candidates for positive selection (Supplementary Fig. S6c).

As previously mentioned we use the geometric mean to summarise TMRCAs across all pairs. And we focus on the fraction of pairs with TMRCA below a chosen cutoff measuring how common recent coalescence is. Our benchmark also points out the distinction between detecting selection in a region and locating its target (Supplementary Fig. S7). XP-EHH was effective for near-fixed and completed sweeps, while flanking iHS often provided evidence of selection even when iHS could not be computed at the target. Our results describe that the use of `gamma_smc` does provide a reater sensitivity in particular for older more complete sweeps. It also gave smaller localisation errors in the near-fixed and recently completed sweep simulations compared to the strongest XP-EHH signal. This localisation advantage, however, did not extend to incomplete sweeps. Increased target interval lengths increased detection, which was a further demonstration of how spatial scale influences performance. In the end, it is always beneficial to use many mutually supportive statistics in selection detection and localization.

In addition, our filtering did not explicitly exclude introgressed regions such as from Neanderthals, and the geometric mean reduces the influence of deep coalescence between introgressed and other haplotypes. Comparing TMRCA patterns within and between ancestry backgrounds would help assess how introgression contributes to the candidate signals.

We further check how the same observed TMRCA profile can be generated by different selection histories(Fig. 4). Models containing only neutral as well as deleterious mutations produced inferred TMRCA drops resembling parts of the same empirical profiles fitted by sweeps. Thus, purifying selection can generate a similar inferred signature without any beneficial mutations. This is consistent with the reduction of linked neutral diversity through background selection [54, 55].

For *GRK2*, the simulated data match the observed TMRCA profile better when we use estimated TMRCA than when we use the true TMRCA recorded in the simulation. The apparent fit therefore depends partly on the inference method. The sweep and purifying-selection analyses also differ in their inputs, that is the former applies neutral markers, whereas the latter includes selected variants. We explored the models’ parameter spaces to different extents and used simplified demographic assumptions. Together with the differences described above, these choices limit our ability to determine which evolutionary process best explains the observed TMRCA landscape. Generally, the simulations show that purifying selection can provide an alternative explanation for a shallow inferred TMRCA profile.

*GRK2* is an example that demonstrates the approach to integrating TMRCA rankings, regional haplotypes, and previous sweeps scans to characterise a candidate region. This gene has maintained its low TMRCA percentile in GIH, and XP-EHH evidence, and coalescence minimum, also CLUES2 marker indication and aDNA help in understanding the variations within the region. Together, these analyses prioritise the region for follow-up and detect distinct positions of interest. Another contrasting example is provided by *TREM2*, where the low coalescence and haplotype structure of the locus exist independently of a TMRCA rank below 1% or strong XP-EHH signal in IBS.

The, here, compiled gene set and the West-Eurasian aDNA scan [15] offer complementary perspectives on selection, connecting present-day genealogical patterns with allele-frequency changes through time. Regional analyses at *GRK2* and the aDNA signal at *BPIFA2* make the value of combining these approaches apparent, while broader temporal and geographic sampling could help clarify the history of loci such as *TREM2*. The released TMRCA ranks let users examine candidates beyond those highlighted here and combine genealogical evidence with haplotype signals, data from other cohorts and future ancient-DNA time series.

Phased cohort populations that are larger and more varied will allow us to extend the gene list and take population structure into account. The benchmark is the starting point for assessing various scenarios of selection and demography, while BGS annotations specific to each population, varying-rate analysis, and studies excluding relatives may improve future research. In particular we show the importance of using fitting simulations for a systematic follow-up of candidates which provided a path towards more reliable evidence evaluation for selection in the face of multiple possible explanations. With our tool, we make pairwise analysis more efficient. Our software and highlighted genes provide a comprehensive blueprint for investigating evolutionary variation throughout diverse populations, and even non-human populations as well when adapting mutation and recombination rates.

## Methods

### gamma_smc model and the GPU inference implementation

Our work builds on the `gamma_smc` implementation of the pairwise sequentially Markovian coalescent (pairwise SMC) [56] framework developed by Schweiger and Durbin [5]. The pairwise SMC takes the observed data (the set of segregating sites) as an observation from a hidden Markov model (HMM) [57] whose hidden state *t* is the pairwise coalescence time. Unlike the commonly used approximation which discretizes *t* into bins, the `gamma_smc` represents the posterior distribution of *t* at each site as a gamma distribution. Forward transitions between subsequent sites are modeled by a flow field which takes the posterior of the current site to its next posterior state. The program `gamma_smc_cu` runs the entire process on a GPU, where each CUDA thread works on one pair of haplotypes. Input preprocessing is done per population.

When GPU memory is insufficient for a whole chromosome, `infer_blockwise` processes smaller, overlapping blocks. The overlap provides context for inference near block boundaries; results from the central regions are then combined into a chromosome-wide output.

For each chromosome and population, we estimate *θ̂* from mean within-diploid heterozygosity divided by the chromosome length. We set 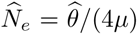, with mutation rate *µ* = 1.25 *×* 10*^−^*^8^ and recombination rate *r* = 10*^−^*^8^ per base per generation. Coalescent times are converted to generations with 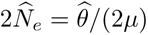.

### stdpopsim accuracy benchmark

For each of the 14 parameterizations in Fig. 1, we simulate one 5 Mb sequence and evaluate all 190 pairs of 20 haplotypes. We calculate pairwise Pearson correlation of natural-log TMRCA and 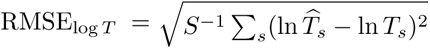.

### Population-scale runtime

To benchmark runtime, we use chromosome 22 from the YRI population containing 313,277 segregating sites and 356 haplotypes. We test the efficiency of the methods using six different numbers of pairs (178, 708, 3,460, 9,780, 31,684 and all 63,190 pairs). CUDA and CPU/ASMC timings were collected in separate runs on the same machine and dataset. We ran `gamma_smc` with one CPU process and ASMC with either one or eight CPU workers. `gamma_smc` saves parameters describing the posterior distribution, whereas `gamma_smc_cu` and ASMC save posterior means. The machine we used for testing has an Intel Xeon Gold 6148 CPU. CUDA uses one V100S-PCIE-32GB GPU, built with CUDA 12.1.

### Controlled detection and localisation benchmark

We tested how well `gamma_smc`, iHS and XP-EHH detect selective sweeps and locate the selected site using the same simulated datasets. We simulated two populations with SLiM, using a 1 Mb sequence, diploid *N_e_*= 10,000, and 100 sampled haplotypes per population. We ran the CPU version of `gamma_smc` on all 4,950 haplotype pairs in the focal population, defined as the population receiving the beneficial mutation in the sweep simulations. We tested whether each method detected selection in 5-kb and 20-kb windows centred on the selected site. We also searched a 500-kb region and measured the distance between the centre of each method’s strongest window and the selected site. Supplementary Section 4.2 provides the simulation and evaluation details.

### Processing of 1000 Genomes data

We used the 30*×* high-coverage data of the 1000 Genomes Project on the GRCh38 assembly [6]: 3,202 samples, including 602 complete trios without pedigree filtering, from 26 populations across 5 continental groups (7 AFR, 5 EUR, 5 EAS, 5 SAS, 4 AMR; Supplementary Table S1). Sample sizes within each population vary from 74 (ASW) to 179 (CEU), giving pair counts ranging from 10,878 to 63,903 per population (829,638 pairs total). Per-chromosome phased VCF files for all 22 autosomes were loaded using cyvcf2 v0.34.0 [58].

### Per-chromosome blockwise inference

For each of the 572 chromosome–population combinations, all within-population unordered haplotype pairs are enumerated. We use pair chunks of 512, a core size of 6,000,000 segregating sites and zero flank, so every chromosome fits in one inference block. This means we can make use of full-chromosome forward–backward context. All results include 22 autosomes and 26 populations, with 497,094 gene-population pairs, including gene–population combinations without a score because the gene contains fewer than two segregating sites.

### Gene-level aggregation: log-space (geometric mean)

For each gene-population combination with at least two segregating sites, the summary statistics are based on the geometric mean of per-pair TMRCAs in the gene region; combinations with fewer sites receive no score:

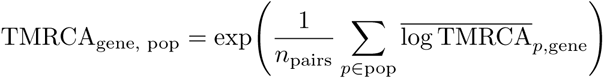

where 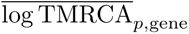 denotes the mean of log posterior mean TMRCA over all segregating sites in the gene body for pair *p*, using positive finite site-level posterior means in physical generations. A quantitative comparison of geometric-vs arithmetic-mean per-gene aggregation at known partial sweeps is given in Supplementary Table S12. In addition, for each gene we store the fraction frac_below_1000 of pairs in the population whose geometric-mean TMRCAs are less than 1,000 generations.

### Within-population gene-level TMRCA ranking and segmental duplication masking

Within each population, we rank genes by their TMRCA scores, assigning the same average rank to genes with identical scores. Genes with fewer than two segregating sites receive no score. After ranking, we exclude 1,123 genes with segmental duplications covering at least half of their gene body, leaving 17,996 genes.

### Candidate prioritisation

We use five stages of filtering described below.

1. Stage 1 filtering removes 1,123 SD genes, leaving 17,996.
2. Stage 2 keeps 620 genes below 1% in at least one population.
3. Stage 3 keeps 579 with all populations of at least one continental group below 5%.
4. Stage 4 excludes genes within 500 kb of the 23 reference loci, leaving 540.
5. Stage 5 clusters these genes by single linkage on each chromosome at a 1 Mb distance threshold, keeping the gene with the lowest minimum rank as representative.

The 143 resulting loci comprise 82 multi-gene clusters and 61 regions with only a single gene.

The seven case studies were selected post hoc using coalescence ranks, haplotype and allele-frequency summaries, regional method comparisons and biological context. Each case has a fixed focal population for its variant and haplotype summaries.

### Sensitivity to screening thresholds

We apply stage-2 cutoffs of 0.5%, 1%, 2% and 5% uniformly across the gene list, holding the SD mask, stage-3 5% continental rule, 500 kb reference exclusion and 1 Mb clustering fixed. These yield 58, 143, 233 and 277 loci. Separately, changing only the continental rule to 1% or 10% yields 52 or 154 loci, and changing only clustering to 500 kb or 2 Mb yields 159 or 124 loci. All eight focal genes are evaluated at every stage-2 cutoff.

### Choice of a representative gene

For each cluster, we choose the gene with the lowest TMRCA percentile in any population as its representative and use its name to label the cluster. All candidate genes belonging to each multi-gene cluster are listed in Supplementary Table S3.

### Cross-population replication scores

Population rank correlations are computed using 18,588 genes with finite ranks in all 26 populations. Following Galwey [46], an effective count summarises the eigenvalues of each continental rank-correlation matrix:

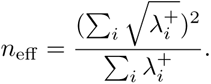

The effective counts are 1.916 (AFR), 1.683 (EUR), 1.753 (EAS), 1.617 (SAS) and 1.960 (AMR); the combined SAS+EUR matrix gives 2.578.

For gene *g* and continent *c*, let 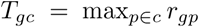 be the highest (least extreme) rank. We report 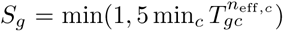 as a descriptive ordering score. The table provides this descriptive score and its rank among all genes. The score emphasises consistently low ranks within a continental group; a population-specific signal can have a weak score even if one population has a low rank. Candidate genes are selected using the filters described above, while this score summarises their patterns across populations.

### Variant-level confirmation

For the case-study genes, a summary of the variant-level evidence was produced from parsed genotypes derived from the 1000 Genomes phased VCF within *±*500 kb of the gene midpoint. For each biallelic variant in the window, we calculated (i) the focal-population VCF alternate-allele frequency *p_f_* and the corresponding frequency *p_o_* among all haplotypes outside the focal superpopulation, (ii) the per-site Hudson *F*_ST_, and (iii) the directional frequency difference Δ = *p_f_ − p_o_*. These are the comparisons in Tables 2 and 3; Figure 3e instead compares IBS with YRI, as labelled. Summary statistics per locus include the depleted-to-enriched site-count ratio *D/E*, where *D* = #*{i* : Δ*_i_ <* 0*}* and *E* = #*{i* : Δ*_i_ >* 0*}* (sites with Δ*_i_* = 0 contribute to neither count), the maximum and the average of the top 10 *F*_ST_ scores per locus, and the genomic coordinates/frequencies of the most differentiated variant (arg max*_i_ |*Δ*_i_|*). For example, the *GRK2* ratio is 21,445*/*3,028 *≈* 7.1. This ratio differs from Hudson’s *F*_ST_: it measures the direction of ALT-allele frequency differences rather than estimating population differentiation.

### Haplotype statistics

We calculated iHS and nSL [13, 59, 60] from the phased genotypes used for TMRCA inference, using selscan v3.0.0 separately for each chromosome and population. We removed population-monomorphic sites and used the original VCF reference/alternate allele coding.

We applied a 5% MAF threshold and default parameters for haplotype extension, along with a constant recombination rate of 1 cM/Mb for iHS. The iHS and nSL values were standardized independently for each chromosome and population using 20 allele-frequency bins of width 0.05. Bins containing fewer than 50 finite scores or with zero or nonfinite standard deviation were discarded. For each gene, we kept the highest absolute score and the proportion of valid scores that deviated from the mean by more than two standard deviations.

We estimated Garud’s *H*_12_ [61] using windows of 400 variants and moving the window forward by 50 variants at a time. For each gene, we kept the highest score obtained from windows centered within the gene body. For the neighbourhood analysis shown in Table S4, we used the same window size and step within *±*500 kb of each gene midpoint, including sites monomorphic in the focal population. Figure 3i,j also includes these sites. Percentiles compare genes with finite scores within the same population and analysis. Higher neighbourhood *H*_12_ and *H*_2_*/H*_1_ percentiles indicate larger values. We calculated XP-EHH with scikit-allel [62], comparing pooled SAS or EAS haplotypes with YRI at biallelic sites polymorphic in both groups. For each comparison, XP-EHH integration spanned the queried genes on each chromosome and their 500,kb flanks, with an additional 2,Mb at both ends of the combined region. Table S4 reports mean raw XP-EHH across each gene and its 500,kb flanks. We also calculated these statistics for five arbitrarily selected comparison genes on other chromosomes.

We used Relate v1.2.4 to infer local genealogies and population-specific demographic histories, with five demographic estimation iterations, a mutation rate of 1.25 *×* 10*^−^*^8^ per base per generation and 28 years per generation. For *BPIFA2* /GIH, *CCDC92* /CDX and *CLEC6A*/CDX, we considered biallelic SNPs available in the Relate input with an unambiguous ancestral allele. Within each neighbourhood, we chose the SNP with the largest absolute difference in ALT-allele frequency between the focal population and populations outside its continental group. For each selected variant, CLUES2 [49, 50] estimated allele-frequency trajectories and selection coefficients from 200 posterior samples of its local genealogy. We used 200 allele-frequency states, analysed the past 2,000 generations and assumed *h* = 0.5 with *|s| <* 0.1. We fixed the present-day derived-allele frequency to its observed value and used approximate confidence intervals from CLUES2. Likelihood-ratio tests provide nominal evidence at loci and variants selected after examining the data. Supplementary Table S7 reports all 13 fits and sample details.

We compared ALT-allele frequencies between GIH and pooled non-SAS haplotypes and between IBS and YRI. We inferred trajectories for focal variants, with *C11orf65* in GIH included for comparison (Fig. 3).

### Genomic factors associated with TMRCA ranks

We also studied specifically how TMRCA ranks relate to gene length, site count, recombination, background selection and haplotype statistics (Supplementary Fig. S6). We included genes with complete annotations and analysed gene bodies and non-overlapping flanking regions separately. Supplementary Methods, Section 4.1, describes the comparisons, controls and uncertainty estimates.

### Recombination context at highlighted genes

We inferred population-specific recombination maps with fastrho v0.1.1 [63], using the domain-randomized-v1 checkpoint and its matching feature statistics in phased mode. We analysed an exploratory set of 242 protein-coding genes, including the case studies and comparison genes in Tables 1–2. Focal populations were chosen using minimum TMRCA ranks in an exploratory target-selection analysis, with specified case-study and comparison populations taking precedence. Adding YRI as a comparison gives 480 distinct gene– population maps. We inferred these maps from the high-coverage GRCh38 phased data using individuals included in 1000 Genomes Phase 3, specifically a subset of the 3,202-sample TMRCA cohort. Within each population, we used biallelic SNPs and excluded sites with missing genotypes, unphased heterozygotes or duplicate positions. We included all remaining polymorphic sites regardless of allele frequency or spacing.

Regions plotted extended 500 kb upstream and downstream from each gene, apart from the case-study loci *GRK2* and *TREM2*, for which we employed the windows indicated in Fig. 3. Inference was done with an additional 500 kb of surrounding sequence on both sides and a mutation rate of 1.25 *×* 10*^−^*^8^ per base per generation. The *ρ* values were mapped into recombination rates using *r* = *ρ/*(4*N_e_*), with *N_e_* = 10,000 to compare them with the constant rate used by `gamma_smc`. We averaged rates in 10-kb and 50-kb windows, weighting each estimate by the length of its overlap with the window.

We also used background-selection (BGS) predictions from the published CADD 6%/deCODE model of Buffalo and Kern [55], evaluated for YRI and CEU in 100-kb bins (Dryad doi:10.5061/dryad.qnk98sfnv).

Panels 3o,p show predictions for CEU and YRI as European and African reference populations, respectively. Gene-level annotations derived from these maps use the YRI predictions. Together, the BGS and recombination annotations provide context for the coalescence signal.

### Simulating and matching genomic TMRCA profiles

At each locus, we fitted simulations to the mean TMRCA for the same set of 20 haplotype pairs considered in the focal analysis, averaged over 20 kb bins within a 960 kb region. The simulations involved generating data on 3 Mb of the genome surrounding each gene midpoint, sampling 206 haplotypes for GIH or 314 for IBS and using the deCODE sex-averaged GRCh38 recombination map and a mutation rate of 1.25 *×* 10*^−^*^8^ per base per generation. We inferred TMRCA using the same population-specific *N_e_* values as in the full-chromosome analysis and a constant recombination rate of 10*^−^*^8^. Empirical inference used full chromosomes, whereas simulated inference used the 3 Mb regions.

We simulated conditioned sweeps with msprime [16] and purifying selection with SLiM [64] through stdpopsim [65]. Purifying selection acted on exons and conserved sequence or a fitted synthetic target, with fixed *s <* 0 and *h* = 0.5. The simulation parameters were chosen by optimising a weighted log-error objective function. The parameter `gamma_smc_cu` used the simulated genotypes to estimate the TMRCA. In the SI material, we describe the fitting procedure in more depth (see Fig. S5).

## Declarations

### Data availability

Source code and Python bindings for `gamma_smc_cu` are available at https://github.com/kevinkorfmann/gamma_smc_cu, with documentation at https://gamma-smc-cu.readthedocs.io/en/latest/. The 1000 Genomes high-coverage data are available from https://www.internationalgenome.org, gene annotations from GENCODE v46 (https://www.gencodegenes.org/human/release_46.html), and segmental duplication annotations from the UCSC genomicSuperDups track (https://hgdownload.soe.ucsc.edu/goldenPath/hg38/database/).

### Authors’ contributions

KK conceived the study, designed and implemented `gamma_smc_cu`, performed the scans and analyses, generated all figures, and wrote the manuscript. SM supervised the project, provided guidance on the HMM implementation, and edited the manuscript. All authors read and approved the final manuscript.

### Generative AI usage statement

We used AI tools to develop code, check numerical results, search and review literature, and edit the manuscript. We verified the outputs and take full responsibility for the work.

## Acknowledgements

We thank Iain Mathieson, Neda Rahnamae, Matteo Fumagalli, and Avery Selberg for input on various stages of the manuscript.

