## Supplementary Information for "Fast pairwise coalescence enables gene-resolution scans for selection in diverse human populations"

### 1 Supplementary Figures

#### 1.1 Per-pair TMRCA histograms at reference loci

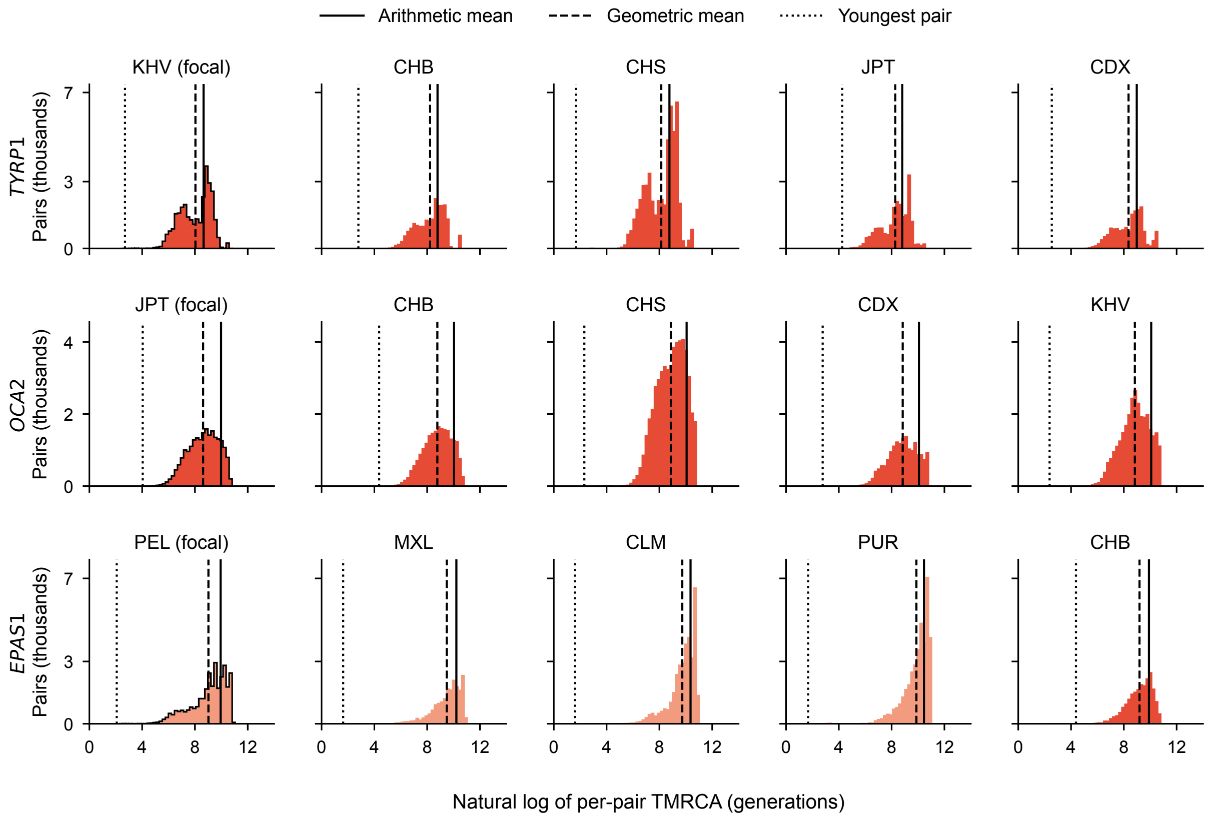

Figure S1: **Pairwise TMRCA distributions at three reference loci.** Rows show *TYRP1*, *OCA2* and *EPAS1*. Each in five populations. The focal population from Table 1 appears first with a black outline. Fill colours identify continental groups (EAS coral, AMR peach). Histograms show the natural log of each pair's geometric-mean TMRCA across the gene, with counts in thousands. Solid and dashed lines mark the all-site/pair arithmetic and geometric summaries, respectively; the dotted line marks the minimum per-pair geometric mean. The solid line shows the mean posterior estimate across sites. In the focal populations, the lowest geometric means across sites for individual pairs are approximately 15, 57, and 8 physical generations. Across the five panels shown here, a total of 61 pairs fall below the histogram range, see TYRP1/CHS (15), EPAS1/PEL (4), EPAS1/MXL (14), EPAS1/CLM (15), and EPAS1/PUR (13). Counts below and above the range are recorded separately, with an overflow count of zero. Looking at TYRP1, which falls below a 10% threshold, while OCA2 and EPAS1 remain above it.

36 ## 1.2 Three-method per-pair TMRCA comparison at 8 loci

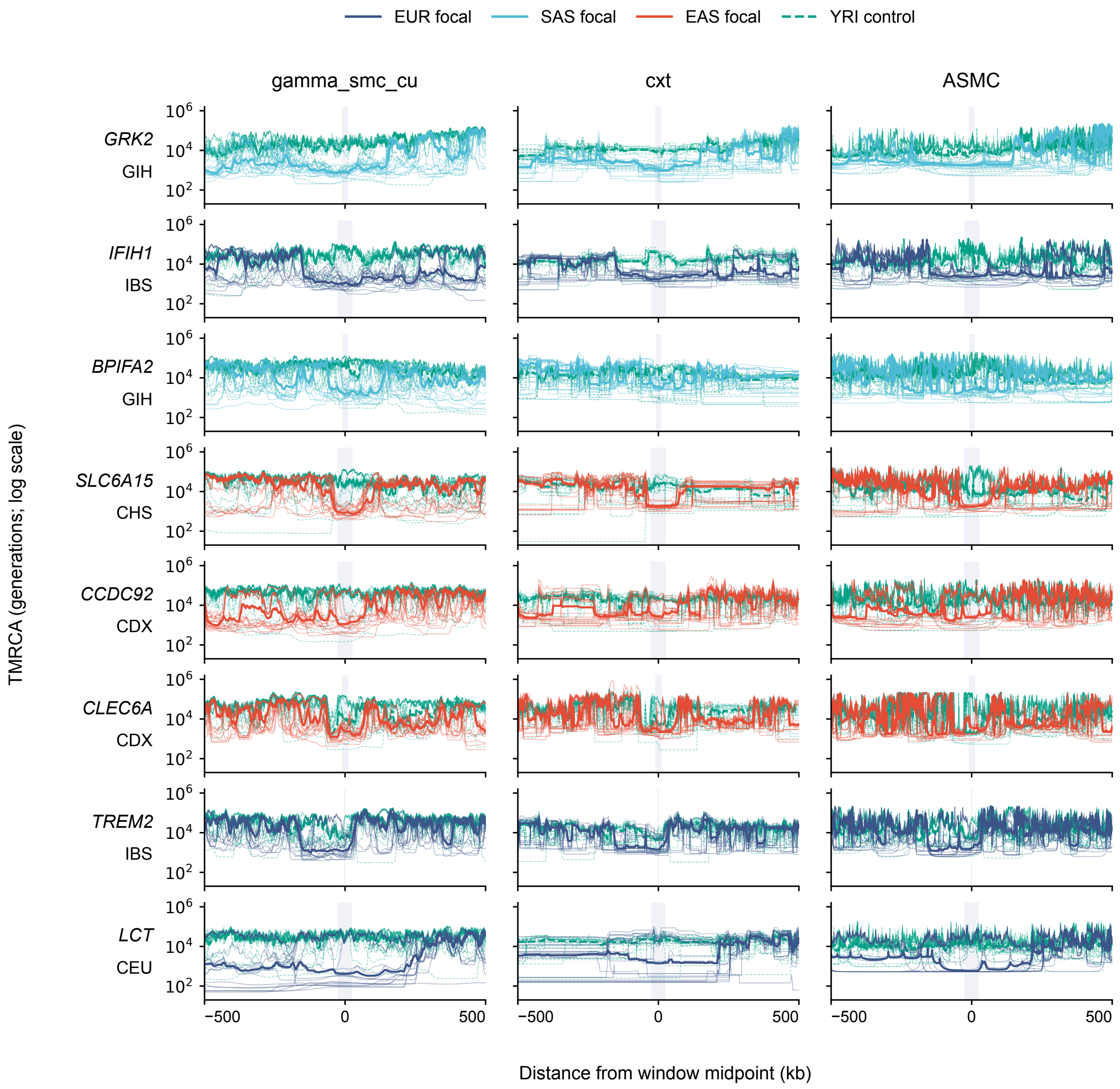

Figure S2: **Three-method TMRCA comparison at eight case-study loci.** Columns show `gamma_smc_cu` pairwise posterior means in physical generations, `cxt` regional estimates and `ASMC` pairwise posterior means. Thin traces show 20 focal-population pairs (EUR navy, SAS cyan or EAS coral) and 20 YRI control pairs (teal); thicker solid and dashed curves show their respective pointwise medians on a log scale. All eight loci have YRI estimates from all three methods in this comparison. Pale bands mark gene intervals – all panels span 1 Mb centred on the decoded window midpoint. The final row shows the *LCT* positive control.

#### 1.3 Gene-level TMRCA localisation within the chr11q13.2 cluster

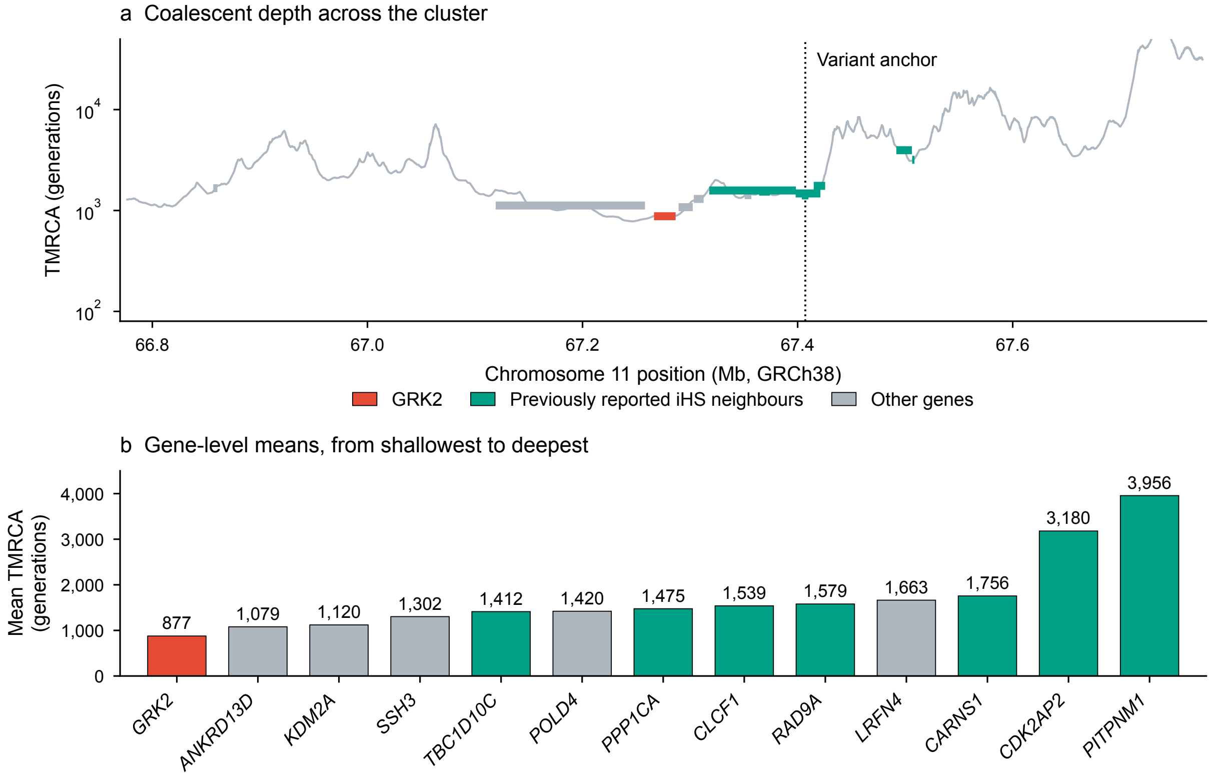

Figure S3: **Gene-level coalescent depth across 13 genes in the *GRK2* 1 Mb neighbourhood in GIH.** (a) Per-site geometric-mean TMRCA across 20 within-GIH pairs, with horizontal bars showing each gene's arithmetic mean across these sites. The dotted line marks the $\Delta$AF/`CLUES2` anchor at chr11:67,407,126, about 120 kb downstream of *GRK2*; the *GRK2* gene body lacks variants at MAF$\geq$5% suitable for iHS. (b) The same gene-level means, ordered from shallowest to deepest; *GRK2* has the lowest value (877 physical generations). Coral: *GRK2*, teal: seven neighbouring genes annotated in the two regional candidate windows of Voight *et al.* [1], and grey: other genes.

38 ## 1.4 TREM cluster paralog layout

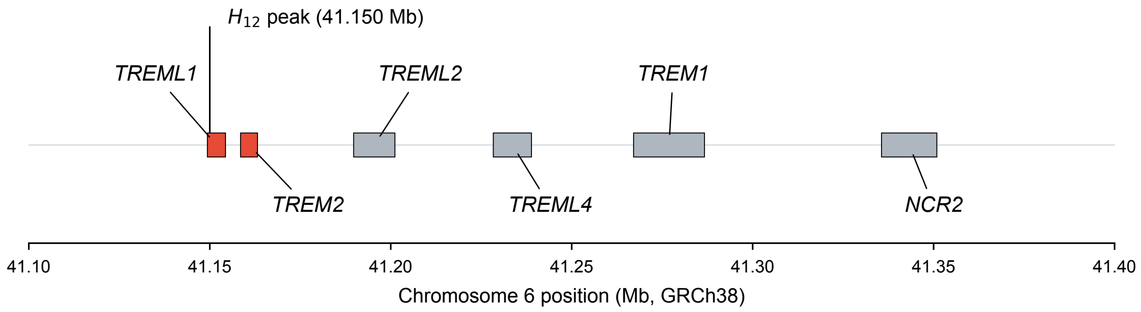

Figure S4: **The genomic organisation of the TREM region around a 300 kb interval on chromosome 6.** Coral blocks show *TREML1* and *TREM2*, and grey blocks show neighbouring genes. The vertical marker presents the $H_{12}$ peak at 41.150 Mb (GRCh38).

#### 1.5 Focal selection-model parameters and search trajectories

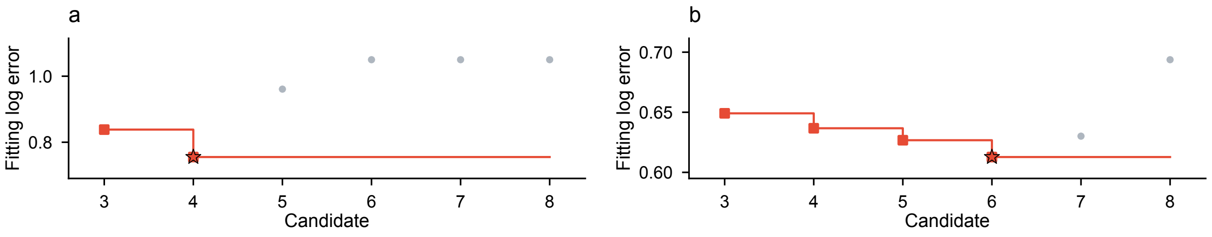

| | |
|---|---|
| Model | Hard sweep |
| Final candidate | #4 |
| Generating $N_e$ | 25,000 |
| Genic s | 0.05 |
| Since sweep endpoint | 2,500 gen |
| Selected site (GRCh38) | 67,226,514 bp |
| Centre offset | −50.000 kb |
| Start → end frequency | 0.000020 → 0.999980 |
| Simulation engine | msprime; conditioned sweep |
| Haplotypes / pairs | 206 / 20 |
| Inference $N_e$ | 15,686.21 |
| Fitting / displayed replicates | 8 / 24 |
| Displayed ensemble | Fresh validation |
| Model / neutral log error | 0.905 / 1.211 |

| | |
|---|---|
| Model | Hard sweep |
| Final candidate | #6 |
| Generating $N_e$ | 15,302.26 |
| Genic s | 0.0060491 |
| Since sweep endpoint | 2.995 gen |
| Selected site (GRCh38) | 41,123,226 bp |
| Centre offset | −37.620 kb |
| Start → end frequency | 0.000033 → 0.999967 |
| Simulation engine | msprime; conditioned sweep |
| Haplotypes / pairs | 314 / 20 |
| Inference $N_e$ | 16,767.68 |
| Fitting / displayed replicates | 8 / 24 |
| Displayed ensemble | Fresh validation |
| Model / neutral log error | 0.626 / 0.899 |

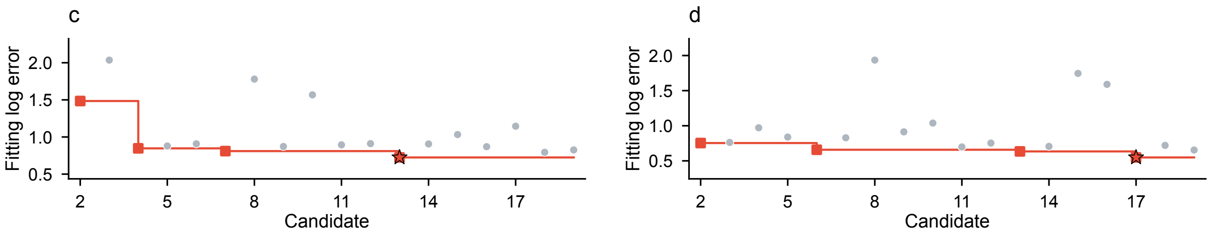

| | |
|---|---|
| Model | Negative selection |
| Final candidate | #13 |
| Generating $N_e$ | 16,000 |
| Homozygous s / h | −0.000351525 / 0.5 |
| Deleterious fraction | 1 |
| Synthetic target width | 500 kb |
| Centre offset | −50.000 kb |
| Q / simulated diploids | 10 / 1,600 |
| Forward burn-in | 10 $N_e$ = 160,000 gen |
| Haplotypes / pairs | 206 / 20 |
| Inference $N_e$ | 15,686.21 |
| Fitting / displayed replicates | 3 / 24 |
| Displayed ensemble | Fresh validation |
| Model / neutral log error | 0.817 / 1.233 |

| | |
|---|---|
| Model | Negative selection |
| Final candidate | #17 |
| Generating $N_e$ | 21,160 |
| Homozygous s / h | −0.005 / 0.5 |
| Deleterious fraction | 0.7 |
| Synthetic target width | 160 kb |
| Centre offset | −50.000 kb |
| Q / simulated diploids | 10 / 2,116 |
| Forward burn-in | 10 $N_e$ = 211,600 gen |
| Haplotypes / pairs | 314 / 20 |
| Inference $N_e$ | 16,767.68 |
| Fitting / displayed replicates | 3 / 24 |
| Displayed ensemble | Fresh validation |
| Model / neutral log error | 0.598 / 0.839 |

Figure S5: **Search trajectories and final simulation parameters.** Panels match Fig. 4: (a,b) *GRK2* and *TREM2* sweeps; (c,d) their purifying (negative) selection fits. Grey points show the tested selection models, the coral line shows the lowest fitting error reached so far, squares mark improvements, and stars mark the final models. Models keep their original numbering, with neutral controls omitted. Sweep panels show the final stage of parameter fitting. The table below each panel gives the parameters in physical units and the numbers of replicates used for fitting and evaluation. Sweep age is the time since the selected allele reached its final frequency. Synthetic blocks are intervals where we simulated deleterious mutations. $Q = 10$ is the approximate rescaling factor used in `SLiM`, and $N_e$ is the diploid effective population size. Each final model was evaluated using 24 new replicates. The objective, common settings and interpretation are described in Supplementary Methods (Section 4.3).

#### 1.6 Genomic covariates and matched haplotype comparisons

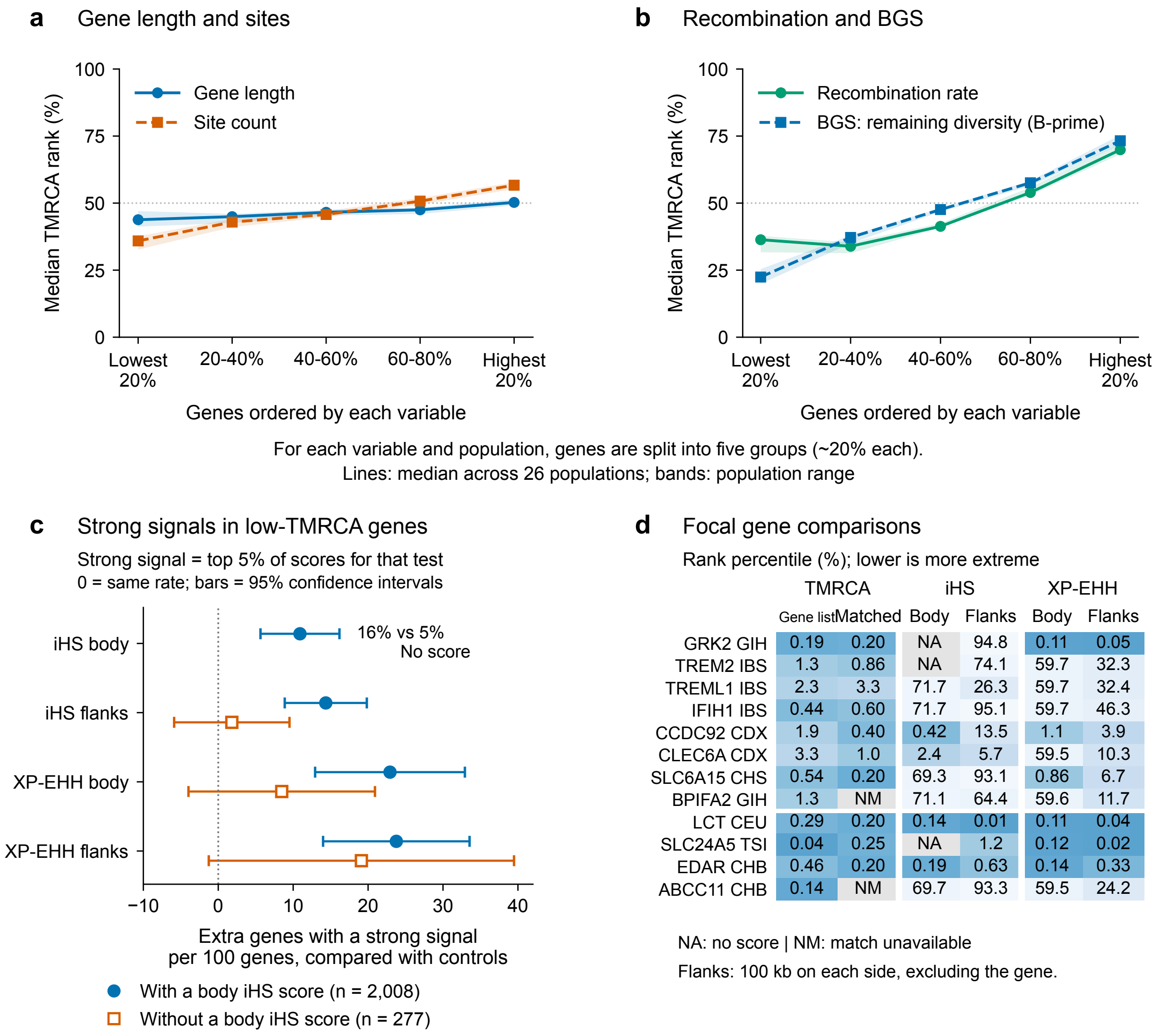

Figure S6: **Genomic covariate dependence and matched haplotype comparisons.** (a,b) Median TMRCA rank in five ordered groups containing approximately 20% of eligible genes each, formed separately for each variable and population. Lines show medians across 26 populations and bands show their range. Lower B-prime denotes stronger BGS. (c) Difference in top-5% haplotype-summary frequency between low-TMRCA genes (original rank $< 1\%$) and covariate-matched other-chromosome controls. Units are the number of extra genes with strong scores per 100 genes. Filled circles represent the 2,008 matched gene–population cases with iHS scores within the gene body. Open squares represent 277 cases without these scores. Bars are 95% chromosome-jackknife intervals with matches refitted. Missing scores are excluded, not encoded as zero. (d) Focal rank percentiles, with lower values more extreme. Matched TMRCA ranks are descriptive and do not replace the original screen. NA denotes no finite score; NM denotes insufficient matching or map coverage. Haplotype summaries are fractions of finite sites with $|\text{iHS}| > 2$ or normalized XP-EHH $> 2$, ranked with average ties. Flanks comprise the disjoint 100 kb on each side. XP-EHH compares each population with YRI, except YRI with CEU; positive scores favour focal haplotypes. XP-EHH was recalculated from the corrected haplotypes. iHS uses the earlier scores at sites present in the corrected SNP data. These comparisons do not calibrate selection specificity or power. Definitions, coverage, sensitivities and source tables are described in Supplementary Methods, Section 4.1.

#### 1.7 Sweep signatures and detection performance

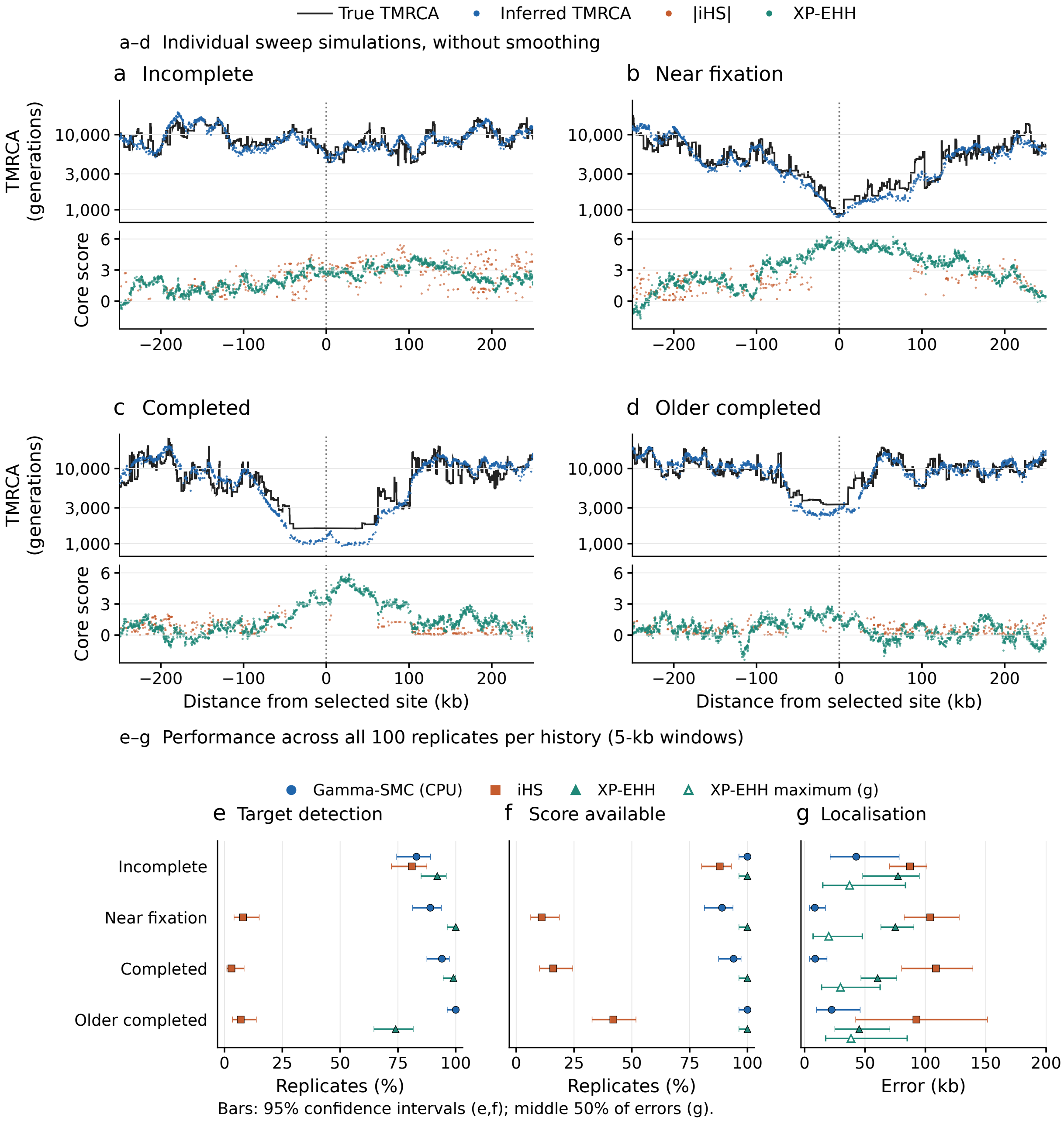

Figure S7: Sweep signatures and quantitative performance in controlled simulations. (a–d) The first evaluation replicate of each history, chosen by identifier without visual selection. Upper tracks show true geometric-mean TMRCA across all 4,950 focal pairs in each marginal tree (black) and `gamma_smc` estimates at SNPs (blue). Lower tracks show unsmoothed, finite core scores. The dotted line marks the selected site, excluded from the assayed markers. iHS requires focal MAF$\geq$ 5%; missing cores remain absent. (e,f) Detection and score availability in 5-kb target intervals across all 100 replicates per history, with Wilson 95% intervals. Missing scores count as non-detections. Thresholds were set using a separate set of neutral simulations to target a 5% false-positive rate. Table S9 reports the rates observed in the neutral evaluation simulations. (g) Median localisation error and interquartile range over all computable replicates in a 500-kb scan. Each replicate includes all tied best windows. Filled haplotype symbols in e–g use fractions of finite cores with |iHS| $>$ 2 or XP-EHH $>$ 2; open triangles in g use maximum XP-EHH. Wider windows, scan detection and other score summaries are reported in Tables S9–S11. The model uses constant population sizes and uniform recombination.

### 2 Supplementary Tables

#### 2.1 1000 Genomes populations: sample sizes, pair counts, and continental groups

Table S1: **1000 Genomes 30× high-coverage populations used in this study.** Pair count = $n(n-1)/2$ where $n$ is the number of haplotypes (2× samples).

| Code | Population | Continental | Samples | Pairs |
|---|---|---|---|---|
| ACB | African Caribbeans in Barbados | AFR | 116 | 26,796 |
| ASW | African Ancestry in Southwest US | AFR | 74 | 10,878 |
| ESN | Esan in Nigeria | AFR | 149 | 44,253 |
| GWD | Gambian in Western Division, Mandinka | AFR | 178 | 63,190 |
| LWK | Luhya in Webuye, Kenya | AFR | 99 | 19,503 |
| MSL | Mende in Sierra Leone | AFR | 99 | 19,503 |
| YRI | Yoruba in Ibadan, Nigeria | AFR | 178 | 63,190 |
| CEU | Utah Residents w/Northern & Western Eur. ancestry | EUR | 179 | 63,903 |
| FIN | Finnish in Finland | EUR | 99 | 19,503 |
| GBR | British in England and Scotland | EUR | 91 | 16,471 |
| IBS | Iberian populations in Spain | EUR | 157 | 49,141 |
| TSI | Toscani in Italia | EUR | 107 | 22,791 |
| CDX | Chinese Dai in Xishuangbanna, China | EAS | 93 | 17,205 |
| CHB | Han Chinese in Beijing, China | EAS | 103 | 21,115 |
| CHS | Southern Han Chinese | EAS | 163 | 52,975 |
| JPT | Japanese in Tokyo, Japan | EAS | 104 | 21,528 |
| KHV | Kinh in Ho Chi Minh City, Vietnam | EAS | 122 | 29,646 |
| BEB | Bengali from Bangladesh | SAS | 131 | 34,191 |
| GIH | Gujarati Indians from Houston, TX | SAS | 103 | 21,115 |
| ITU | Indian Telugu from the UK | SAS | 107 | 22,791 |
| PJL | Punjabi from Lahore, Pakistan | SAS | 146 | 42,486 |
| STU | Sri Lankan Tamil from the UK | SAS | 114 | 25,878 |
| CLM | Colombians from Medellín, Colombia | AMR | 132 | 34,716 |
| MXL | Mexican Ancestry from Los Angeles, USA | AMR | 97 | 18,721 |
| PEL | Peruvians from Lima, Peru | AMR | 122 | 29,646 |
| PUR | Puerto Ricans from Puerto Rico | AMR | 139 | 38,503 |
| **Total** | | | **3,202** | **829,638** |

#### 2.2 The 143-locus reference catalogue

Table S2 lists the 143 representatives from the reference screen: 82 multi-gene clusters and 61 singletons. Single-linkage clustering includes only genes that pass stages 1–4, so a singleton may have nearby genes that failed these filters. Table S3 lists the genes in each multi-gene cluster. The case studies were chosen post hoc for detailed follow-up and include loci outside this screen.

Table S2: **The 143 loci identified by the 1% reference screen.** Rank is the minimum within-population percentile across all 26 populations; the representative is the lowest-ranked stage-4 member. $n$ is the number of stage-4 genes in the distance cluster. Score is the descriptive continental replication score (Methods), not a $p$-value or FDR. The case studies were selected post hoc for detailed follow-up and need not belong to this catalogue.

| # | Representative | Chr | Start (Mb) | Pop | Rank (%) | $n$ | Score |
|---|---|---|---|---|---|---|---|
| 1 | *SHCBP1* | 16 | 46.579 | PEL | 0.027 | 9 | 6.1e-06 |

*Continued*

| # | Representative | Chr | Start (Mb) | Pop | Rank (%) | $n$ | Score |
|---|---|---|---|---|---|---|---|
| 2 | *OR4C6* | 11 | 55.662 | LWK | 0.084 | 1 | 1.6e-05 |
| 3 | *CTF1* | 16 | 30.897 | CHB | 0.101 | 26 | 0.0014 |
| 4 | *SLC9A5* | 16 | 67.238 | JPT | 0.122 | 41 | 7.8e-05 |
| 5 | *EXOC6B* | 2 | 72.176 | JPT | 0.128 | 3 | 0.00013 |
| 6 | *DCAF4L1* | 4 | 41.982 | CHB | 0.149 | 4 | 0.00066 |
| 7 | *P4HTM* | 3 | 48.990 | CDX | 0.149 | 49 | 0.001 |
| 8 | *BCL2L1* | 20 | 31.664 | JPT | 0.160 | 3 | 0.00037 |
| 9 | *PPM1D* | 17 | 60.600 | GBR | 0.170 | 6 | 0.00019 |
| 10 | *BMI1* | 10 | 22.321 | FIN | 0.181 | 4 | 0.00032 |
| 11 | *GRK2* | 11 | 67.266 | GIH | 0.191 | 16 | 0.00043 |
| 12 | *PLA2G10* | 16 | 14.673 | LWK | 0.201 | 3 | 6.9e-05 |
| 13 | *PSMB2* | 1 | 35.600 | TSI | 0.207 | 17 | 0.00053 |
| 14 | *CENPW* | 6 | 126.340 | JPT | 0.213 | 3 | 0.00078 |
| 15 | *PPP3CB* | 10 | 73.436 | TSI | 0.228 | 13 | 0.0004 |
| 16 | *ORC2* | 2 | 200.909 | JPT | 0.234 | 3 | 0.00062 |
| 17 | *PIGV* | 1 | 26.787 | LWK | 0.243 | 11 | 0.00032 |
| 18 | *TSSK3* | 1 | 32.352 | ITU | 0.254 | 5 | 8.5e-05 |
| 19 | *FBXO4* | 5 | 41.925 | CHS | 0.254 | 3 | 0.001 |
| 20 | *PRMT9* | 4 | 147.638 | FIN | 0.255 | 2 | 0.00043 |
| 21 | *VPS37C* | 11 | 61.130 | JPT | 0.255 | 3 | 0.0012 |
| 22 | *VDAC3* | 8 | 42.392 | PEL | 0.260 | 8 | 0.00012 |
| 23 | *UFM1* | 13 | 38.350 | FIN | 0.276 | 1 | 0.00074 |
| 24 | *PSME3* | 17 | 42.824 | TSI | 0.281 | 9 | 0.00074 |
| 25 | *FMNL3* | 12 | 49.636 | CHB | 0.292 | 7 | 0.0026 |
| 26 | *TMEM145* | 19 | 42.313 | JPT | 0.314 | 19 | 0.0029 |
| 27 | *DOK1* | 2 | 74.549 | GBR | 0.324 | 5 | 0.00054 |
| 28 | *PHYHIPL* | 10 | 59.177 | PEL | 0.334 | 1 | 0.0054 |
| 29 | *ERMN* | 2 | 157.319 | IBS | 0.339 | 2 | 0.0023 |
| 30 | *BUB1* | 2 | 110.638 | TSI | 0.350 | 1 | 0.00079 |
| 31 | *ZNF546* | 19 | 39.984 | FIN | 0.351 | 3 | 0.0043 |
| 32 | *PLPP5* | 8 | 38.263 | ESN | 0.370 | 7 | 0.00071 |
| 33 | *ZNF510* | 9 | 96.755 | IBS | 0.370 | 2 | 0.001 |
| 34 | *RNF11* | 1 | 51.236 | CLM | 0.381 | 6 | 0.00099 |
| 35 | *AMBRA1* | 11 | 46.396 | CDX | 0.383 | 4 | 0.0036 |
| 36 | *OTX1* | 2 | 63.050 | MSL | 0.386 | 2 | 0.00057 |
| 37 | *KCNE2* | 21 | 34.364 | ASW | 0.396 | 1 | 0.0005 |
| 38 | *KIF23* | 15 | 69.414 | TSI | 0.414 | 2 | 0.003 |
| 39 | *OSR2* | 8 | 98.944 | PJL | 0.418 | 1 | 0.00099 |
| 40 | *EPN2* | 17 | 19.216 | TSI | 0.419 | 1 | 0.0022 |
| 41 | *KMT2A* | 11 | 118.436 | PEL | 0.429 | 3 | 0.00032 |
| 42 | *GOLM2* | 15 | 44.289 | PEL | 0.435 | 3 | 0.00083 |
| 43 | *GLMN* | 1 | 92.246 | JPT | 0.436 | 5 | 0.00094 |
| 44 | *SGCB* | 4 | 52.021 | ESN | 0.438 | 4 | 0.0021 |
| 45 | *IFIH1* | 2 | 162.267 | IBS | 0.439 | 3 | 0.00082 |
| 46 | *KAT6B* | 10 | 74.825 | FIN | 0.446 | 1 | 0.0081 |
| 47 | *ALG2* | 9 | 99.216 | ESN | 0.449 | 2 | 0.00081 |
| 48 | *PPP1R42* | 8 | 66.964 | GBR | 0.451 | 6 | 0.0034 |
| 49 | *MRPS5* | 2 | 95.085 | YRI | 0.454 | 12 | 0.00031 |
| 50 | *SYF2* | 1 | 25.222 | BEB | 0.455 | 2 | 0.0018 |
| 51 | *APOF* | 12 | 56.361 | ITU | 0.461 | 8 | 0.0016 |
| 52 | *C2orf66* | 2 | 196.804 | CHS | 0.477 | 3 | 0.00093 |
| 53 | *PCYT2* | 17 | 81.901 | PUR | 0.486 | 6 | 0.0045 |
| 54 | *PYGO2* | 1 | 154.957 | PEL | 0.488 | 5 | 0.0019 |
| 55 | *TRIR* | 19 | 12.731 | GIH | 0.488 | 2 | 0.0018 |
| 56 | *ENSG00000266086* | 17 | 57.989 | KHV | 0.488 | 3 | 0.0033 |
| 57 | *METTL25* | 12 | 82.359 | FIN | 0.499 | 1 | 0.002 |

*Continued*

| # | Representative | Chr | Start (Mb) | Pop | Rank (%) | $n$ | Score |
|---|---|---|---|---|---|---|---|
| 58 | *CELF6* | 15 | 72.285 | IBS | 0.503 | 5 | 0.0055 |
| 59 | *ZHX1* | 8 | 123.248 | BEB | 0.503 | 3 | 0.0006 |
| 60 | *PCLAF* | 15 | 64.364 | PEL | 0.504 | 5 | 0.0041 |
| 61 | *RIPPLY2* | 6 | 83.853 | IBS | 0.508 | 2 | 0.013 |
| 62 | *NIT1* | 1 | 161.118 | YRI | 0.512 | 8 | 0.0015 |
| 63 | *HECTD2* | 10 | 91.409 | ITU | 0.513 | 1 | 0.0012 |
| 64 | *FABP5* | 8 | 81.281 | PJL | 0.518 | 1 | 0.0018 |
| 65 | *SLC6A15* | 12 | 84.859 | KHV | 0.520 | 1 | 0.0015 |
| 66 | *HS2ST1* | 1 | 86.915 | CHS | 0.525 | 1 | 0.0027 |
| 67 | *TRUB1* | 10 | 114.938 | KHV | 0.525 | 1 | 0.0046 |
| 68 | *RUNX1T1* | 8 | 91.955 | JPT | 0.527 | 1 | 0.002 |
| 69 | *PROCA1* | 17 | 28.703 | LWK | 0.539 | 8 | 0.00076 |
| 70 | *NME9* | 3 | 138.261 | PEL | 0.551 | 1 | 0.013 |
| 71 | *FBXO48* | 2 | 68.459 | JPT | 0.559 | 1 | 0.0016 |
| 72 | *GTSE1* | 22 | 46.297 | BEB | 0.561 | 4 | 0.0018 |
| 73 | *PDZD9* | 16 | 21.984 | MXL | 0.567 | 3 | 0.0013 |
| 74 | *FKBP3* | 14 | 45.116 | PUR | 0.576 | 5 | 0.0017 |
| 75 | *CYP3A5* | 7 | 99.648 | CEU | 0.577 | 4 | 0.0022 |
| 76 | *MCM4* | 8 | 47.960 | GBR | 0.579 | 3 | 0.002 |
| 77 | *IGIP* | 5 | 140.126 | GWD | 0.586 | 8 | 0.00063 |
| 78 | *GPR179* | 17 | 38.325 | PEL | 0.594 | 1 | 0.035 |
| 79 | *GRSF1* | 4 | 70.816 | GBR | 0.595 | 2 | 0.012 |
| 80 | *MRPL16* | 11 | 59.806 | TSI | 0.599 | 3 | 0.014 |
| 81 | *PPARD* | 6 | 35.343 | FIN | 0.606 | 1 | 0.037 |
| 82 | *H2BC21* | 1 | 149.884 | ITU | 0.609 | 3 | 0.0017 |
| 83 | *POU5F2* | 5 | 93.733 | GIH | 0.615 | 1 | 0.0043 |
| 84 | *HEMGN* | 9 | 97.927 | YRI | 0.618 | 1 | 0.0018 |
| 85 | *C6orf52* | 6 | 10.671 | CLM | 0.618 | 2 | 0.003 |
| 86 | *FEV* | 2 | 218.981 | FIN | 0.627 | 2 | 0.0062 |
| 87 | *TMEM168* | 7 | 112.762 | KHV | 0.632 | 1 | 0.0067 |
| 88 | *BMP2K* | 4 | 78.776 | CEU | 0.641 | 1 | 0.0041 |
| 89 | *ZNF660* | 3 | 44.578 | CHS | 0.641 | 3 | 0.004 |
| 90 | *OR1L4* | 9 | 122.724 | PJL | 0.645 | 2 | 0.0054 |
| 91 | *SPINK13* | 5 | 148.268 | TSI | 0.652 | 1 | 0.0027 |
| 92 | *DCN* | 12 | 91.140 | FIN | 0.653 | 1 | 0.0058 |
| 93 | *KLLN* | 10 | 87.859 | ACB | 0.654 | 1 | 0.00072 |
| 94 | *SOX4* | 6 | 21.594 | PUR | 0.660 | 1 | 0.0065 |
| 95 | *ATP1A1* | 1 | 116.373 | PJL | 0.661 | 1 | 0.0046 |
| 96 | *B3GALT9* | 9 | 120.792 | BEB | 0.662 | 1 | 0.011 |
| 97 | *CEP95* | 17 | 64.507 | STU | 0.677 | 4 | 0.003 |
| 98 | *PEG10* | 7 | 94.656 | ESN | 0.681 | 1 | 0.00059 |
| 99 | *OR2T10* | 1 | 248.590 | CEU | 0.683 | 1 | 0.015 |
| 100 | *GGN* | 19 | 38.384 | GWD | 0.686 | 3 | 0.0039 |
| 101 | *BAG2* | 6 | 57.172 | STU | 0.688 | 3 | 0.0032 |
| 102 | *ABHD17B* | 9 | 71.862 | ITU | 0.688 | 1 | 0.0049 |
| 103 | *NFE2* | 12 | 54.292 | LWK | 0.692 | 2 | 0.0015 |
| 104 | *TUT1* | 11 | 62.575 | MSL | 0.692 | 2 | 0.002 |
| 105 | *SET* | 9 | 128.683 | CLM | 0.692 | 1 | 0.0052 |
| 106 | *EPB42* | 15 | 43.197 | FIN | 0.696 | 1 | 0.0053 |
| 107 | *HGH1* | 8 | 144.138 | PJL | 0.698 | 4 | 0.0032 |
| 108 | *ARL5A* | 2 | 151.789 | PEL | 0.705 | 1 | 0.0074 |
| 109 | *BBX* | 3 | 107.523 | CHS | 0.716 | 1 | 0.021 |
| 110 | *PLK4* | 4 | 127.881 | LWK | 0.723 | 1 | 0.002 |
| 111 | *CDC6* | 17 | 40.288 | GIH | 0.726 | 2 | 0.0053 |
| 112 | *ZFTA* | 11 | 63.760 | PEL | 0.737 | 1 | 0.0032 |
| 113 | *FGF8* | 10 | 101.770 | MSL | 0.739 | 1 | 0.00091 |

*Continued*

| # | Representative | Chr | Start (Mb) | Pop | Rank (%) | $n$ | Score |
|---|---|---|---|---|---|---|---|
| 114 | *MRPL46* | 15 | 88.459 | CLM | 0.740 | 2 | 0.0031 |
| 115 | *TOMM20L* | 14 | 58.396 | LWK | 0.750 | 1 | 0.0015 |
| 116 | *ABCE1* | 4 | 145.098 | ASW | 0.750 | 1 | 0.0011 |
| 117 | *BCLAF1* | 6 | 136.257 | MXL | 0.758 | 2 | 0.0033 |
| 118 | *SLC35A3* | 1 | 99.969 | PEL | 0.769 | 2 | 0.014 |
| 119 | *SRSF11* | 1 | 70.206 | ASW | 0.772 | 2 | 0.0052 |
| 120 | *BCL7B* | 7 | 73.536 | LWK | 0.781 | 1 | 0.0017 |
| 121 | *C9orf43* | 9 | 113.410 | ASW | 0.782 | 1 | 0.0024 |
| 122 | *SLC25A36* | 3 | 140.942 | BEB | 0.794 | 1 | 0.0067 |
| 123 | *HVCN1* | 12 | 110.628 | CHS | 0.816 | 3 | 0.01 |
| 124 | *PSMA8* | 18 | 26.134 | ITU | 0.820 | 1 | 0.035 |
| 125 | *TCHHL1* | 1 | 152.084 | ASW | 0.824 | 1 | 0.00084 |
| 126 | *NSD2* | 4 | 1.871 | CLM | 0.835 | 1 | 0.0046 |
| 127 | *DHFR2* | 3 | 94.048 | MSL | 0.840 | 1 | 0.0023 |
| 128 | *DEFB112* | 6 | 50.042 | JPT | 0.841 | 1 | 0.0038 |
| 129 | *STRAP* | 12 | 15.882 | ITU | 0.842 | 2 | 0.026 |
| 130 | *OGN* | 9 | 92.383 | GWD | 0.844 | 1 | 0.006 |
| 131 | *PPP1R12A* | 12 | 79.774 | GBR | 0.850 | 1 | 0.0071 |
| 132 | *RAB43* | 3 | 129.088 | FIN | 0.861 | 2 | 0.0054 |
| 133 | *OCLN* | 5 | 69.492 | PJL | 0.862 | 1 | 0.0094 |
| 134 | *MAMDC4* | 9 | 136.851 | GWD | 0.866 | 1 | 0.015 |
| 135 | *MYD88* | 3 | 38.139 | PJL | 0.904 | 1 | 0.0069 |
| 136 | *GSTCD* | 4 | 105.709 | KHV | 0.913 | 1 | 0.0081 |
| 137 | *ENTHD1* | 22 | 39.743 | YRI | 0.914 | 1 | 0.0073 |
| 138 | *GEN1* | 2 | 17.754 | CDX | 0.937 | 2 | 0.023 |
| 139 | *EPHX3* | 19 | 15.227 | ASW | 0.941 | 1 | 0.0018 |
| 140 | *LRRC42* | 1 | 53.946 | ITU | 0.947 | 1 | 0.014 |
| 141 | *EDC3* | 15 | 74.631 | CEU | 0.985 | 1 | 0.0049 |
| 142 | *AVIL* | 12 | 57.797 | GWD | 0.992 | 1 | 0.009 |
| 143 | *YIPF2* | 19 | 10.922 | GWD | 0.997 | 1 | 0.0062 |

Table S3: **Membership of the 82 multi-gene clusters in the reference screen.** Only genes passing stages 1–4 enter these single-linkage clusters; the remaining 61 loci are singletons. Span is the distance from the start of the first gene to the end of the last gene in each cluster.

| Representative | Chr | $n$ | Span (Mb) | Stage-4 members |
|---|---|---|---|---|
| *SHCBP1* | 16 | 9 | 0.886 | SHCBP1, VPS35, ORC6, MYLK3, C16orf87, GPT2, DNAJA2, NETO2, ITFG1 |
| *CTF1* | 16 | 26 | 2.139 | ZG16, KIF22, MAZ, PRRT2, ENSG00000280893, PAGR1, ENSG00000281348, MVP, ZNF688, SRCAP, ENSG00000282034, TMEM265, PHKG2, CFAP119, RNF40, ZNF629, BCL7C, CTF1, FBXL19, ORAI3, SETD1A, HSD3B7, STX1B, PRSS8, PRSS36, ZNF267 |
| *SLC9A5* | 16 | 41 | 1.209 | CBFB, PHAF1, B3GNT9, TRADD, FBXL8, ENSG00000265690, HSF4, NOL3, EXOC3L1, E2F4, ELMO3, TMEM208, FHOD1, SLC9A5, PLEKHG4, KCTD19, LRRC36, TPPP3, ZDHHC1, HSD11B2, ATP6V0D1, AGRP, RIPOR1, CTCF, CARMIL2, ACD, PARD6A, ENKD1, C16orf86, GFOD2, RANBP10, TSNAXIP1, CENPT, NUTF2, EDC4, NRN1L, PSKH1, CTRL, ENSG00000261884, LCAT, ESRP2 |
| *EXOC6B* | 2 | 3 | 0.760 | EXOC6B, SPR, EMX1 |
| *DCAF4L1* | 4 | 4 | 0.218 | TMEM33, DCAF4L1, SLC30A9, BEND4 |

*Continued*

| Representative | Chr | $n$ | Span (Mb) | Stage-4 members |
|---|---|---|---|---|
| *P4HTM* | 3 | 49 | 3.906 | PFKFB4, UCN2, COL7A1, UQCRC1, TMEM89, SLC26A6, CELSR3, NCKIPSD, PRKAR2A, SLC25A20, ARIH2, P4HTM, WDR6, DALRD3, NDUFAF3, IMPDH2, ENSG00000290315, QRICH1, QARS1, USP19, LAMB2, CCDC71, KLHDC8B, C3orf84, IHO1, USP4, APEH, SEMA3B, LSMEM2, IFRD2, TUSC2, NPRL2, CYB561D2, ENSG00000272104, TMEM115, CISH, MANF, RBM15B, RAD54L2, TEX264, GRM2, ENSG00000285749, RRP9, PARP3, GPR62, PCBP4, RPL29, BAP1, PHF7 |
| *BCL2L1* | 20 | 3 | 0.210 | HM13, COX4I2, BCL2L1 |
| *PPM1D* | 17 | 6 | 0.625 | HEATR6, CA4, USP32, CHCT1, APPBP2, PPM1D |
| *BMI1* | 10 | 4 | 0.138 | COMMD3, COMMD3-BMI1, BMI1, SPAG6 |
| *GRK2* | 11 | 16 | 0.835 | CCDC87, CCS, RBM14, SPTBN2, TOP6BL, KDM2A, GRK2, ANKRD13D, SSH3, RAD9A, POLD4, ENSG00000256514, CLCF1, PPP1CA, TBC1D10C, CARNS1 |
| *PLA2G10* | 16 | 3 | 0.259 | PARN, BFAR, PLA2G10 |
| *PSMB2* | 1 | 17 | 1.120 | ENSG00000284773, TMEM35B, ENSG00000271741, ZMYM6, ZMYM1, SFPQ, ZMYM4, KIAA0319L, NCDN, TFAP2E, PSMB2, C1orf216, AGO4, AGO1, AGO3, TEKT2, ADPRS |
| *CENPW* | 6 | 3 | 0.392 | HINT3, TRMT11, CENPW |
| *PPP3CB* | 10 | 13 | 0.733 | OIT3, PLA2G12B, P4HA1, NUDT13, ECD, FAM149B1, DNAJC9, MRPS16, CFAP70, ANXA7, MSS51, PPP3CB, USP54 |
| *ORC2* | 2 | 3 | 0.182 | NIF3L1, ORC2, HYCC2 |
| *PIGV* | 1 | 11 | 1.158 | ARID1A, PIGV, ZDHHC18, GPN2, GPATCH3, NUDC, NR0B2, TENT5B, FAM76A, STX12, PPP1R8 |
| *TSSK3* | 1 | 5 | 0.205 | HDAC1, TSSK3, FAM229A, BSDC1, ZBTB8B |
| *FBXO4* | 5 | 3 | 0.212 | OXCT1, RIMOC1, FBXO4 |
| *PRMT9* | 4 | 2 | 0.067 | TMEM184C, PRMT9 |
| *VPS37C* | 11 | 3 | 0.193 | CD5, VPS37C, VWCE |
| *VDAC3* | 8 | 8 | 0.806 | IKBKB, POLB, DKK4, VDAC3, THAP1, RNF170, HOOK3, ENSG00000254673 |
| *PSME3* | 17 | 9 | 0.669 | GHDC, VPS25, WNK4, COA3, CNTD1, BECN1, PSME3, AOC2, AOC3 |
| *FMNL3* | 12 | 7 | 0.398 | SPATS2, KCNH3, MCRS1, PRPF40B, FAM186B, FMNL3, TMBIM6 |
| *TMEM145* | 19 | 19 | 0.471 | RABAC1, ENSG00000285505, ATP1A3, GRIK5, ZNF574, POU2F2, DEDD2, ENSG00000288671, ZNF526, GSK3A, ENSG00000268643, ERF, CIC, PAFAH1B3, PRR19, TMEM145, MEGF8, CNFN, LIPE |
| *DOK1* | 2 | 5 | 0.122 | AUP1, HTRA2, LOXL3, DOK1, M1AP |
| *ERMN* | 2 | 2 | 0.070 | GALNT5, ERMN |
| *ZNF546* | 19 | 3 | 0.107 | ZNF546, ZNF780B, ZNF780A |
| *PLPP5* | 8 | 7 | 0.714 | ZNF703, ERLIN2, BAG4, DDHD2, PLPP5, NSD3, LETM2 |
| *ZNF510* | 9 | 2 | 0.139 | PRXL2C, ZNF510 |
| *RNF11* | 1 | 6 | 1.185 | RNF11, EPS15, TXNDC12, ENSG00000285839, KTI12, PRPF38A |
| *AMBRA1* | 11 | 4 | 0.291 | CHRM4, AMBRA1, HARBI1, ATG13 |
| *OTX1* | 2 | 2 | 0.384 | EHBP1, OTX1 |
| *KIF23* | 15 | 2 | 0.042 | KIF23, RPLP1 |
| *KMT2A* | 11 | 3 | 0.130 | ENSG00000285827, KMT2A, TTC36 |
| *GOLM2* | 15 | 3 | 0.375 | GOLM2, EIF3J, SPG11 |
| *GLMN* | 1 | 5 | 0.407 | BTBD8, C1orf146, GLMN, RPAP2, GFI1 |
| *SGCB* | 4 | 4 | 0.758 | LRRC66, SGCB, USP46, ERVMER34-1 |
| *IFIH1* | 2 | 3 | 0.201 | FAP, IFIH1, GCA |
| *ALG2* | 9 | 2 | 0.014 | ALG2, SEC61B |

*Continued*

| Representative | Chr | $n$ | Span (Mb) | Stage-4 members |
|---|---|---|---|---|
| *PPP1R42* | 8 | 6 | 0.494 | MYBL1, C8orf44-SGK3, SGK3, MCMDC2, TCF24, PPP1R42 |
| *MRPS5* | 2 | 12 | 1.832 | MAL, MRPS5, ENSG00000289685, ZNF514, ZNF2, ZNF892, PROM2, STARD7, TMEM127, CIAO1, SNRNP200, ANKRD39 |
| *SYF2* | 1 | 2 | 0.140 | SYF2, TMEM50A |
| *APOF* | 12 | 8 | 0.205 | MYL6, SMARCC2, ANKRD52, COQ10A, CNPY2, IL23A, STAT2, APOF |
| *C2orf66* | 2 | 3 | 0.165 | GTF3C3, C2orf66, PGAP1 |
| *PCYT2* | 17 | 6 | 0.040 | ALYREF, ANAPC11, NPB, PCYT2, SIRT7, MAFG |
| *PYGO2* | 1 | 5 | 0.037 | PYGO2, SHC1, CKS1B, FLAD1, LENEP |
| *TRIR* | 19 | 2 | 0.018 | TRIR, GET3 |
| *ENSG00000266086* | 17 | 3 | 0.036 | VEZF1, ENSG00000266086, SRSF1 |
| *CELF6* | 15 | 5 | 0.142 | ENSG00000273025, CELF6, ENSG00000260729, HEXA, TMEM202 |
| *ZHX1* | 8 | 3 | 0.056 | C8orf76, ZHX1-C8orf76, ZHX1 |
| *PCLAF* | 15 | 5 | 0.821 | PCLAF, TRIP4, UBAP1L, PDCD7, CLPX |
| *RIPPLY2* | 6 | 2 | 0.114 | RIPPLY2, CYB5R4 |
| *NIT1* | 1 | 8 | 0.107 | NECTIN4, KLHDC9, NIT1, DEDD, UFC1, USP21, PPOX, B4GALT3 |
| *PROCA1* | 17 | 8 | 0.683 | NLK, IFT20, ALDOC, SPAG5, ENSG00000258472, PROCA1, RAB34, RPL23A |
| *GTSE1* | 22 | 4 | 0.087 | CDPF1, PKDREJ, TTC38, GTSE1 |
| *PDZD9* | 16 | 3 | 0.362 | IGSF6, UQCRC2, PDZD9 |
| *FKBP3* | 14 | 5 | 0.238 | C14orf28, KLHL28, TOGARAM1, PRPF39, FKBP3 |
| *CYP3A5* | 7 | 4 | 0.866 | TMEM225B, ZSCAN25, CYP3A5, SPACDR |
| *MCM4* | 8 | 3 | 1.149 | PRKDC, MCM4, SNAI2 |
| *IGIP* | 5 | 8 | 0.856 | MATR3, MZB1, PROB1, SPATA24, ENSG00000279686, ECSCR, PURA, IGIP |
| *GRSF1* | 4 | 2 | 0.136 | RUFY3, GRSF1 |
| *MRPL16* | 11 | 3 | 0.209 | PATL1, MRPL16, CBLIF |
| *H2BC21* | 1 | 3 | 0.261 | H2BC21, BOLA1, VPS45 |
| *C6orf52* | 6 | 2 | 0.038 | C6orf52, PAK1IP1 |
| *FEV* | 2 | 2 | 0.012 | FEV, CRYBA2 |
| *ZNF660* | 3 | 3 | 0.093 | ZKSCAN7, ZNF660, ZNF197 |
| *OR1L4* | 9 | 2 | 0.027 | OR1L4, OR1L6 |
| *CEP95* | 17 | 4 | 1.456 | TACO1, CEP95, SMURF2, GNA13 |
| *GGN* | 19 | 3 | 0.025 | PSMD8, GGN, SPRED3 |
| *BAG2* | 6 | 3 | 0.267 | BEND6, BAG2, RAB23 |
| *NFE2* | 12 | 2 | 0.800 | TARBP2, NFE2 |
| *TUT1* | 11 | 2 | 0.032 | ENSG00000255508, TUT1 |
| *HGH1* | 8 | 4 | 0.062 | ENSG00000290230, EXOSC4, GPAA1, HGH1 |
| *CDC6* | 17 | 2 | 0.130 | CDC6, TOP2A |
| *MRPL46* | 15 | 2 | 0.021 | MRPL46, MRPS11 |
| *BCLAF1* | 6 | 2 | 0.059 | MTFR2, BCLAF1 |
| *SLC35A3* | 1 | 2 | 0.114 | SLC35A3, ENSG00000283761 |
| *SRSF11* | 1 | 2 | 0.108 | LRRC40, SRSF11 |
| *HVCN1* | 12 | 3 | 0.214 | VPS29, TCTN1, HVCN1 |
| *STRAP* | 12 | 2 | 0.283 | EPS8, STRAP |
| *RAB43* | 3 | 2 | 0.062 | GP9, RAB43 |
| *GEN1* | 2 | 2 | 0.278 | RAD51AP2, GEN1 |

#### 2.3 Garud's $H_{12}$ and XP-EHH at *GRK2* and the four East/South-Asian additional examples

Table S4: **Haplotype statistics at five focal genes and five comparison genes.** We compare *GRK2*, *CCDC92*, *SLC6A15*, *BPIFA2* and *CLEC6A* with five other genes using Garud's $H_{12}$, $H_2/H_1$ and XP-EHH. We calculate $H_{12}$ in 400-site windows, moving 50 sites at a time within ±500 kb of each gene midpoint, including sites monomorphic in the focal population. We take the highest $H_{12}$ value in this region and measure $H_2/H_1$ in the same window. Both are reported as percentiles relative to other genes analysed in the same population. Mean XP-EHH compares pooled haplotypes from the focal population's superpopulation (SAS or EAS) with YRI across each gene and its ±500 kb flanks. The $H_{12}$ results for *TREML1*/*TREM2* appear in Fig. 3j.

| Gene | Focal pop | $H_{12}$ percentile | $H_2/H_1$ percentile | Mean XP-EHH |
|---|---|---|---|---|
| *GRK2 and four of the five additional examples* | | | | |
| *CCDC92* | CDX | 91.3 | 7.6 | 1.56 |
| *SLC6A15* | CHS | 99.5 | 5.9 | 0.86 |
| *GRK2* | GIH | 94.5 | 1.9 | 1.28 |
| *BPIFA2* | GIH | 97.6 | 92.4 | 0.95 |
| *CLEC6A* | CDX | 87.2 | 3.2 | 1.13 |
| *Comparison genes* | | | | |
| *ADAM22* | CDX | 92.3 | 98.1 | 0.93 |
| *CCDC70* | GIH | 77.5 | 62.2 | 0.40 |
| *TMEM30A* | CDX | 63.1 | 55.6 | 0.73 |
| *ZNF420* | CDX | 66.5 | 14.0 | 1.28 |
| *RAB11FIP3* | GIH | 80.8 | 5.9 | 0.95 |

#### 2.4 Top 50 SD-masked genome-wide candidates

Table S5: **Top 50 genes by minimum TMRCA rank after unique-base segmental-duplication masking.** Percentiles are from the TMRCA gene list. This ranking precedes continental replication, reference-locus exclusion and clustering.

| # | Gene | Chr | Pop | Min rank (%) | Max rank (%) |
|---|---|---|---|---|---|
| 1 | *SHCBP1* | 16 | PEL | 0.027 | 0.64 |
| 2 | *SLC24A5* | 15 | TSI | 0.037 | 22.64 |
| 3 | *MYEF2* | 15 | FIN | 0.043 | 14.66 |
| 4 | *CTXN2* | 15 | TSI | 0.053 | 26.76 |
| 5 | *VPS35* | 16 | PEL | 0.080 | 3.04 |
| 6 | *OR4C6* | 11 | LWK | 0.084 | 0.47 |
| 7 | *CTF1* | 16 | CHB | 0.101 | 19.90 |
| 8 | *ORAI3* | 16 | CHB | 0.112 | 8.39 |
| 9 | *NETO2* | 16 | PEL | 0.122 | 4.49 |
| 10 | *SLC9A5* | 16 | JPT | 0.122 | 9.55 |
| 11 | *ZRANB3* | 2 | CEU | 0.127 | 42.82 |
| 12 | *EXOC6B* | 2 | JPT | 0.128 | 17.56 |
| 13 | *FBXL19* | 16 | CHB | 0.138 | 31.43 |
| 14 | *ABCC11* | 16 | CHB | 0.143 | 29.24 |
| 15 | *DCAF4L1* | 4 | CHB | 0.149 | 20.72 |
| 16 | *P4HTM* | 3 | CDX | 0.149 | 11.12 |
| 17 | *RANBP10* | 16 | CHB | 0.154 | 25.18 |
| 18 | *TMEM33* | 4 | JPT | 0.154 | 34.32 |
| 19 | *DALRD3* | 3 | CDX | 0.154 | 6.42 |
| 20 | *ORC6* | 16 | PEL | 0.159 | 4.75 |
| 21 | *BCL2L1* | 20 | JPT | 0.160 | 21.45 |
| 22 | *GPT2* | 16 | PEL | 0.164 | 6.43 |
| 23 | *TMEM208* | 16 | CHB | 0.165 | 17.06 |
| 24 | *WDR6* | 3 | CDX | 0.165 | 6.32 |
| 25 | *PPM1D* | 17 | GBR | 0.170 | 26.82 |
| 26 | *FHOD1* | 16 | CHB | 0.170 | 14.27 |
| 27 | *NDUFAF3* | 3 | CDX | 0.170 | 5.40 |
| 28 | *CFAP119* | 16 | JPT | 0.176 | 2.80 |
| 29 | *PHKB* | 16 | PEL | 0.180 | 2.07 |
| 30 | *BCL7C* | 16 | CHB | 0.181 | 22.33 |
| 31 | *BMI1* | 10 | FIN | 0.181 | 3.77 |
| 32 | *R3HDM1* | 2 | CEU | 0.185 | 42.66 |
| 33 | *DPH5* | 1 | PEL | 0.186 | 17.43 |
| 34 | *IMPDH2* | 3 | CDX | 0.186 | 5.62 |
| 35 | *GRK2* | 11 | GIH | 0.191 | 19.39 |
| 36 | *LONP2* | 16 | CHB | 0.191 | 11.78 |
| 37 | *ARIH2* | 3 | CDX | 0.192 | 15.21 |
| 38 | *DARS1* | 2 | CEU | 0.196 | 59.97 |
| 39 | *ITFG1* | 16 | PEL | 0.196 | 3.64 |
| 40 | *SLC30A9* | 4 | CHB | 0.197 | 27.57 |
| 41 | *COMMD3-BMI1* | 10 | FIN | 0.197 | 6.13 |
| 42 | *COX4I2* | 20 | JPT | 0.197 | 51.16 |
| 43 | *PLA2G10* | 16 | LWK | 0.201 | 3.18 |
| 44 | *RAB3GAP1* | 2 | CEU | 0.207 | 38.07 |
| 45 | *PSMB2* | 1 | TSI | 0.207 | 34.90 |
| 46 | *AGRP* | 16 | KHV | 0.212 | 16.36 |
| 47 | *CYB561A3* | 11 | FIN | 0.213 | 23.16 |
| 48 | *CENPW* | 6 | JPT | 0.213 | 14.90 |
| 49 | *RNF40* | 16 | JPT | 0.218 | 4.32 |
| 50 | *COMMD3* | 10 | GIH | 0.223 | 11.90 |

#### 2.5 Prior-literature comparison at the *TREML1*/*TREM2* cluster (chr6p21.1)

We compare the *TREML1*/*TREM2* coalescence pattern outside Africa with previous selection studies. Across non-African populations, *TREM2* TMRCA ranks range from 1.07–5.95%, above our 1% cutoff. Table S6 lists the 52 resources we examined, with their methods, study populations and findings at this locus. We searched supplementary tables and genomic coordinates where available, highlighting separately whether the locus was absent from the literature checked.

The resources use different approaches to study the evolutionary history and biological roles of this region. We group their findings by the question studied, distinguishing positive selection from balancing selection, purifying selection, introgression and disease association. The table also identifies reviews and studies focused on individual loci.

For example, `Flex-Sweep` is reporting selection signal at *TREML1* in YRI, with a score of 0.818 above its default self-defined threshold of 0.5 [2]. Notably, the reported region also overlaps approximately 1.6 kb of *TREM2*, although only *TREML1* is named in the table. This shows why we checked genomic coordinates as well as gene names. Our own iHS analysis ranks genes by the fraction of sites with extreme scores within each gene. It places *TREM2* among the top 0.4% in BEB, STU and ITU (Table 3).

Table S6: **Prior-literature comparison for positive selection at the *TREML1*/*TREM2* cluster (chr6:41,116,895–41,130,924 GRCh37; chr6p21.1).** Fifty-two literature resources examined (source details in the repository): published catalogs, supplementary tables, released raw statistics, ML-based selection methods, ancient-DNA scans, archaic-introgression catalogs, immune-gene-focused reviews, regional cohort scans, and disease-genetics literature. "No" means no positive-selection call identified in the inspected material, not necessarily a tested, sub-threshold result. Locus-specific demonstrations and non-selection studies are labelled separately; balancing-selection, purifying-selection and introgression evidence are distinguished from positive-selection results. "chr6:41 Mb window hit" is a coordinate-level grep over the chr6:41,000,000–41,300,000 GRCh37 interval (or GRCh38 equivalent) against available per-position output inspected; checks limited to named candidate lists are labelled as such (BED, BigWig, supplementary table). Window-based iHS summaries differ from our measure of the fraction of sites with extreme iHS scores within each gene, which recovers South Asian signals in the top 0.4%. `Flex-Sweep` annotates *TREML1* in YRI and its interval partially overlaps *TREM2* (row 18), despite lacking a *TREM2* gene-name annotation.

| # | Resource | Year | Method | Cohort | TREM2 / TREML1 hit | chr6:41 Mb window / note |
|---|---|---|---|---|---|---|
| 1 | Voight *et al.* [1] | 2006 | iHS | HapMap I | no | not in landmark top-region list |
| 2 | Sabeti *et al.* [3] | 2007 | LRH + XP-EHH | HapMap 2 | no | published top-22 list excludes TREM cluster; supplementary Table S10 has rs9349180 (Sabeti hg17 chr6:41,293,452 = GRCh37 chr6:41,185,474, CHB+JPT) inside the *TREML3P* pseudogene, ∼55 kb downstream of TREM2 focal block (*not* at TREML1/TREM2) |
| 3 | Pickrell *et al.* [4] | 2009 | iHS / XP-EHH | HGDP | published-list check | Candidate genes in supplementary PDF; unavailable browser not treated as a negative test |
| 4 | Grossman *et al.* [5] | 2013 | CMS composite | various | no focal-gene annotation | Original supplementary Tables S2–S10: neither focal gene named; complete raw CMS statistics not reanalysed |
| 5 | Field *et al.* SDS [6] | 2016 | SDS | UK10K (n=3195) | TREM2: 0 sites; TREML1: 4 | Original UK10K release: TREML1 max \|SDS\|=0.390; chr6:41.10–41.20 Mb has 52 sites, max 2.088 (96.05th absolute-score percentile) |

*(continued on next page)*

*(continued from previous page)*

| # | Resource | Year | Method | Cohort | TREM2 / TREML1 hit | chr6:41 Mb window / note |
|---|---|---|---|---|---|---|
| 6 | Johnson & Voight [7] | 2018 | iHS | 26 1000G panels | no interval overlap | Original Supplementary Table 3: no candidate interval overlaps either focal gene; nearby intervals occur in African panels |
| 7 | `PopHumanScan` [8] | 2019 | 8 separate statistics | 22 non-admixed 1000G populations | no | 268-publication annotation resource; 80 Bitarello NCD2 rows in chr6:41.0–41.3 Mb represent 11 distinct intervals; none overlaps the focal gene bodies |
| 8 | Bitarello *et al.* [9] | 2018 | NCD2 balancing | 1000G | no | Table S5: 11 distinct 3-kb intervals, some overlapping, spanning 41,200,605–41,299,605 GRCh37, ∼70–169 kb beyond *TREM2*; no focal-gene overlap |
| 9 | Speidel *et al.* `Relate` [10] | 2019 | ARG-based | 1000G | absent from reported list | Original Supplementary Table 3: no focal-gene annotation or chr6:41.0–41.3 Mb candidate region |
| 10 | Souilmi *et al.* [11] | 2021 | VIP / coronavirus | various | not in CoV-VIP set | Neither focal gene is in the 420-gene CoV-VIP set or the selected examples in original Table S1 |
| 11 | Kerner *et al.* [12] | 2023 | aBC (genome-wide) | 2879 anc. + mod. EUR | absent from 89-lead list | Original Table S2: no lead at chr6:41.0–41.3 Mb; 89 is the number of candidate loci, not a restricted immune-gene panel |
| 12 | Irving-Pease *et al.* [13] | 2024 | ancestry-`CLUES` | 1015 after QC | no scored focal-gene variant | Original Tables S2.1.3/S2.1.4 contain five positions in chr6:41.0–41.3 Mb, none in either focal gene; minimum regional $P \approx 3 \times 10^{-6}$ (EHG), no listed sweep |
| 13 | Le *et al.* [14] | 2022 | admixture-aware aDNA | 1291 ancient Europeans | published-list check | 25 reported candidate loci; no focal-gene candidate identified in the inspected main text and Table 1 |
| 14 | Garcia-Calleja *et al.* GCAT [15] | 2025 | Iberian-specific | 704 IBR | no focal-gene annotation | Original supplementary workbook: neither focal gene named in the inspected candidate lists; no gene-specific null inferred |
| 15 | Akbari *et al.* [16] | 2026 | aDNA time-series | 15,836 ancient + 6,438 modern W. Eurasians | no | PASS maxima: TREM2 POSTERIOR=0.0698, TREML1=0.0099; 1045 PASS variants in 41.0–41.3 Mb, max=0.603. Including warnings: 1553 variants; TREML1 max=0.1047 |
| 16 | Maravall-López *et al.* [17] | 2026 | aDNA time-series (LMM) | Akbari-source data | below genome-wide threshold | Released LMM statistics: TREML1 13 variants, min $P = 0.116$; TREM2 4, min $P = 0.400$; chr6:41.0–41.3 Mb 1553, min $P = 0.0031$. Shared Akbari source data |
| 17 | Zhao *et al.* `HaploSweep` [18] | 2024 | haplotype clustering | 1000G | no listed soft-sweep overlap | Original Tables S12–S14: no candidate gene interval at chr6:41.0–41.3 Mb in CHB, CEU or YRI |

*(continued on next page)*

*(continued from previous page)*

| # | Resource | Year | Method | Cohort | TREM2 / TREML1 hit | chr6:41 Mb window / note |
|---|---|---|---|---|---|---|
| 18 | Lauterbur *et al.* `Flex-Sweep` [2] | 2023 | flex sweep | 1000G YRI | TREML1 annotated; partial TREM2 overlap | 260 kb streak chr6:40,867,841–41,127,841 (GRCh37, YRI demographic): 5 upstream genes above strict 0.99 threshold (*APOBEC2*, *OARD1*, *TSPO2*, *UNC5CL*, *NFYA*); *TREML1* at score 0.818 (above default 0.5 threshold, below strict 0.99); streak overlaps ∼1.6 kb of *TREM2*, which lacks a gene-name row |
| 19 | Hejase *et al.* `SIA` [19] | 2022 | ARG + deep learning | CEU | not assessed | selected phenotype-associated loci; no genome-wide TREM test |
| 20 | Refoyo-Martínez *et al.* `GRoSS` [20] | 2019 | graph-based allele-frequency test | various | absent from top lists | Original Tables S2/S3: no focal-gene annotation or chr6:41.0–41.3 Mb candidate region |
| 21 | Akbari *et al.* `iSAFE` [21] | 2018 | fine-mapper | 22 known sweeps | not assessed | not in test loci |
| 22 | `FineMAV` (Szpak *et al.*) [22] | 2018 | variant-level | 1000G Phase 3 | no top-100 overlap | Original Additional file 2: no top-100 variant at chr6:41.0–41.3 Mb in AFR, EAS, EUR or EAS+EUR |
| 23 | Luo *et al.* *Sci. Bull.* [23] | 2023 | Han Chinese SDS | 3946 Han Chinese | not resolved per variant | Abstract reports 24 selected loci, including MHC, ALDH2 and ADH1B; complete TREM-region statistics not independently checked |
| 24 | Pybus *et al.* 1000G Browser [24] | 2014 | 11-statistic aggregator | 1000G P1 | not re-queried | Aggregator; live browser unavailable for direct query |
| 25 | Metspalu *et al.* [25] | 2011 | iHS South Asia | various | absent from top-20 lists | Original Tables S5/S6: neither focal gene among the top-20 Indian iHS or XP-EHH regions |
| 26 | Quintana-Murci *Cell* review [26] | 2019 | immune-selection review | – | review; no new locus test | Context for immune adaptation; omission is not a negative test |
| 27 | Barreiro & Quintana-Murci *Hum. Genet.* [27] | 2020 | immune review | – | review; no new locus test | Context for immune adaptation; omission is not a negative test |
| 28 | Quach *et al.* *Cell* [28] | 2016 | immune + Neanderthal | various | no focal-gene candidate annotation | Original Tables S4B/S4C: neither focal gene in FST/iHS candidate lists; expression-panel membership is a separate result |
| 29 | Deschamps *et al.* *AJHG* [29] | 2016 | innate-immunity FCS | 1000G | no | innate-immunity-restricted FCS scan; all 5 TREM-cluster genes in the 1,553-gene panel (Table S1, "sensor" class; TREM2 `SnIPRE` $f$=0.688, panel median 0.447) but none among the 57 positive-selection targets (Table 2) — panel inclusion and candidate-list absence, not a gene-specific null-test result |
| 30 | Barreiro & Quintana-Murci *NRG* [30] | 2010 | foundational immune review | – | review; no new locus test | Context for immune adaptation; omission is not a negative test |
| 31 | Nandakumar *et al.* *MBE* [31] | 2025 | mammalian immune | comparative | not a human sweep test | Rodent comparative codon-selection analysis; does not test recent selection at the human TREM locus |
| 32 | Patin & Quintana-Murci *Annu. Rev. Immunol.* [32] | 2025 | review | – | review; no new locus test | Context for ancient-DNA studies; omission is not a negative test |
| 33 | Fumagalli & Sironi *Curr. Opin. Immunol.* [33] | 2014 | immune-selection review | – | review; no new locus test | Immune-selection review; omission is not a negative test |

*(continued on next page)*

*(continued from previous page)*

| # | Resource | Year | Method | Cohort | TREM2 / TREML1 hit | chr6:41 Mb window / note |
|---|---|---|---|---|---|---|
| 34 | Gurdasani *et al.* AGVP *Nature* [34] | 2015 | iHS / differentiation | 1481 genotyped; 320 WGS | no focal-gene annotation | Original Supplementary Note 8, Tables 1–4: neither focal gene named in the reported iHS candidates |
| 35 | Choudhury *et al. Nature* [35] | 2020 | high-depth Africa | various | no focal-gene annotation | Original selection Tables S6–S14 contain neither focal gene name; complete variant statistics not reanalysed |
| 36 | Lindo *et al. Sci. Adv.* [36] | 2018 | ancient + modern Andean | Peru/ Bolivia/ Chile | absent from top lists | Original Tables S4/S5 list Andean PBS candidates without either focal gene |
| 37 | Vernot & Akey *Science* [37] | 2014 | Neanderthal intro. | various | not resolved | Introgression study; a complete focal-region candidate-list check was not established |
| 38 | Racimo *et al. MBE* [38] | 2017 | adaptive intro. | various | no listed interval overlap | Original Table S3 (2,041 rows): no candidate interval at chr6:41.0–41.3 Mb |
| 39 | Browning *et al.* `Sprime` [39] | 2018 | introgression | various | introgression catalog | Archaic segment inference alone is not a positive-selection test |
| 40 | Skov *et al.* [40] | 2020 | Neand. intro. Iceland | 27566 IS | no positive-selection claim (introgression catalog) | Supplementary dataset 1: chr6:40,971,209–41,154,236 *GRCh38*, 183,027 bp, 51 DAV-SNPs, frequency count 15. Overlaps most of *TREML1*; ends 4,270 bp before *TREM2*. Relationship to our signal unresolved |
| 41 | Villanea *et al. Science* (MUC19) [41] | 2025 | MUC19 introgression + selection | various | not assessed | locus-specific MUC19 study |
| 42 | Zeberg & Pääbo [42, 43] | 2020/21 | COVID Neand. intro. | various | not assessed at TREM | Locus-specific COVID-associated introgression at chr3 and OAS on chr12 |
| 43 | Dannemann *et al.* [44] | 2016 | introgression | various | not assessed at TREM | Adaptive-introgression study of TLR6–TLR1–TLR10; no TREM-locus test |
| 44 | Simonti *et al.* [45] | 2016 | Neand. + EHR phenotype | various | not selection test | phenotype associations with introgressed alleles |
| 45 | van den Belt & Alachiotis `FASTER-NN` [46] | 2025 | CNN purifying (negative) selection | 1000G CEU | n/a (purifying scan, not a positive-selection scan) | The method is general but the paper's only human-genome application is a purifying-selection scan in CEU; consequently the paper does not produce a positive-selection candidate-region list |
| 46 | Salazar-Tortosa *et al.* MDR [47] | 2023 | MDR iHS | 5×1000G | not a locus-level comparison | Mixture-regression analysis of genomic iHS patterns; inspected supplement does not establish a focal-gene selection test |
| 47 | Colbran *et al.* bioRxiv [48] | 2026 | aDNA 7244 individuals | 5 continental regions | absent from 31-locus list | Original Supplementary Table 1 lists significant loci, without TREML1/TREM2; no gene-specific null result inferred |
| 48 | Barton *et al.* "Convergent" bioRxiv [49] | 2026 | aDNA E. + W. Eurasia | various | not resolved | Abstract reports 40 signals in 1,862 ancient East Eurasians; full candidate statistics not independently checked |
| 49 | `dbPSHP` HKU [50] | 2014 | aggregator | HM3 + 1KG | not re-queried | Live database unavailable for direct query |
| 50 | Sims *et al. Nat. Genet.* [51] | 2017 | rare-coding GWAS | AD case/control | risk variant only | R62H AD-risk association; R47H also evaluated; no selection test |
| 51 | Bellenguez *et al. Nat. Genet.* [52] | 2022 | GWAS meta-analysis | AD case/control | risk variant only | no selection claim |
| 52 | Carrasquillo *et al. Alz. & Demen.* [53] | 2017 | eQTL + AD | brain | not selection | regulatory variant rs9357347-C |

### 3 Extended Case Studies

We provide supporting details for the two focal loci in Fig. 3, followed by five additional case studies. Figure S8 summarises their population ranks, ancient-DNA selection statistics, variant summaries and literature evidence, with *LCT* as a positive control. Table 3 reports the complementary genetic evidence.

#### 3.1 *GRK2/ADRBK1*: further supporting lines of evidences from haplotype to literature

*GRK2* (also *ADRBK1*; G protein-coupled receptor kinase 2) is located at chr11q13.2 and falls within a genomic segment already known to carry prior selection signatures. Voight et al.[1] showed neighboring 100 kb candidate windows for HapMap East Asians and Yoruba (Protocols S1 and S3); out of the 13 genes included in our regional coalescence analysis, seven genes appear in these windows. There have been further scans focusing on the vicinity of *ADRBK1*, for example, `PopHumanScan`'s literature annotations cite the regional GIH and INS South Asian populations from Liu et al.[54]. Our co-ordinate comparison of Johnson & Voight [7] entries with `PopHumanScan` [8] has revealed iHS candidate windows for *GRK2*. The 11q13.2 locus has therefore been documented in prior literature. Our methodological contribution here is (i) the lowest TMRCA falls within *GRK2*, with nearby allele-frequency differences providing additional context, and (ii) TMRCA ranks are consistently low across all five European and all five South Asian populations (0.19–1.16%, with nine of ten below 1%). In GIH, pairwise TMRCA detects this pattern where gene-body iHS is undefined (Fig. 3, Table 2, first row).

**Cross-population replication.** *GRK2* ranks between 0.19% and 1.16% across South Asian and European populations. In South Asia, the ranks are 0.19% in GIH, 0.24% in BEB, 0.27% in PJL, 0.23% in ITU and 0.31% in STU. In Europe, they are 0.33% in IBS, 0.43% in TSI, 0.50% in GBR, 0.69% in CEU and 1.16% in FIN. Nine of the ten populations fall below the 1% cutoff, with FIN just above it.

Across AFR and EAS the ranks range from 7.25–19.39%; PUR and CLM rank 3.45% and 5.51%, respectively. The candidate allele is also present in EAS (CDX 70%, CHB 60%, CHS 56%, JPT 62%, KHV 68%; average 63%), but at lower frequencies than in SAS (95.1%) and EUR (91.9%). TMRCA averages across all haplotype pairs within a population, so the lower frequency of the candidate allele could help explain the weaker EAS signal. Haplotypes with this allele are similar in EAS and SAS. The average number of sequence differences between these groups is 0.96 times that among SAS haplotypes with the allele. Within EAS, haplotypes with and without the allele differ by 8.5 times the average among those with the allele. This similarity is consistent with shared ancestry, while its timing and relationship to selection remain unresolved. Candidate-allele frequencies also vary across AMR populations (PUR 71%, CLM 68%, MXL 61%, PEL 30%). Determining their ancestry and how they reached the Americas would require local ancestry analysis.

The low TMRCA ranks across SAS and EUR indicate relatively recent coalescence at *GRK2*. Previous studies also report regional iHS signals in South Asian and European populations [1, 7, 54]. In GIH, all variants within *GRK2* fall below the minor allele frequency threshold required to calculate iHS. In BEB and IBS, some variants meet this threshold and have iHS scores. TMRCA analysis includes rare variants within the gene, allowing us to examine coalescence where iHS is unavailable.

The TMRCA analysis at *GRK2* uses only modern phased 1000 Genomes data – ancient-DNA results from Akbari *et al.* [16] are used for comparison. Ancient-DNA evidence for selection in the broader 11q13 region comes via rs11604662 at GRCh38 chr11:67,500,577 (Akbari's published GRCh37 coordinate chr11:67,268,048 lifted over to GRCh38 to match our analysis), which lies in the *PITPNM1* intron ∼214 kb downstream of *GRK2* and is included among Akbari's $\pi \geq 0.99$ loci (POSTERIOR = 0.99, $\hat{s} = 0.005$, $P_X = 5.7 \times 10^{-9}$, FDR= 0.01). Importantly, Akbari's genome-wide-significant locus is *PITPNM1*, not *GRK2*: within the queried GRCh37 interval at *GRK2* (chr11:67,033,904–67,054,029), Akbari's maximum $|X|$ is 2.57 and maximum POSTERIOR is 0.114, insufficient for genome-wide significance. Both studies show signals in the same region. Akbari *et al.* track allele-frequency changes at individual variants across 18,000 years of West Eurasian ancient DNA. Our analysis measures pairwise coalescence across gene bodies in present-day populations. These different measures can place the ancient-DNA peak in *PITPNM1* and the lowest TMRCA in *GRK2*. Identifying the selected variant and linking the two signals would require further evidence.

Furthermore, in the 20 kb gene body (chr11:67,266,473–67,286,556) there are only 41 polymorphic variants in GIH out of 400 assayed high-coverage variant positions (approximately 90% of these positions are monomorphic in GIH) none of which pass the `selscan` MAF$\geq$ 5% filter for iHS computation. iHS is undefined if one allele has reached fixation; however, the pairwise coalescence statistic still identifies a *GRK2* gene-body valley in GIH whereas iHS is unavailable (Fig. 3a,g). In BEB, which has 57 polymorphic gene-body variants, there is still more polymorphism in the gene body. The fraction-extreme iHS statistic for the `selscan` analysis places the gene body in the top 0.44% of BEB.

**Variant-level evidence.** We compare allele frequencies in GIH with those in pooled 1000 Genomes populations outside South Asia, using variants within ±500 kb of the gene midpoint. The ALT allele is less common in GIH at 21,445 sites and more common at 3,028 sites, a ratio of 7.1:1. The highest Hudson's $F_{\mathrm{ST}}$ is 0.43. The largest allele-frequency difference occurs at chr11:67,407,126, where the ALT allele has a frequency of 1.9% in GIH and 47% in the pooled comparison group (Table 3). Garud's $H_{12}$ reaches 0.569 in a window centred within *GRK2* at chr11:67,278,452 (Fig. 3i), indicating that a small number of haplotypes account for much of the variation in this window.

**Evidence for a shared haplotype background.** Similar haplotypes across continents could reflect shared ancestry, selection acting independently on existing variants, or gene flow. To compare their sequence backgrounds, we counted SNP differences between pairs of haplotypes in a ±25 kb window containing the most-differentiated SNP at chr11:67,407,126. The dataset included 3,202 individuals, 6,404 phased haplotypes and 1,067 biallelic SNPs, excluding the focal variant. We grouped haplotypes by population and whether they had the candidate allele, whose frequency is 95.1% in SAS, 91.9% in EUR and 16.5% in AFR.

We calculated average differences within and between groups using up to 200 randomly selected haplotypes per group, sampled without replacement. To estimate uncertainty, we repeated the calculation 500 times, each time sampling up to 200 haplotypes with replacement from each full group. The 95% bootstrap intervals span the 2.5th to 97.5th percentiles of the resulting distance ratios. We used the same approach for EAS. These intervals reflect variation from resampling individual haplotypes. Accounting for relationships between haplotypes from the same person or family would require further analysis.

We express the distances relative to the average difference among SAS haplotypes with the candidate allele. On this scale, the distance between SAS and EUR haplotypes with the allele was 1.20 (95% bootstrap interval 1.11–1.35). The distance from SAS haplotypes with the allele to those without it was 8.88 within SAS (7.41–10.03) and 9.73 for the comparison with AFR (8.24–10.63). SAS and EUR haplotypes with the allele are therefore much more similar to each other than to the groups without it, consistent with a shared haplotype background. Distinguishing shared ancestral selection, independent selection on existing variants and gene flow would require further evidence.

**Comparison of TMRCA estimates across three methods.** The TMRCA signatures at *GRK2* in GIH was replicated using `gamma_smc_cu`, `cxt` [55] and `ASMC` [56] (Supplementary Information: Figure S2). All three methods show a qualitative coalescence reduction, although their absolute estimates differ.

**Selection coefficient from ARG inference with `CLUES2`.** We used `Relate` [10] to obtain genealogies at the most-differentiated SNP, located 120 kb downstream of *GRK2* at chr11:67,407,126. From these genealogies, `CLUES2` [57, 58] estimated a selection coefficient of $\hat{s} = 0.00519$ for the derived G allele in GIH (reported 95% CI 0.00218–0.00820, unadjusted $-\log_{10} p = 2.23$). This allele has a frequency of 98.1% in GIH and 53% in pooled 1000 Genomes haplotypes outside South Asia.

`CLUES2` uses variation at this SNP and its genealogies, allowing us to estimate allele-frequency changes even when variation within *GRK2* is limited. The estimates depend on the assumed demographic and selection model. Figure 3d shows the inferred trajectory alongside the *C11orf65* comparison in GIH.

**Biological relevance.** The gene *GRK2* facilitates desensitization of $\beta$-adrenergic receptors; excessive expression of *GRK2* in vascular smooth muscle leads to epinephrine-mediated hypertensive and pulmonary edema in mice [59]. Data from the GTEx (v10 release) indicate a high median *GRK2* transcript expression level in spleen (415 TPM) and blood (372 TPM; https://gtexportal.org/home/gene/GRK2). Cohn *et al.* [60] previously connected the gene to population-level phenotype variation, showing that within a Black-American cohort, lymphocyte *GRK2* protein expression is ∼2-fold higher and GRK kinase activity >40% higher in individuals with elevated systolic blood pressure (≥130 mmHg) than in normotensives. These functional and association studies provide some cardiovascular context for the SAS+EUR candidate. Indeed, Cagliani *et al.* [61] did note that there were three haplotype groups in *ADRB2*, with TMRCA estimates varying between 1.05 to 1.65 million years through various methods, and suggested that the pattern was indicative of either balancing selection or selective sweep in progress. They also reported a possible selective sweep at *ADRB3* in African populations.

#### 3.2 *TREML1 / TREM2*: supporting population and literature comparisons

The *TREML1 / TREM2* locus lies within the TREM immunoreceptor cluster on chromosome 6. *TREML1* is approximately 5.0 kb long and is expressed on platelets and megakaryocytes, where it helps regulate inflammation and clotting at sites of vascular injury [62]. *TREM2* is approximately 4.7 kb long and encodes an innate immune receptor in microglia. Both proteins belong to the TREM receptor family. TREM2 signals through DAP12, while TREML1/TLT-1 has a different intracellular signalling structure.

The rare *TREM2* coding variant rs75932628 (R47H) increases Alzheimer's disease risk and has a minor allele frequency below 1% in Europeans [63, 64]. Here, we examine common variation across the *TREML1*–*TREM2* region.

**Population differences in TMRCA and haplotype structure.** Across the 19 non-African populations, *TREM2* TMRCA ranks range from 1.07% to 5.95%, with ten populations below 2% and 18 below 5%. European ranks fall within a narrower range of 1.07–1.72%, including 1.26% in IBS. African ranks range from 3.05% to 6.50%, overlapping those outside Africa. *TREML1* ranks range from 1.56% to 5.67% across non-African populations. Both genes remain above the reference screen's 1% cutoff.

Garud's $H_{12}$ reaches higher peaks in the seven non-African populations tested (0.58–0.73) than in YRI (0.18), although the peak locations vary. In IBS, FIN, GBR and CEU, the peak is centred within *TREML1* at chr6:41,150,476, reaching 0.646, 0.690, 0.701 and 0.610, respectively. In TSI and JPT, it falls between the two genes at chr6:41,155,516, reaching 0.583 and 0.726. The CHB peak is farther downstream, outside the two-gene block, at chr6:41,214,879 ($H_{12} = 0.653$).

At chr6:41,166,068, approximately 2.9 kb downstream of *TREM2*, the derived G allele has a frequency of 7% in African populations and 52% in non-African populations. Within the two-gene block, the four variants with a minor allele frequency (MAF) of at least 5% in IBS have similar frequencies across European populations. The largest difference between IBS and CEU, FIN, GBR or TSI is 4.7 percentage points.

**Comparison with previous studies.** We examined 52 literature resources for previous findings at this locus. Supplementary Table S6 records the findings and the material available for each comparison. We distinguish a gene missing from a candidate list from a gene that was tested and fell below the study's threshold.

The candidate lists we checked in `PopHumanScan` [8] and five earlier haplotype-based scans [1, 3–5, 25] contain no entry naming *TREML1* or *TREM2*. Searching genomic coordinates, however, identifies several nearby or overlapping findings.

Bitarello *et al.* [9] report candidate regions for balancing selection in LWK, YRI, GBR and TSI. Within chr6:41.0–41.3 Mb (GRCh37), their 80 table entries correspond to 11 distinct 3 kb intervals, some of which overlap. These intervals lie approximately 70–169 kb downstream of *TREM2*.

Sabeti *et al.* [3] list rs9349180 among differentiated alleles in HapMap Phase 2. This variant lies within the *TREML3P* pseudogene, approximately 55 kb downstream of *TREM2*, at GRCh37 chr6:41,185,474.

`Flex-Sweep` [2] lists a 260 kb putative selection region in the YRI population at GRCh37 chr6:40,867,841–41,127,841. There are five upstream genes whose classifier scores are all at least 0.99. *TREML1* has a score of 0.818, greater than the default threshold of 0.5. The reported region covers approximately 1.6 kb of *TREM2*, whereas we only mention *TREML1* of the two focal genes is mentioned in the table. This provides earlier evidence for a regional selection signal in YRI.

Skov *et al.* [40] detect a segment similar to the Altai archaic genome at GRCh38 chr6:40,971,209–41,154,236. The segment contains 51 variants shared with archaic genomes, overlaps most of *TREML1*, and ends about 4.3 kb before *TREM2*. This finding adds evidence of archaic ancestry in the region.

The wider comparison surrounds work with ancient DNA, genealogical studies, and other types of work that explore evolutionary history [6, 10, 11, 13–16, 42–45]. Kerner *et al.* [12] analyzed 2,879 ancient and modern individuals from West Eurasia and found 89 loci under positive selection, including immune genes. TREM is not among their candidates.Deschamps *et al.* [29] examined 1,553 innate-immunity genes and highlighted 57 using Fisher's combined score. Table S6 gives the scope and locus-specific findings for these comparisons.

Maravall-López *et al.* [17] provide linear mixed model (LMM) selection statistics based on the Akbari ancient-DNA framework. Their released results include 13 variants within *TREML1* and four within *TREM2*, with minimum nominal $P$-values of 0.116 and 0.400, respectively. Across the surrounding chr6:41.0–41.3 Mb interval (GRCh37), the smallest $P$-value is 0.0031, above the genome-wide threshold of $5 \times 10^{-8}$. These results use the same underlying ancient-DNA source as the Akbari analysis.

**Gene-level and window-based iHS comparisons.** Our integrated haplotype score (iHS) analysis ranks genes by the fraction of sites with absolute normalised scores above 2. In South Asia, *TREM2* ranks in the top 0.11% in BEB and 0.38% in STU. *TREML1* ranks in the top 0.11% in BEB, 0.21% in ITU, 0.29% in STU and 3.6% in GIH.

In European and East Asian populations, our filtering and scoring procedure produces no iHS scores within *TREM2*. *TREML1* has four scored sites in each European population and one in each East Asian population, all with $|iHS| \leq 2$.

We further looked at iHS statistics provided by the genome browser PopHuman which provides population genetic statistics from the 1000 Genomes Project. The corresponding catalog `PopHumanScan` uses these statistics for the detection of candidate regions under selection [8]. Among the 26 populations considered, the highest mean value of $|iHS|$ within a 10 kb window overlapping either gene was 2.15 in ITU, placing this window in the 99.21st percentile of autosomal values, below the 99.95th-percentile threshold for `PopHumanScan` when considering its 22 non-admixed populations. Windows overlapping either gene in all European and East Asian populations are below the 50th percentile.

PopHuman considers the average absolute value of the iHS statistic within 10 kb windows, while in our analysis we rank the genes on the basis of the fraction of scored sites within each gene with $|iHS| > 2$. This gives the ratios 11/11 and 6/6 for *TREML1* and *TREM2*, respectively, in BEB, 11/12 and 6/7 in STU, and 11/12 and 0/6 in ITU. These high ratios are associated with strong ranks for both genes in BEB and STU and for *TREML1* in ITU. Differences in how scores are combined, the populations and the filters may contribute to the differences between our results and the window-based results.

**Ancient-DNA selection statistics.** We examined the published ancient-DNA selection results from Akbari *et al.* [16]. Among variants passing the study's quality filters, the highest absolute selection statistic within *TREM2* is $|X| = 2.10$, at rs7748513. The highest posterior probability of selection is 0.070 across the *TREML1*–*TREM2* block.

In the surrounding region, the highest posterior probability is 0.75 at rs11760063, approximately 400 kb downstream of *TREM2*. Both values fall below the study's 0.99 threshold for selection in its West Eurasian ancient-DNA analysis.

**Focal-variant selection estimates.** We used `CLUES2` [57, 58] to estimate selection coefficients for seven variants in the region, using 314 haplotypes from 157 IBS individuals. Estimates for the derived alleles range from −0.00185 to 0.00361. One approximate 95% interval is entirely positive, while the other six include zero (Supplementary Table S7).

Figure 3f shows how the estimated allele frequencies changed over time at GRCh38 chr6:41,189,316 and chr6:41,137,356, with 95% credible intervals. It also includes *C11orf65* in GIH for comparison. We chose the seven variants for follow-up after inspecting the regional signal.

**Comparison of TMRCA estimates across three methods.** We compared `gamma_smc_cu`, `ASMC` [56] and `cxt` [55] using the same 20 haplotype pairs. At each of 5,397 shared genomic positions, we took the median TMRCA across these pairs and log-transformed it. Pearson correlations were 0.876 between `gamma_smc_cu` and `ASMC`, 0.930 between `gamma_smc_cu` and `cxt`, and 0.882 between `ASMC` and `cxt` (Supplementary Fig. S2). These comparisons show similar regional patterns across methods using the same genotype data.

We also examined variant density, here the *TREM2* gene body contains 28.4 SNPs per kb, compared with 22.8 SNPs per kb in the flanks. Across the surrounding ±500 kb region, all 100 non-overlapping 10 kb windows contain at least 157 SNPs, with a median of 225 per window. In the Akbari dataset, three of four gene-body variants and 3,319 of 4,055 surrounding variants pass the study's quality filters.

**Location within the TREM cluster.** There are several related genes in this region, including *TREML2*, *TREML4*, *TREM1* and *NCR2* (Supplementary Fig. S4). In IBS data, the highest $H_{12}$ window is centred within *TREML1*, at chr6:41,150,476. The identification of the functional variant would require further fine-mapping and tests of how nearby variants affect gene expression in relevant cell types like microglia, monocytes and megakaryocytes.

A second allele-frequency difference exists at chr6:41,485,209, with a difference of 40 percentage points between IBS and pooled non-European haplotypes. This variant has weak linkage disequilibrium with common variants in the *TREML1*–*TREM2* block (pairwise $r^2 \leq 0.03$ in IBS). The nearest 400-site window plotted has $H_{12} = 0.05$. We treat this as a separate regional feature and report it in Tables 2–3 as part of the surrounding allele-frequency comparison.

*CCDC92*, located on chromosome 12q24.31, encodes a coiled-coil domain protein. Its TMRCA rank is 1.85% in CDX and below 5% in all five East Asian populations. We selected it for follow-up despite its rank being above the 1% cutoff. Compared with populations outside East Asia, the ALT allele is less common in CDX at 6.5 times as many sites as it is more common. The maximum Hudson's $F_{\text{ST}}$ is 0.53.

GWAS associates this genomic region with fat distribution and waist-to-hip ratio [65]. Deletion of *Ccdc92* in mice leads to decreased obesity and insulin resistance associated with a high-fat diet [66]. This gene is located approximately 12 Mb away from the European selection locus at *ALDH2*/*SH2B3* [67, 68].

At rs10773048 (GRCh38 chr12:123,959,862), within *CCDC92*, 172 of 186 CDX haplotypes have the derived C allele. For this allele, `CLUES2` estimates a selection coefficient of $\hat{s} = 0.00438$ (approximate 95% CI $[0.00141, 0.00735]$, unadjusted $-\log_{10} p = 2.04$, Table S7).

Previous studies also report selection signals nearby. Our searches of genomic coordinates in `PopHumanScan` [8] identify candidate regions reported by Liu *et al.* [54] in several East and South Asian populations. Within 500 kb of the gene, Johnson & Voight [7] report iHS candidate windows in CEU, IBS and KHV, and Bitarello *et al.* [9] report balancing-selection windows in GBR and TSI. These windows lie outside the gene body, while our analysis identifies low TMRCA within *CCDC92* in CDX. In West Eurasian ancient DNA, Akbari *et al.* [16] report two nearby variants, rs79478560 and rs58463871, each within 500 kb of the gene and with a posterior probability of selection of 0.99.

#### 3.3 *CLEC6A* (chr12p13.31, CDX) — a candidate directional-selection signal

*CLEC6A* encodes Dectin-2, a receptor that recognises fungal mannan. The gene is part of the Dectin-2 gene cluster on chr12p13.31, adjacent to the Dectin-1 cluster which has expanded through gene duplication in mammals [69]. Its TMRCA ranking is 3.32% for CDX and has a minimum value of 3.30% for KHV. Four out of five East Asian groups have percentages lower than 5% such that the gene falls outside both 1% reference screen and exploratory screen where all five populations have to be lower than 5%. The highest value of $H_{12}$ within ±500 kb is 0.65 while the highest value of Hudson's $F_{\text{ST}}$ is 0.46.

Bitarello *et al.* [9] report balancing-selection windows overlapping *CLEC6A* in LWK, YRI and TSI. Their 13 table entries represent three overlapping 3 kb intervals, repeated across populations, target frequencies and candidate sets. Johnson & Voight [7] also report nearby iHS peaks in ACB, ASW, GWD, PJL and YRI. In the ancient-DNA results from Akbari *et al.* [16], no reported lead variant with a posterior probability of selection of at least 0.99 lies within ±500 kb (Fig. S8b).

At rs1805725 (GRCh38 chr12:8,945,337), approximately 467 kb beyond *CLEC6A*, the derived C allele occurs in 123 of 186 CDX haplotypes. `CLUES2` estimates a selection coefficient of $\hat{s} = 0.00044$, with an

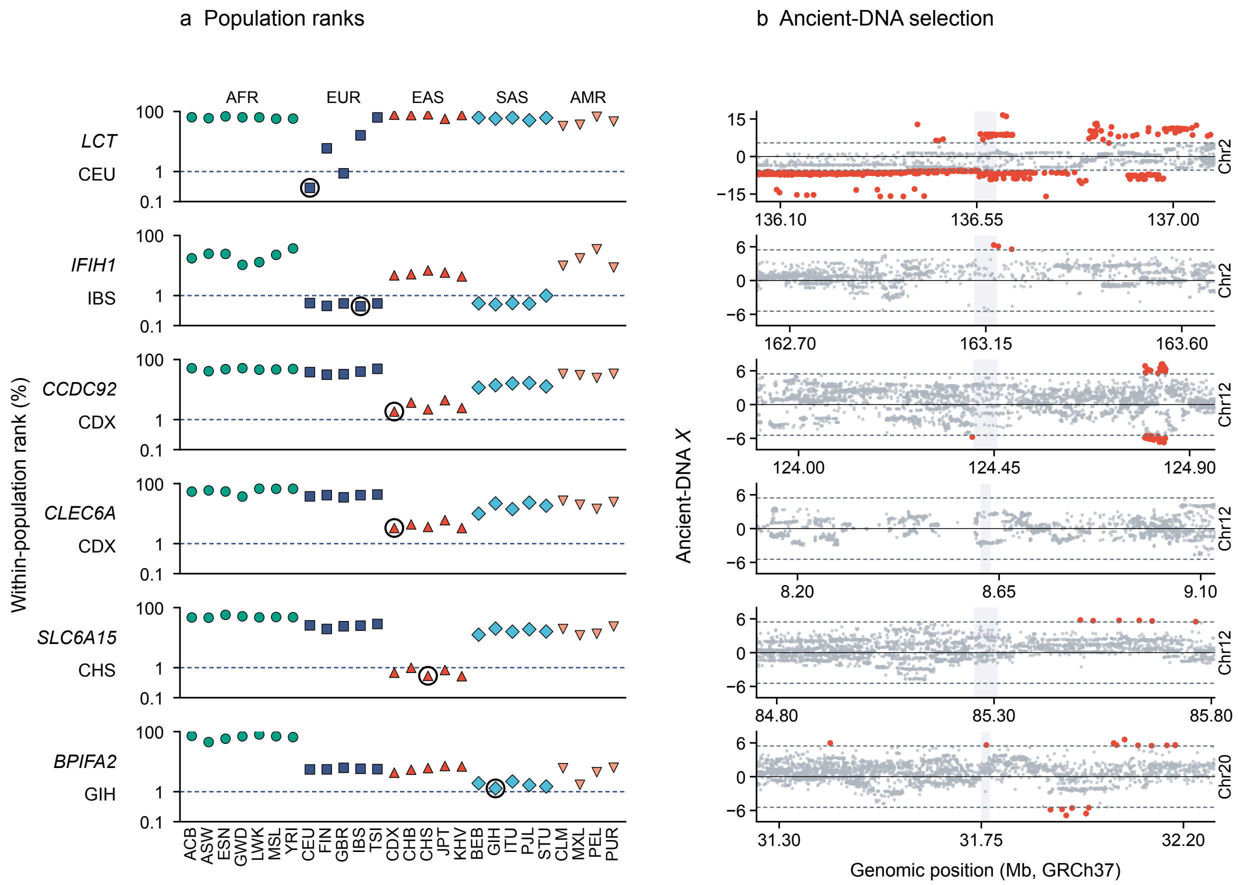

c Variant summaries

| | $\log_{10}$(D/E) | Max. $F_{ST}$ | Max. \|ΔAF\| |
|---|---|---|---|
| *LCT* | 1.10 | 0.64 | 0.68 |
| *IFIH1* | 0.93 | 0.51 | 0.59 |
| *CCDC92* | 0.81 | 0.53 | 0.58 |
| *CLEC6A* | 0.77 | 0.46 | 0.49 |
| *SLC6A15* | 0.72 | 0.60 | 0.61 |
| *BPIFA2* | 0.82 | 0.29 | 0.36 |

d Prior literature

| V06 | S07 | M11 | P09 | G13 | JV18 | PHS | Ak26 |
|---|---|---|---|---|---|---|---|
| ■ | ■ | □ | □ | ■ | ■ | ■ | ■ |
| □ | □ | □ | □ | □ | ■ | ■ | ■ |
| □ | □ | □ | □ | □ | □ | ■ | □ |
| □ | □ | □ | □ | □ | □ | ■ | □ |
| □ | □ | □ | □ | □ | □ | ■ | □ |
| □ | □ | □ | □ | □ | ■ | ■ | ■ |

Figure S8: **Population, variant and literature evidence for five additional case studies and the *LCT* positive control.**(a) TMRCA ranks across 26 populations. Circles highlight focal populations, and the dashed line marks 1%. Marker shapes and colours distinguish continental groups (AFR teal, EUR navy, EAS coral, SAS cyan, AMR peach).(b) Ancient-DNA $X$ statistics [16] within 500 kb of each gene. Pale bands mark gene bodies, and dashed lines mark $|X| = 5.45$. The *LCT* axis is expanded to show its full range.(c) Allele-frequency comparisons showing the $\log_{10}$ ratio of sites where the ALT allele is less common to sites where it is more common in the focal population (D/E), the maximum Hudson's $F_{\mathrm{ST}}$ and the largest absolute allele-frequency difference.(d) Filled squares mark genes named in the candidate lists we checked or directly overlapping a candidate region. Open squares mark genes with no match in those lists. `PopHumanScan` includes several types of selection. For Akbari, a filled square marks a variant within the gene that passes the study's quality filters and has a posterior probability of selection of at least 0.99. Rows follow the same order as in (c).Sources: [1, 3–5, 7, 8, 16, 25].

approximate 95% interval spanning zero ($[-0.00264, 0.00351]$, unadjusted $-\log_{10} p = 0.11$, Table S7).

#### 3.4 *SLC6A15* (chr12q21.3, CHS) — strong East Asian TMRCA signal

*SLC6A15* encodes the sodium-dependent neutral amino-acid transporter $B^0$AT2, which is strongly expressed in the hippocampus. A depression-associated allele has been linked to lower expression of this gene in the hippocampus [70]. Its TMRCA rank is 0.54% in CHS and ranges from 0.52% to 0.99% across the five East Asian populations. Sites where the ALT allele is less common in CHS outnumber those where it is more common by 5.2:1. The maximum Hudson's $F_{ST}$ is 0.60 for CHS compared with populations outside East Asia, and the largest allele-frequency difference occurs at chr12:84,849,501.

Previous studies also report selection signals at or near *SLC6A15*. `PopHumanScan` [8] lists findings in YRI, CHB and CEU, and Barreiro *et al.* [71] identify variants within the gene with $F_{ST} > 0.65$. Johnson & Voight [7] report iHS peaks within 500 kb of the gene in ASW, GWD, KHV, LWK, MXL and TSI. Our analysis adds evidence of low TMRCA within *SLC6A15* in CHS. In the ancient-DNA results from Akbari *et al.* [16], the nearest lead variant with a posterior probability of selection of at least 0.99 is rs74237290. It lies approximately 190 kb beyond the end of the gene at higher genomic coordinates (Fig. S8b).

At GRCh38 chr12:84,886,570, 322 of 326 CHS haplotypes have the derived C allele. For this allele, `CLUES2` estimates a selection coefficient of $\hat{s} = 0.00675$ (approximate 95% CI $[0.00366, 0.00984]$, unadjusted $-\log_{10} p = 3.46$, Table S7).

#### 3.5 *BPIFA2* (chr20q11.21, GIH) — low TMRCA with weak iHS within the gene

*BPIFA2*, also called *SPLUNC2* or *C20orf70*, encodes an antimicrobial protein found in saliva. It ranks among the lowest 1.28% of genes for TMRCA in GIH, also with ranks between 1% and 5% across all five South Asian populations. For every site where the ALT allele is more common in GIH than in the comparison group, about 6.6 sites show the opposite pattern. The highest Hudson's $F_{ST}$ is 0.29. Within the gene, iHS and nSL scores are weak (Table 3), while nearby $H_{12}$ values are elevated (Table S4). The mix of genes in this cluster varies substantially across mammalian species [72].

At rs1570034 (GRCh38 chr20:33,251,230), approximately 70 kb beyond *BPIFA2*, 153 of 206 GIH haplotypes have the derived C allele. For this allele, `CLUES2` estimates a selection coefficient of $\hat{s} = 0.00378$ (approximate 95% CI $[0.00076, 0.00681]$, unadjusted $-\log_{10} p = 1.68$, Table S7).

Previous studies also report selection signals at *BPIFA2*, listed under its alternative name *C20orf70*. `PopHumanScan` [8] records findings in YRI, CHB and CEU and attributes findings in CHS and MEX to Liu *et al.* [54]. Searching the genomic coordinates of Johnson & Voight [7] entries in `PopHumanScan`, we also find iHS candidate windows overlapping the gene in ESN and KHV.

In West Eurasian ancient DNA, Akbari *et al.* [16] report a selection signal at *BPIFA2* that meets their genome-wide significance threshold. The variant rs117186940, within the gene at GRCh37 chr20:31,760,550, passes their quality filters and has $X = 5.57$, a posterior probability of selection of 0.9901 and an estimated selection coefficient of $s = 0.0182$. A second variant, rs293709, lies approximately 169 kb downstream and has a posterior probability of selection of 0.99. Our analysis adds evidence of low TMRCA within *BPIFA2* in the South Asian GIH population.

#### 3.6 *IFIH1* (chr2q24.2, IBS) — gene-level recovery at a previously reported selection locus

*IFIH1* encodes MDA5, a protein that detects double-stranded RNA inside cells and helps activate the type-I interferon immune response. Fumagalli *et al.* [73] found evidence of local selection at the gene, while the type-1-diabetes susceptibility alleles showed patterns consistent with neutral or nearly neutral evolution. `PopHumanScan` [8] includes *IFIH1* among the candidate annotation features. Additionally, our coordinate comparison identifies nearby iHS candidate windows reported by Johnson & Voight [7].

In our analysis, *IFIH1* has a TMRCA rank of 0.44% in IBS. Sites where the ALT allele is less common in the focal population outnumber those where it is more common by 8.5:1, and the maximum Hudson's $F_{ST}$ is 0.51 (Table 3). We include it as a previously reported locus where our analysis also identifies low coalescence times within the gene. This finding adds to the evidence considered in studies of selection on interferon signalling [26]. Akbari *et al.* [16] also report selection signals in the broader 2q24 region (Fig. S8b).

#### 3.7 Interpretation of reference-locus ranks

Four reference loci remain above the 10% rank threshold. *HBB* (82.7% in PEL) illustrates that the shallow-coalescence criterion need not recover a balancing-selection locus [74]. *MC1R* (25.7% in JPT) has been reported to show strong functional constraint in Africa and relaxed constraint outside Africa [75]. *OCA2* (24.8% in JPT) shows limited gene-level TMRCA depression. *EPAS1* (50.9% in PEL) is a Tibetan high-altitude adaptation locus [76], and this cohort lacks a Tibetan population. *TYRP1* ranks at 8.49% in KHV (Fig. S1).

A few observations are worth making here. First, the *ALDH2* locus (whose alcohol-metabolism variant rs671 is concentrated in East Asian populations [77]) shows its lowest TMRCA rank in TSI (1.24%), with ranks of 3.68% in FIN, 3.88% in IBS, 4.53% in GBR, 6.93% in CEU. Notably, this low-TMRCA signal may not correspond to an independently selected target at the *ALDH2* locus itself, but instead reflect hitchhiking on the well-known sweep at chromosome 12q24 in Europeans, where *SH2B3* R262W (rs3184504 – celiac disease and autoimmunity) has been proposed as the functional driver [67]. We note however that an alternative interpretation has been put forward by Schaschl *et al.* [68], who argue that selection in Europeans acts directly on regulatory variants affecting *ALDH2* expression rather than on *SH2B3*. These genes lie in the same chromosome-12q24 region, with a ∼315 kb gap from the end of the *SH2B3* gene body to the start of *ALDH2* in our GENCODE annotation. Linked selection could therefore produce a TMRCA signal at *ALDH2* without identifying it as the causal target. Consistent with this, the TMRCA signal at *SH2B3* itself reaches 1.95% in TSI (see Table 1). A Han Chinese selection scan also identifies the *ALDH2* region [23]. Second, the low-TMRCA region around the *CYP3A* cluster on chromosome 7 overlaps a region with reported non-African selection signatures and a proposed role for salt-homeostasis adaptation [78]. Third, *FADS1* has a lower TMRCA rank in ITU (0.93%) than the neighbouring *FADS2* (7.20%). The difference identifies a stronger gene-body coalescent signal at *FADS1*. Finally, our within-population ranking places several classical HLA genes, including *HLA-A*, *HLA-B* and *HLA-C*, above the 99th percentile across all 26 populations, consistent with the established balancing-selection signature at HLA [79, 80].

#### 3.8 Prior-literature status across the five SI case studies

Across these five loci (Table 2, Table S2): *CCDC92*, *SLC6A15* and *BPIFA2* have prior regional selection reports, alongside the gene-level TMRCA signals examined here. *CLEC6A* overlaps published balancing-selection windows and shows a shallow TMRCA in CDX. *IFIH1* provides a literature-supported comparison for gene-level recovery. The published candidate gene annotations inspected in `PopHumanScan` and the scans of Voight, Sabeti, Metspalu, Pickrell and Grossman contain no entry naming *TREML1* or *TREM2*, although unnamed overlapping windows or subthreshold signals are not excluded. Gene-level iHS recovers the locus in BEB/STU/ITU at top 0.4%, but differences in aggregation, cohorts and filtering prevent attributing the catalog miss solely to window dilution. *TREM2* ranks remain in the lower tail (1.07–5.95% in non-African panels) but fail the 1% criterion in all populations. A full per-resource comparison is in Table S6.

##### 3.9 Focal-variant selection estimates

Table S7: **Exploratory `CLUES2` estimates at all analysed focal variants.** Coordinates are GRCh38. Counts give observed derived haplotypes over the population sample – the conditioning frequency uses that observed count. The Flips column counts leaf-state changes made by the converter to reconcile observed alleles with the focal genealogy. The GRK2 and C11orf65 arguments were rounded to 0.981 and 0.311, respectively, selecting the same frequency-grid states as their exact fractions. Confidence intervals are the `CLUES2` approximation. Likelihood-ratio statistics are nominal and unadjusted for the post hoc selection of loci and variants. The C11orf65 comparison is in GIH and is not a demonstrated neutral locus or an IBS-matched control.

| Locus | Pop. | Chr:position | Derived/count | Flips | $\hat{s}$ | Approx. 95% CI | $-\log_{10} p$ |
|---|---|---|---|---|---|---|---|
| *GRK2* | GIH | 11:67,407,126 | G 202/206 | 0 | 0.00519 | $[0.00218, 0.00820]$ | 2.23 |
| *C11orf65* | GIH | 11:108,387,060 | T 64/206 | 0 | −0.00068 | $[-0.00350, 0.00215]$ | 0.20 |
| *TREM2* | IBS | 6:41,121,942 | A 208/314 | 0 | 0.00054 | $[-0.00286, 0.00394]$ | 0.12 |
| *TREM2* | IBS | 6:41,137,356 | T 175/314 | 2 | 0.00361 | $[0.00050, 0.00672]$ | 1.58 |
| *TREM2* | IBS | 6:41,166,068 | G 179/314 | 0 | 0.00240 | $[-0.00032, 0.00512]$ | 1.07 |
| *TREM2* | IBS | 6:41,176,920 | C 212/314 | 0 | 0.00198 | $[-0.00106, 0.00501]$ | 0.65 |
| *TREM2* | IBS | 6:41,189,316 | T 203/314 | 15 | 0.00048 | $[-0.00272, 0.00369]$ | 0.12 |
| *TREM2* | IBS | 6:41,189,932 | C 111/314 | 0 | −0.00185 | $[-0.00463, 0.00093]$ | 0.68 |
| *TREM2* | IBS | 6:41,191,484 | G 203/314 | 0 | 0.00193 | $[-0.00050, 0.00436]$ | 0.91 |
| *BPIFA2* | GIH | 20:33,251,230 | C 153/206 | 4 | 0.00378 | $[0.00076, 0.00681]$ | 1.68 |
| *SLC6A15* | CHS | 12:84,886,570 | C 322/326 | 0 | 0.00675 | $[0.00366, 0.00984]$ | 3.46 |
| *CCDC92* | CDX | 12:123,959,862 | C 172/186 | 0 | 0.00438 | $[0.00141, 0.00735]$ | 2.04 |
| *CLEC6A* | CDX | 12:8,945,337 | C 123/186 | 0 | 0.00044 | $[-0.00264, 0.00351]$ | 0.11 |

### 4 Extended Methods

#### 4.1 Genome-wide covariate and haplotype comparisons

**Analysis set and genomic annotations.** Supplementary Fig. S6 preserves the corrected 19,119-gene gene list, its original within-population ranks and its unique-base segmental-duplication mask. We obtained site counts from the 572 chromosome–population datasets used to build the gene list. Counts for all 497,094 gene–population combinations matched those in the corrected SNP data used for XP-EHH. Covariates were gene-body length, site count, deCODE sex-averaged GRCh38 recombination rate from the `stdpopsim` catalog [81], and the Buffalo–Kern YRI CADD6/deCODE $B'$ annotation [82]. Map gaps were preserved, with at least 95% coverage required in both maps. Excluding SD-masked genes and undefined gene list ranks left 17,803–17,936 genes per population (the complete-covariate sets contained 14,193–14,298). These are shared reference annotations, not population-specific calibrations for all 26 populations. We repeated the comparisons using `pyrho`-YRI recombination, Buffalo–Kern CEU $B'$ and regulatory/phastCons background-selection (BGS) annotations to check sensitivity to the choice of map and model. Each comparison used the genes with available values for those annotations.

**Rank dependence and conditional focal ranks.** Marginal Spearman correlations and partial correlations of ranked variables used the same complete-covariate set within each population – each partial correlation controlled the other three ranked covariates. Table S8 reports the estimates. Leave-one-chromosome-out jackknife intervals use the 22 autosomes and $\hat{\theta} \pm 1.96\,\mathrm{SE}_{\mathrm{jackknife}}$. Within-population covariate quintiles contain approximately 20% of eligible genes each. The figure shows the median across populations of each group's median TMRCA rank, with the range across populations as a descriptive band. For the twelve focal examples, covariates were converted to within-population percentiles. Controls were on other chromosomes and within 0.20 in each percentile coordinate – the nearest at most 500 by Euclidean distance were used, with a minimum of 100 required. Conditional rank was $(1+n_{<}+n_{=}/2)/(1+n_{\mathrm{controls}})$. Missing map coverage or insufficient matching remained undefined. These conditional ranks use different denominators and do not replace the original candidate-screen ranks. Conditioning on site count can remove biological signal as well as assay dependence.

**Aligned iHS and genome-wide XP-EHH.** `selscan` iHS scores were normalized using the chromosome/population frequency-bin procedure in the main Methods. Normalization used the full historical stream before removing every duplicated physical position and including only scored cores present in the corrected phased biallelic SNP cache. Historical haplotype construction, including its low-frequency filtering, was unchanged: this alignment is not a fresh iHS inference from corrected haplotypes. Unlike the historical gene summaries, all summaries here record missing values explicitly and rank only finite non-SD gene scores. Consequently their percentiles need not equal the historical case-study percentiles.

XP-EHH was newly computed with `scikit-allel` [83] over full chromosomes, separately for each population against YRI; YRI was compared against CEU. Sites segregating in the pooled comparison were used, including fixed differences and sites fixed in either single population, without a focal MAF cutoff. Physical-distance integration used minimum EHH 0.05, gap scaling 20 kb, maximum gap 200 kb and exclusion of incomplete chromosome-edge scores. Positive values indicate longer focal-population haplotypes. All finite scores across the 22 autosomes were standardized together for each contrast. This analysis differs from the historical pooled-superpopulation regional XP-EHH summaries in Table S4.

Regional summaries used the disjoint 100 kb on each side of the gene, excluding the gene body. 500-kb flanks and maximum-score summaries were used as sensitivities. The primary summary was the fraction of finite cores with $|\text{iHS}| > 2$ or normalized XP-EHH $> 2$. We recorded the numbers of returned, finite and extreme scores separately. When no finite scores were available, we recorded the fraction as missing. Haplotype percentiles were descending average ranks among computable non-SD genes in the population. Strictly below 5% defines top-tail support; ties mean this need not include exactly 5% of genes. Associations with decreasing TMRCA rank used marginal and partial Spearman correlations on each statistic's computable complete-covariate set, controlling all four ranked covariates for the partial estimates. Score-availability summaries were also computed for all non-SD finite-rank genes. Their population-averaged intervals used 1,000 whole-chromosome bootstrap samples (seed 20260927), with chromosome weights shared across populations and statistics. Missing body iHS was separated into absence of a corrected-cache core at MAF $\geq$ 5% and absence of a finite aligned score despite such a core.

**Matched haplotype support.** Cases had original TMRCA rank below 1% and controls had rank at least 1%. Both required complete covariates, and controls exactly matched body-iHS availability. Candidate controls lay on other chromosomes within 0.15 in every covariate-percentile coordinate. Up to 20 nearest controls were used, with at least five required. Of 2,414 eligible cases, all 2,008 with body iHS matched; 277 of 406 without body iHS matched and 129 did not. For each statistic, a case needed a finite score and at least five finite-score controls among its selected matches. The difference in top-tail frequency is the mean across computable cases of case top-tail status minus the proportion among that case's finite controls. Each case receives equal weight. All 43,100 case–control pairs met the matching rules: the genes were on different chromosomes and differed by at most 0.15 in each covariate percentile, on a scale from 0 to 1. Within each population, we averaged each covariate difference across the matched pairs. The largest absolute mean difference was 0.028.

Confidence intervals use a 22-autosome leave-one-out jackknife. We left out one chromosome at a time, removed its genes from both the case and control groups, and repeated the matching. Covariate percentiles and score thresholds remained fixed at their full-genome values. An interval crossing zero does not establish excess support. These intervals account for chromosome blocks, not all demographic dependence or map-estimation uncertainty. Neither matching nor concordance of statistics establishes a causal selective history or calibrated selection FDR.

Table S8: **Genome-wide covariate dependence of TMRCA rank.** Marginal and partial Spearman correlations with increasing TMRCA rank. Within each population, all estimates use the same $n$ genes with complete covariates and no overlap with segmental duplications (SD). Partial correlations control the other three ranked covariates. Sites denotes site count, recombination uses the deCODE reference map, and background selection (BGS) uses YRI $B'$, the estimated fraction of neutral diversity remaining under BGS.

| Population | $n$ | Marginal correlation | | | | Partial correlation | | | |
|---|---|---|---|---|---|---|---|---|---|
| | | Length | Sites | Recomb. | BGS | Length | Sites | Recomb. | BGS |
| ACB | 14,298 | 0.066 | 0.200 | 0.404 | 0.570 | -0.625 | 0.633 | 0.146 | 0.261 |
| ASW | 14,280 | 0.066 | 0.210 | 0.402 | 0.567 | -0.636 | 0.645 | 0.139 | 0.264 |
| LWK | 14,291 | 0.067 | 0.212 | 0.410 | 0.573 | -0.646 | 0.655 | 0.150 | 0.251 |
| GWD | 14,296 | 0.065 | 0.199 | 0.405 | 0.571 | -0.623 | 0.631 | 0.142 | 0.262 |
| MSL | 14,287 | 0.069 | 0.212 | 0.411 | 0.576 | -0.642 | 0.651 | 0.150 | 0.255 |
| ESN | 14,292 | 0.069 | 0.208 | 0.402 | 0.566 | -0.627 | 0.636 | 0.143 | 0.256 |
| YRI | 14,291 | 0.069 | 0.203 | 0.405 | 0.570 | -0.618 | 0.627 | 0.145 | 0.260 |
| CEU | 14,255 | 0.038 | 0.172 | 0.326 | 0.476 | -0.524 | 0.533 | 0.091 | 0.237 |
| TSI | 14,238 | 0.043 | 0.187 | 0.326 | 0.485 | -0.528 | 0.539 | 0.086 | 0.254 |
| FIN | 14,214 | 0.044 | 0.200 | 0.322 | 0.470 | -0.540 | 0.552 | 0.091 | 0.221 |
| GBR | 14,222 | 0.043 | 0.195 | 0.327 | 0.475 | -0.539 | 0.551 | 0.093 | 0.231 |
| IBS | 14,268 | 0.039 | 0.168 | 0.326 | 0.483 | -0.504 | 0.512 | 0.090 | 0.245 |
| CHB | 14,223 | 0.027 | 0.177 | 0.328 | 0.481 | -0.508 | 0.518 | 0.100 | 0.257 |
| JPT | 14,203 | 0.042 | 0.198 | 0.335 | 0.482 | -0.511 | 0.524 | 0.107 | 0.249 |
| CHS | 14,253 | 0.039 | 0.171 | 0.335 | 0.484 | -0.490 | 0.500 | 0.105 | 0.256 |
| CDX | 14,193 | 0.036 | 0.188 | 0.336 | 0.487 | -0.509 | 0.520 | 0.101 | 0.255 |
| KHV | 14,235 | 0.029 | 0.170 | 0.326 | 0.478 | -0.502 | 0.510 | 0.096 | 0.251 |
| GIH | 14,239 | 0.040 | 0.187 | 0.337 | 0.504 | -0.555 | 0.565 | 0.085 | 0.258 |
| PJL | 14,271 | 0.041 | 0.175 | 0.340 | 0.505 | -0.544 | 0.552 | 0.090 | 0.261 |
| BEB | 14,263 | 0.042 | 0.176 | 0.342 | 0.510 | -0.541 | 0.549 | 0.089 | 0.267 |
| STU | 14,262 | 0.043 | 0.185 | 0.341 | 0.506 | -0.548 | 0.557 | 0.093 | 0.256 |
| ITU | 14,261 | 0.043 | 0.187 | 0.340 | 0.507 | -0.548 | 0.558 | 0.092 | 0.259 |
| MXL | 14,252 | 0.041 | 0.182 | 0.328 | 0.491 | -0.506 | 0.517 | 0.083 | 0.253 |
| PUR | 14,285 | 0.041 | 0.162 | 0.338 | 0.505 | -0.493 | 0.500 | 0.086 | 0.248 |
| CLM | 14,282 | 0.038 | 0.163 | 0.335 | 0.497 | -0.487 | 0.495 | 0.091 | 0.246 |
| PEL | 14,242 | 0.053 | 0.176 | 0.306 | 0.466 | -0.417 | 0.430 | 0.066 | 0.256 |

#### 4.2 Simulations to evaluate sweep detection and localisation

**Simulation design.** We tested how well each method detects selective sweeps and locates the selected variant. We fixed the simulation design and random seeds before evaluating the results, separately from the simulations fitted to the empirical loci.

We simulated two populations, each with diploid $N_e$ = 10,000, that split 6,000 generations ago from an ancestral population of the same size. Population sizes remained constant, with no migration. Each simulated sequence was 1 Mb long, with uniform recombination and mutation rates of $10^{-8}$ and $1.25 \times 10^{-8}$ per base per generation, respectively. We used `stdpopsim` 0.3.0 [84] with `SLiM` 4.3 [85] and no rescaling ($Q = 1$). Forward simulations began with a burn-in of $0.1N_e$ generations before the population split. We then used `msprime` [86] to complete the older neutral ancestry of lineages that had not yet coalesced. We sampled 50 diploid individuals from each population.

We introduced one beneficial mutation into the focal population at a position chosen uniformly within the central 200 kb. Fitnesses were 1, $1 + s/2$ and $1 + s$ for individuals with zero, one or two copies of the beneficial allele, respectively, with $s = 0.02$. We simulated four sweep histories:

- **Incomplete sweep:** the mutation was introduced 800 generations ago and reached a population frequency of 0.25–0.75 at sampling.
- **Near fixation:** the mutation was introduced 1,200 generations ago and reached a population frequency of 0.80–0.99 at sampling.

- **Completed sweep:** the mutation was introduced 2,000 generations ago and reached fixation by sampling.
- **Older completed sweep:** the mutation was introduced 4,000 generations ago and reached fixation at least 2,000 generations before sampling.

The simulations were run until they satisfied the frequency and fixation criteria corresponding to the specific history that was designated for them. The criteria dictated which simulations would be included in the benchmark. The inference procedures relied on the genotypes generated by those simulations. The accepted allele-frequency dynamics, number of simulation restarts, the frequencies at the end of the simulation and time to fixation were recorded. Neutral simulations used the same demographic model, with a randomly chosen position serving as the target for detection and localisation.

We used 1,000 independent simulation replicates. Of these, 600 were neutral simulations, divided into three separate sets of 200 to standardise scores, set detection thresholds and evaluate false-positive rates. The remaining 400 were used to evaluate sweep detection, with 100 replicates for each of the four sweep histories. Each replicate had a reproducible random seed based on its dataset, sweep or neutral history, and replicate number. Before running the inference methods, we removed the selected variant so that detection relied on nearby neutral markers linked to it.

**TMRCA and haplotype scores.** We used the CPU reference implementation, `gamma_smc`, to analyse all variable sites and all 4,950 pairs among the 100 focal-population haplotypes across the full 1 Mb sequence. We fixed $\theta = 4N_e\mu = 0.0005$ and $\rho = 4N_e r = 0.0004$ across replicates and converted posterior mean TMRCA to generations using $2N_e = 20{,}000$. Each interval required at least two sites. Its score was the negative average of log posterior mean TMRCA across pairs and sites, so higher scores indicate more recent coalescence.

We estimated iHS and XP-EHH using `scikit-allel` 1.3.13 [83]. Sites for iHS required a minor allele frequency of 5% or higher in the focal population. Sites for XP-EHH varied in the two populations in combination, including fixed differences between them. The positive XP-EHH values correspond to longer haplotypes in the focal population. EHH minimum value, gap scaling, and maximum gap size were 0.05, 20 kb, and 200 kb, respectively. Scores with incomplete haplotype windows at the edge of sequences were excluded.

The standardisation of both statistics relied on 200 neutral standardisation replicates. For iHS we used 20 derived allele frequency bins with at least 50 finite scores. For XP-EHH the standardisation relied on the overall neutral mean and standard deviation. The same standardisation approach was applied to the evaluation replicates.

For each interval, the primary statistic was the number of scores above the threshold ($|\mathrm{iHS}_z| > 2$ or $\mathrm{XP\text{-}EHH}_z > 2$) out of all available standardised scores. Missing scores were not counted. We used also alternative comparison measures based on the maximum absolute iHS and maximum signed XP-EHH in each interval and on the proportion of high scores in separate 100 kb intervals to the left and right of the target interval.

**True and inferred TMRCA profiles.** For Fig. S7a–d, we used the first evaluation replicate of each sweep history in the recorded identifier order. True TMRCA profiles show the geometric mean across all 4,950 focal haplotype pairs for each local genealogical tree. Inferred profiles show the geometric mean across the same pairs at each focal-population SNP. Haplotype scores are plotted at the sites where they were calculated, using absolute standardised iHS and signed standardised XP-EHH. These profiles are shown without smoothing or window averaging. We checked the true TMRCA calculation for 20 local trees by calculating TMRCA separately for every pair.

We also aligned the profiles by their target position across all 600 evaluation replicates, including 200 neutral replicates and 100 for each sweep history. At each SNP in the focal population, we calculated the true mean log TMRCA across all 4,950 haplotype pairs. Pairs sharing the same most recent common ancestor have the same TMRCA, so we grouped them by ancestor and weighted each ancestor's log age by the number of pairs in that group. We checked this calculation in 150 local trees by calculating TMRCA separately for every pair.

We averaged true and inferred log TMRCA over the same sites in 20 kb windows, each containing at least two sites. We then exponentiated these averages to obtain geometric means. Window centres moved in 5 kb steps from 250 kb before to 250 kb after the target. Haplotype profiles used the maximum-score measures defined above. At each position, lines show the median across replicates with available scores, and shaded bands show the interquartile range. We also report how many replicates contributed at each position.

**Detection thresholds and missing scores.** We measured detection in 5 kb and 20 kb intervals centred on the selected variant or the randomly chosen neutral target. Intervals included their left boundary and excluded their right boundary. For each method and interval width, we set a separate threshold using the 200 neutral calibration replicates. We sorted their scores from lowest to highest, placing missing scores at $-\infty$, and used the 191st score as the threshold. A detection required a finite score strictly above this threshold.

This rule targets a false-positive rate of 5%, with ties potentially lowering the rate. We measured the resulting false-positive rate using the 200 independent neutral evaluation replicates. For each sweep history, detection power was the fraction of all 100 evaluation replicates exceeding the threshold, with missing scores counted as non-detections. We also report the fraction of replicates with a computable score and detection power among those replicates. The maximum-score comparisons in Table S10 show how results change when interval scores use the strongest site rather than the fraction of high-scoring sites.

**Locating the selected variant.** We scanned the central 500 kb with windows of each width, moving 5 kb at a time. For detection anywhere in this scan, we calibrated separate thresholds using the maximum score from each neutral calibration replicate.

For each selected replicate, we identified the highest-scoring window. When several windows tied, we calculated the distance from the true selected position to each tied window centre and used the median distance as the localisation error. We report the median and interquartile range of these errors across all selected replicates with computable scores (Table S11). We also report medians restricted to detected replicates, how often windows tied for the maximum, and the genomic span of those ties.

Detection rates, neutral call rates and score availability are reported with Wilson 95% intervals across independent replicates. For differences between methods, we used 2,000 bootstrap resamples of replicate identifiers, keeping each replicate's results paired across methods.

Table S9: **Held-out neutral call rates in the controlled benchmark.** Percentages and Wilson 95% confidence intervals among 200 independent neutral evaluation replicates. For each window width, statistic and detection approach, we set a threshold using 200 separate neutral simulations to target a 5% false-positive rate. A call requires a score above the threshold. Tied scores can therefore give rates below 5%. Missing scores count as non-calls. Target intervals are centred on randomly chosen positions. For scans, we use the highest-scoring window in the central 500 kb. This benchmark uses the CPU implementation, `gamma_smc`.

| Statistic | Width (kb) | Target calls (%) | Scan calls (%) |
|---|---|---|---|
| `gamma_smc` | 5 | 3.5 (1.7–7.0) | 4.0 (2.0–7.7) |
| iHS fraction | 5 | 4.0 (2.0–7.7) | 0.0 (0.0–1.9) |
| XP-EHH fraction | 5 | 0.0 (0.0–1.9) | 0.0 (0.0–1.9) |
| iHS, 100-kb flanks | 5 | 2.0 (0.8–5.0) | 3.5 (1.7–7.0) |
| XP-EHH, 100-kb flanks | 5 | 2.5 (1.1–5.7) | 3.0 (1.4–6.4) |
| Maximum absolute iHS | 5 | 7.5 (4.6–12.0) | 4.5 (2.4–8.3) |
| Maximum XP-EHH | 5 | 0.0 (0.0–1.9) | 2.0 (0.8–5.0) |
| `gamma_smc` | 20 | 4.5 (2.4–8.3) | 5.5 (3.1–9.6) |
| iHS fraction | 20 | 5.5 (3.1–9.6) | 4.5 (2.4–8.3) |
| XP-EHH fraction | 20 | 3.5 (1.7–7.0) | 4.0 (2.0–7.7) |
| iHS, 100-kb flanks | 20 | 1.5 (0.5–4.3) | 3.5 (1.7–7.0) |
| XP-EHH, 100-kb flanks | 20 | 3.5 (1.7–7.0) | 3.0 (1.4–6.4) |
| Maximum absolute iHS | 20 | 6.0 (3.5–10.2) | 4.5 (2.4–8.3) |
| Maximum XP-EHH | 20 | 2.0 (0.8–5.0) | 2.0 (0.8–5.0) |

Table S10: **Detection with alternative haplotype summaries.** Percentages and Wilson 95% intervals across 100 independent replicates per selected history. Maximum scores use absolute iHS or signed XP-EHH inside the target window. Flank scores use the fraction of scored sites with |iHS| > 2 or XP-EHH > 2 in the separate 100-kb intervals immediately to the left and right of the target window. All thresholds use independent neutral calibration. Missing scores count as non-detections. Corresponding neutral call rates are in Table S9.

**Target-interval detection**

| Width (kb) | History | iHS max. (%) | XP-EHH max. (%) | iHS flanks (%) | XP-EHH flanks (%) |
|---|---|---|---|---|---|
| 5 | Incomplete | 81.0 (72.2–87.5) | 90.0 (82.6–94.5) | 95.0 (88.8–97.8) | 95.0 (88.8–97.8) |
| 5 | Near fixation | 8.0 (4.1–15.0) | 100.0 (96.3–100.0) | 91.0 (83.8–95.2) | 100.0 (96.3–100.0) |
| 5 | Completed | 2.0 (0.6–7.0) | 99.0 (94.6–99.8) | 70.0 (60.4–78.1) | 99.0 (94.6–99.8) |
| 5 | Older completed | 2.0 (0.6–7.0) | 70.0 (60.4–78.1) | 8.0 (4.1–15.0) | 89.0 (81.4–93.7) |
| 20 | Incomplete | 93.0 (86.3–96.6) | 93.0 (86.3–96.6) | 95.0 (88.8–97.8) | 95.0 (88.8–97.8) |
| 20 | Near fixation | 22.0 (15.0–31.1) | 100.0 (96.3–100.0) | 92.0 (85.0–95.9) | 100.0 (96.3–100.0) |
| 20 | Completed | 7.0 (3.4–13.7) | 99.0 (94.6–99.8) | 70.0 (60.4–78.1) | 99.0 (94.6–99.8) |
| 20 | Older completed | 1.0 (0.2–5.4) | 74.0 (64.6–81.6) | 4.0 (1.6–9.8) | 87.0 (79.0–92.2) |

**Scan-level detection**

| Width (kb) | History | iHS max. (%) | XP-EHH max. (%) | iHS flanks (%) | XP-EHH flanks (%) |
|---|---|---|---|---|---|
| 5 | Incomplete | 95.0 (88.8–97.8) | 93.0 (86.3–96.6) | 95.0 (88.8–97.8) | 95.0 (88.8–97.8) |
| 5 | Near fixation | 87.0 (79.0–92.2) | 100.0 (96.3–100.0) | 97.0 (91.5–99.0) | 100.0 (96.3–100.0) |
| 5 | Completed | 51.0 (41.3–60.6) | 99.0 (94.6–99.8) | 86.0 (77.9–91.5) | 99.0 (94.6–99.8) |
| 5 | Older completed | 4.0 (1.6–9.8) | 73.0 (63.6–80.7) | 7.0 (3.4–13.7) | 86.0 (77.9–91.5) |
| 20 | Incomplete | 95.0 (88.8–97.8) | 93.0 (86.3–96.6) | 95.0 (88.8–97.8) | 95.0 (88.8–97.8) |
| 20 | Near fixation | 87.0 (79.0–92.2) | 100.0 (96.3–100.0) | 96.0 (90.2–98.4) | 100.0 (96.3–100.0) |
| 20 | Completed | 51.0 (41.3–60.6) | 99.0 (94.6–99.8) | 85.0 (76.7–90.7) | 99.0 (94.6–99.8) |
| 20 | Older completed | 4.0 (1.6–9.8) | 73.0 (63.6–80.7) | 6.0 (2.8–12.5) | 86.0 (77.9–91.5) |

Table S11: **Localisation under primary and alternative summaries.** Median error (interquartile range) in kb across all computable selected replicates; the subscript gives the number of computable replicates out of 100. Each replicate includes all tied strongest windows and uses their median absolute distance from the true target. Windows scan the central 500 kb in 5-kb steps. These summaries are not conditioned on detection. Flank scores use disjoint 100-kb regions on each side of a window, while the other scores use the window itself.

**5-kb windows**

| Statistic | Incomplete | Near fixation | Completed | Older completed |
|---|---|---|---|---|
| `gamma_smc` | 42.7 (21.2–78.3)$_{100}$ | 8.4 (4.0–17.3)$_{100}$ | 8.7 (4.2–18.5)$_{100}$ | 22.4 (9.8–46.0)$_{100}$ |
| iHS fraction | 87.2 (70.4–101.3)$_{100}$ | 104.1 (82.4–128.0)$_{100}$ | 108.8 (80.4–139.3)$_{100}$ | 92.6 (42.3–151.4)$_{100}$ |
| XP-EHH fraction | 77.3 (48.1–95.0)$_{100}$ | 75.1 (63.2–90.5)$_{100}$ | 60.5 (46.7–76.3)$_{100}$ | 45.2 (25.1–70.6)$_{100}$ |
| iHS, 100-kb flanks | 64.2 (34.9–107.7)$_{100}$ | 66.0 (33.7–107.8)$_{100}$ | 72.0 (39.9–124.5)$_{100}$ | 95.1 (45.5–211.6)$_{100}$ |
| XP-EHH, 100-kb flanks | 47.9 (31.8–79.4)$_{100}$ | 34.3 (17.1–53.5)$_{100}$ | 31.8 (14.0–51.1)$_{100}$ | 50.1 (21.0–72.4)$_{100}$ |
| Maximum absolute iHS | 50.4 (17.2–99.6)$_{100}$ | 97.2 (54.5–143.5)$_{100}$ | 142.1 (80.8–185.8)$_{100}$ | 142.1 (75.5–232.5)$_{100}$ |
| Maximum XP-EHH | 37.2 (15.0–83.5)$_{100}$ | 20.0 (7.0–47.8)$_{100}$ | 29.7 (14.0–62.5)$_{100}$ | 38.4 (17.5–85.0)$_{100}$ |

**20-kb windows**

| Statistic | Incomplete | Near fixation | Completed | Older completed |
|---|---|---|---|---|
| `gamma_smc` | 34.3 (16.4–65.9)$_{100}$ | 7.4 (3.8–15.7)$_{100}$ | 9.4 (3.8–20.6)$_{100}$ | 19.7 (7.9–48.5)$_{100}$ |
| iHS fraction | 67.9 (46.4–94.3)$_{100}$ | 80.7 (56.2–105.9)$_{100}$ | 70.8 (48.4–109.4)$_{100}$ | 82.7 (37.2–154.0)$_{100}$ |
| XP-EHH fraction | 70.0 (42.5–90.0)$_{100}$ | 67.4 (54.5–81.8)$_{100}$ | 50.4 (33.7–65.2)$_{100}$ | 35.0 (19.9–68.0)$_{100}$ |
| iHS, 100-kb flanks | 64.9 (36.0–103.2)$_{100}$ | 65.2 (35.9–105.8)$_{100}$ | 65.7 (32.5–102.4)$_{100}$ | 96.5 (43.1–202.9)$_{100}$ |
| XP-EHH, 100-kb flanks | 51.1 (33.8–79.4)$_{100}$ | 35.6 (17.6–62.2)$_{100}$ | 35.1 (19.1–52.5)$_{100}$ | 52.8 (28.7–80.5)$_{100}$ |
| Maximum absolute iHS | 50.4 (17.2–99.6)$_{100}$ | 97.2 (54.5–143.5)$_{100}$ | 142.1 (80.8–179.7)$_{100}$ | 142.1 (75.5–232.5)$_{100}$ |
| Maximum XP-EHH | 37.2 (15.0–83.5)$_{100}$ | 20.0 (7.0–47.8)$_{100}$ | 29.7 (14.0–62.5)$_{100}$ | 38.4 (17.5–85.0)$_{100}$ |

#### 4.3 Matching TMRCAs with Simulations

We compared selective sweeps and purifying selection to ask whether they could produce similar TMRCA profiles around *GRK2* and *TREM2*. We inferred TMRCA from the simulated genotypes using `gamma_smc_cu`, with the same population-specific $N_e$ values used in the full-chromosome analysis and a constant recombination rate of $10^{-8}$ per base per generation. For each locus, we compared the observed profile to the simulated one using 48 bins of 20 kb, which covered 960 kb. Denote the arithmetic mean of the inferred TMRCA in bin $b$ by $O_b$ and the simulated mean by $P_b$. We estimated the parameters of the simulation by minimizing

$$E = \left[ \frac{0.75}{|B|} \sum_{b \in B} \left( \log \frac{P_b}{O_b} \right)^2 + \frac{0.25}{|F|} \sum_{b \in F} \left( \log \frac{P_b}{O_b} \right)^2 \right]^{1/2},$$

where $B$ contains all 48 bins and $F$ contains the bins in the central 200 kb. This gives additional weight to the central region. The average TMRCA was first computed across the same 20 haplotype pairs, followed by averaging along the sequence within each bin. For the inferred TMRCA profiles, each site's estimated value represented the interval bounded by the midpoints to its neighbouring sites. These intervals were weighted by their overlap with each bin. For the true TMRCA values, the exact genomic interval of each simulated tree was used. The error was calculated from the TMRCA profile averaged across simulation runs. We simulated hard sweeps using `msprime` 1.4.4 and its `SweepGenicSelection` model [86]. At both loci, the selected allele started at frequency $1/(2N_e)$ and reached $1 - 1/(2N_e)$. A neutral period between the sweep endpoint and sampling represented the time elapsed since the sweep. Looking further back in time, ancestry before the sweep followed the standard coalescent. After an initial exploratory search, we tested nine parameter combinations using eight random seeds for each. The figure shows this final fitting stage. To test purifying selection, 20 sets of parameters were tested at each locus with the use of the same three random seeds per locus. They were made up of 18 models having mutations that are deleterious and two control models that were neutral. The best-fit purifying selection model was chosen from the 18 selected models, and the two control models were used for the comparison. The initial set of models had mutations within the regions occupied by the Ensembl Havana 104 exons and UCSC `phastConsElements100way` conserved elements. Subsequent models differed in population size, selection intensity, width and position of the selected interval, and the proportion of mutations that were deleterious. Neutral controls included the population size used for chromosome-wide inference, a smaller constant population size for *GRK2*, and a model with two periods of different population sizes for *TREM2*. We ran the purifying-selection simulations forward in time using `SLiM` 4.3 [85] through `stdpopsim` 0.3.0 [84]. Fitnesses were 1, $1+hs$ and $1+s$ for individuals with zero, one or two copies of a deleterious allele, respectively, with $s < 0$ and $h = 0.5$. Within each selected interval, a specified fraction of new mutations had the same deleterious effect, and the remainder were neutral. Simulations began with a burn-in of $10N_e$ physical generations. We used an approximate rescaling factor of $Q = 10$ to decrease the computational time needed. This divided the simulated census population size by ten, multiplied mutation rates and selection coefficients by ten, and adjusted recombination through `stdpopsim`. Times were then converted to physical generations. For the main purifying-selection analysis, TMRCA was estimated from the selected and neutral biallelic sites generated by the simulation, with each site counted once. As an extra check, neutral mutations were independently placed on the same purifying-selection genealogies, and TMRCA was estimated again. True TMRCA was also calculated directly from the simulated genealogies. For the sweep simulations, TMRCA was estimated from neutral mutations placed on the simulated genealogies. Differences in fit between the two model families therefore reflect both their genealogies and how variants were generated. After choosing the final parameters, we evaluated each model shown in the figure using 24 replicates. Shaded regions show the 10th–90th percentiles across these replicates.

#### 4.4 Geometric vs arithmetic per-gene aggregation

Table S12 compares the ranks under arithmetic and geometric gene aggregation. Each population is ranked over all finite gene scores separately for each summary.

Table S12: **Geometric vs arithmetic per-gene aggregation at reference loci.** For each gene we report the within-population rank percentile in the expected population under each statistic. LCT, MCM6, ADH1B, EDAR and KITLG have higher rank percentiles under arithmetic aggregation.

| Gene | Pop | Arithmetic mean | Geometric mean |
|---|---|---|---|
| LCT | CEU | 18.51% | 0.29% |
| MCM6 | CEU | 13.13% | 0.28% |
| EDAR | CHB | 4.79% | 0.46% |
| ADH1B | CHS | 15.90% | 2.18% |
| SLC24A5 | GBR | 0.04% | 0.06% |
| ABCC11 | CHB | 0.10% | 0.14% |
| KITLG | MXL | 6.30% | 0.63% |

Both summaries place SLC24A5 and ABCC11 below 0.2%. KITLG ranks at 6.30% under arithmetic aggregation and 0.63% under geometric aggregation.

#### 4.5 Sensitivity of candidate selection

The reference screen uses a 1% cutoff. Stage-2 cutoffs of 0.5%, 2% and 5% are applied uniformly with the remaining filters fixed. Separately, continental replication and clustering are varied one at a time. These sensitivity analyses are exploratory and were performed after inspecting the scan results. The reference threshold is fixed across the case studies.

Table S13: **Sensitivity under eight configurations.** The SD mask and 500 kb reference exclusion are fixed. Counts in stages 2–4 are genes; stage 5 counts distance-clustered loci.

| Configuration | Rank (%) | Cont. (%) | Mb | Stage 2 | Stage 3 | Stage 4 | Loci |
|---|---|---|---|---|---|---|---|
| Rank 0.5% | 0.5 | 5 | 1 | 226 | 215 | 193 | 58 |
| Reference screen | 1 | 5 | 1 | 620 | 579 | 540 | 143 |
| Rank 2% | 2 | 5 | 1 | 1289 | 1047 | 996 | 233 |
| Rank 5% | 5 | 5 | 1 | 2858 | 1246 | 1189 | 277 |
| Continental 1% | 1 | 1 | 1 | 620 | 173 | 160 | 52 |
| Continental 10% | 1 | 10 | 1 | 620 | 608 | 566 | 154 |
| Clustering 500 kb | 1 | 5 | 0.5 | 620 | 579 | 540 | 159 |
| Clustering 2 Mb | 1 | 5 | 2 | 620 | 579 | 540 | 124 |

Table S14: **All focal genes under the four uniform stage-2 cutoffs.** Each entry gives the number of populations below the cutoff, followed by screen status: R, representative; M, eligible stage-4 member represented by another gene; –, fails the multistage screen. The focal population comparison is fixed for each case, and both TREM genes are included. Counts alone do not establish eligibility because other filters also apply.

| Gene | Population set | 0.5% | 1% | 2% | 5% |
|---|---|---|---|---|---|
| *GRK2* | SAS+EUR | 7/10; R | 9/10; R | 10/10; R | 10/10; R |
| *TREML1* | non-AFR | 0/19; – | 0/19; – | 7/19; M | 18/19; M |
| *TREM2* | non-AFR | 0/19; – | 0/19; – | 10/19; R | 18/19; R |
| *IFIH1* | SAS+EUR | 2/10; R | 10/10; R | 10/10; R | 10/10; R |
| *CCDC92* | EAS | 0/5; – | 0/5; – | 1/5; M | 5/5; M |
| *CLEC6A* | EAS | 0/5; – | 0/5; – | 0/5; – | 4/5; – |
| *SLC6A15* | EAS | 0/5; – | 5/5; R | 5/5; R | 5/5; R |
| *BPIFA2* | SAS | 0/5; – | 0/5; – | 4/5; R | 5/5; R |

#### 4.6 Comparison of inference methods cited in this work

The table below compares the inference methods cited in this work with `gamma_smc_cu`.

| Method | Year | Sample requirement | Recommended use | Main limitation for this scan |
|---|---|---|---|---|
| `PSMC` [87] | 2011 | 1 diploid individual | Single-individual demographic-history reconstruction | Single-pair output, no population aggregation, poor power at recent times |
| `MSMC` / `MSMC2` [88, 89] | 2014/2020 | Phased haplotypes; `MSMC`: 2, 4 and 8 in primary analyses; `MSMC2`: pairwise composite | Population separation and recent $N_e$ estimation | `MSMC2` combines pairwise likelihoods, cost grows with pair count |
| `ASMC` [56] | 2018 | Pairwise; biobank-scale | Biobank-scale pairwise TMRCA via precomputed decoding quantities | Discretised TMRCA bins, higher per-pair CPU cost in the chromosome-22 benchmark |
| `gamma_smc` [90] | 2023 | Pairwise | Continuous TMRCA posterior across species (CPU) | CPU inference is slower in the chromosome-22 benchmark shown here |
| `gamma_smc_cu` (this work) | 2026 | Pairwise; cohort-scale | Cohort-scale pairwise TMRCA inference and gene-level summaries on CUDA GPUs | Requires a CUDA-capable GPU |
| `cxt` [55] | 2026 | Pair plus local frequency spectrum | Pairwise TMRCA and genome-wide demographic inference | All-pair cohort throughput not benchmarked here |
| `Relate` [10] | 2019 | Population haplotypes | ARG-based scan with selection on inferred branches | Pairwise summaries require extraction from the inferred genealogies |
| `tsinfer`+`tsdate` [91, 92] | 2019/2022 | Population haplotypes | Unified ARG inference + node dating across modern + ancient genomes | Requires high-quality phased genomes, ARG quality varies regionally |
| `ARG-Needle` [93] | 2023 | Biobank haplotypes | Threaded ARG for biobank scale; complex-trait analyses | Genome-wide pairwise output is implicit in the ARG, not directly indexed |
| iHS / nSL (`selscan`) [1, 94, 95] | 2006/2014 | Phased haplotypes | Ongoing selective sweeps from extended-haplotype homozygosity | Require suitable focal polymorphism, optional window summaries can dilute compact-gene signals |
| XP-EHH [3, 95] | 2007 | Two phased populations | Sweeps near or at fixation in one population | Depends on the comparison population and persistence of haplotype structure |
| Akbari *et al.* aDNA time-series [16] | 2026 | Dated ancient + modern samples | Directional selection within the geographic/temporal range of ancient samples | Limited to populations with dated ancient sampling (predominantly West Eurasia) |
| Field 2016 SDS [6] | 2016 | Modern whole-genome sequences | Very recent selection (∼2–3 kya in UK10K) | Requires reliable singleton calls, time horizon depends on sample size and demography |

Table S15: **Inference methods cited in this work, with their developmental milestones, recommended use cases, and drawbacks relative to the genome-wide pairwise-coalescence analysis reported here.** The table covers methods relevant to the analysis, rather than a comprehensive software comparison.

#### 4.7 Segmental-duplication mask source

In the unmasked ranking, 26 of the top 30 genes meet the duplication exclusion rule. After masking, the lowest-ranked genes include *SHCBP1*, *SLC24A5*, *MYEF2*, *CTXN2*, *VPS35* and *OR4C6*. The mask counts overlap each base once.

The UCSC `genomicSuperDups` track (GRCh38) used for SD masking (Methods) was downloaded from `https://hgdownload.soe.ucsc.edu/goldenPath/hg38/database/genomicSuperDups.txt.gz`.
